# Haplotype-aware pantranscriptome analyses using spliced pangenome graphs

**DOI:** 10.1101/2021.03.26.437240

**Authors:** Jonas A. Sibbesen, Jordan M. Eizenga, Adam M. Novak, Jouni Sirén, Xian Chang, Erik Garrison, Benedict Paten

## Abstract

Pangenomics is emerging as a powerful computational paradigm in bioinformatics. This field uses population-level genome reference structures, typically consisting of a sequence graph, to mitigate reference bias and facilitate analyses that were challenging with previous reference-based methods. In this work, we extend these methods into transcriptomics to analyze sequencing data using the pantranscriptome: a population-level transcriptomic reference. Our novel toolchain can construct spliced pangenome graphs, map RNA-seq data to these graphs, and perform haplotype-aware expression quantification of transcripts in a pantranscriptome. This workflow improves accuracy over state-of-the-art RNA-seq mapping methods, and it can efficiently quantify haplotype-specific transcript expression without needing to characterize a sample’s haplotypes beforehand.

## Introduction

Transcriptome profiling by RNA-seq has matured into a standard and essential tool for investigating cellular state. Bioinformatics workflows for processing RNA-seq data vary, but they generally begin by comparing sequencing reads to the sequence of a reference genome or reference transcriptome [1–4]. This is an expedient method that makes it practical to analyze the large volume of data produced by modern high-throughput sequencing.

Reference-based methods also have costs. When a sample’s genome differs from the reference, bioinformatics tools must account for the resulting mismatches between the sequencing data and the reference. This results in reduced ability to correctly identify reads with their transcript-of-origin, with larger genomic variation leading to a greater reduction in accuracy. This problem is known as reference bias [5].

Computational pangenomics has emerged as a powerful methodology for mitigating reference bias. Pangenomics approaches lean heavily on abundant, publicly-available data about common genomic variation for certain species (notably including humans). These methods incorporate population variation into the reference itself, usually in the form of a sequence graph [6, 7]. Mapping tools for pangenomic references have demonstrated reduced reference bias when mapping DNA reads [8–10]. This in turn facilitates downstream tasks that are frustrated by mapping biases, such as structural variant calling [11, 12].

The sequence graph formalism used in pangenomics has an additional attractive feature for RNA-seq data: it can represent splice junctions with little modification. Without this benefit, RNA-seq mappers for conventional references must make use of sometimes elaborate algorithmic heuristics to align over known splice junctions [2, 13]. Alternatively, they can map to only known isoforms, but this technique has difficulty estimating mapping uncertainty due to the re-use of exons across isoforms [14]. There is also evidence that population information can reduce reference bias problems that are particular to RNA-seq data. Accounting for population variation at splice-site motifs has been shown to aid in identifying novel splice sites [15].

The current methodological landscape in pangenomics is ripe to be extended to pantranscriptomics: using populations of reference transcriptomes to inform transcriptomic analyses. There is some precedent in a few existing transcriptomic methods that have used sequence graphs. AERON [16] uses splicing graphs and GraphAligner [17] to identify gene fusions. ASGAL [18] uses splicing graphs to identify novel splicing events. Finally, the pangenomic aligner HISAT2 [19] has its origins in the RNA-seq aligner HISAT [20], and it retains many of HISAT’s features for RNA-seq data. The performance of HISAT2 for pantranscriptomic mapping has not yet been characterized in published literature.

One transcriptomic analysis that is particularly prone to reference bias is allele-specific expression (ASE). ASE seeks to identify differences in gene expression between the two copies of a gene in a diploid organism. These differences are indicative of various biological processes, including cis-acting transcriptional regulation, nonsense-mediated decay, and genomic imprinting [21, 22]. The differences are identified by measuring the ratio between RNA-seq reads containing each allele of a heterozygous variant. However, the reads containing the non-reference allele are systematically less mappable because of reference bias, which can lead to both degraded and illusory signals of ASE [23]. Several approaches have been developed to deal with reference bias for ASE detection. Some methods filter out biased sites [24]. Others can mitigate bias at the read mapping stage, but require variant calls, often with phasing, for the individual being analyzed [25–27]. The variant information is either incorporated into the mapping algorithm to reduce reference bias or used to create a sample-specific diploid reference to map against. phASER phases called genotypes using read-backed and population-based phasing to produce estimates of haplotype-specific gene expression [28].

Pantranscriptomic approaches using existing haplotype panels for inferring haplotype-specific expressions have also been developed. AltHapAlignR maps reads to the linear reference genome and seven alternative HLA haplotypes to infer haplotype-specific transcript expression in the HLA region [29]. HLApers first aligns reads against all known HLA haplotypes to estimate the most likely haplotypes and then infers haplotype-specific gene expression [30]. However, both of these pantransciptomic approaches are limited to smaller genomic regions.

In this work, we present a novel bioinformatics toolchain for whole genome pantranscriptomic analysis, which consists of additions to the vg toolkit and a new standalone tool, rpvg. First, the vg rna tool can combine genomic variation data and transcript annotations to construct a spliced pangenome graph. Next, vg mpmap can align RNA-seq reads to these graphs with high accuracy. Finally, rpvg can use the alignments produced by vg mpmap to quantify haplotype-specific transcript expression. Moreover, the information about population variation that is embedded in the pantranscriptome reference makes it possible to do so without first characterizing the sample genome, and without restricting focus to SNVs.

## Results

### Haplotype-aware transcriptome analysis pipeline

In short, our pipeline works as follows (see Methods for a more detailed description). First, we construct a spliced pangenome graph using vg rna, a method developed as part of the vg toolkit [8]. vg rna adds splice junctions from a transcript annotation into a pangenome graph as edges and then labels the paths in the graph that correspond to transcripts (Figure 1a). Simultaneously, vg rna constructs a set of haplotype-specific transcripts (HSTs) from the transcript annotation and a set of known haplotypes by projecting the transcript paths onto each haplotype. vg rna uses the Graph Burrows-Wheeler Transform (GBWT) to efficiently store the HST paths, allowing the pipeline to scale to a pantranscriptome with millions of transcript paths [31]. Next, RNA-seq reads are mapped to the spliced pangenome graph using vg mpmap, a new splice-aware graph mapper in the vg toolkit that can align across both annotated and unannotated splice junctions (Figure 1b). vg mpmap produces multipath alignments that capture the local uncertainty of an alignment to different paths in the graph (Supplementary Figure 1). Lastly, the expression of the HSTs are inferred from the multipath alignments using rpvg (Figure 1c). rpvg uses a nested inference scheme that first infers the most probable underlying haplotype combinations (i.e. diplotypes) and then estimates the HST expression using expectation maximization conditioned on the haplotypes.

**Figure 1:**
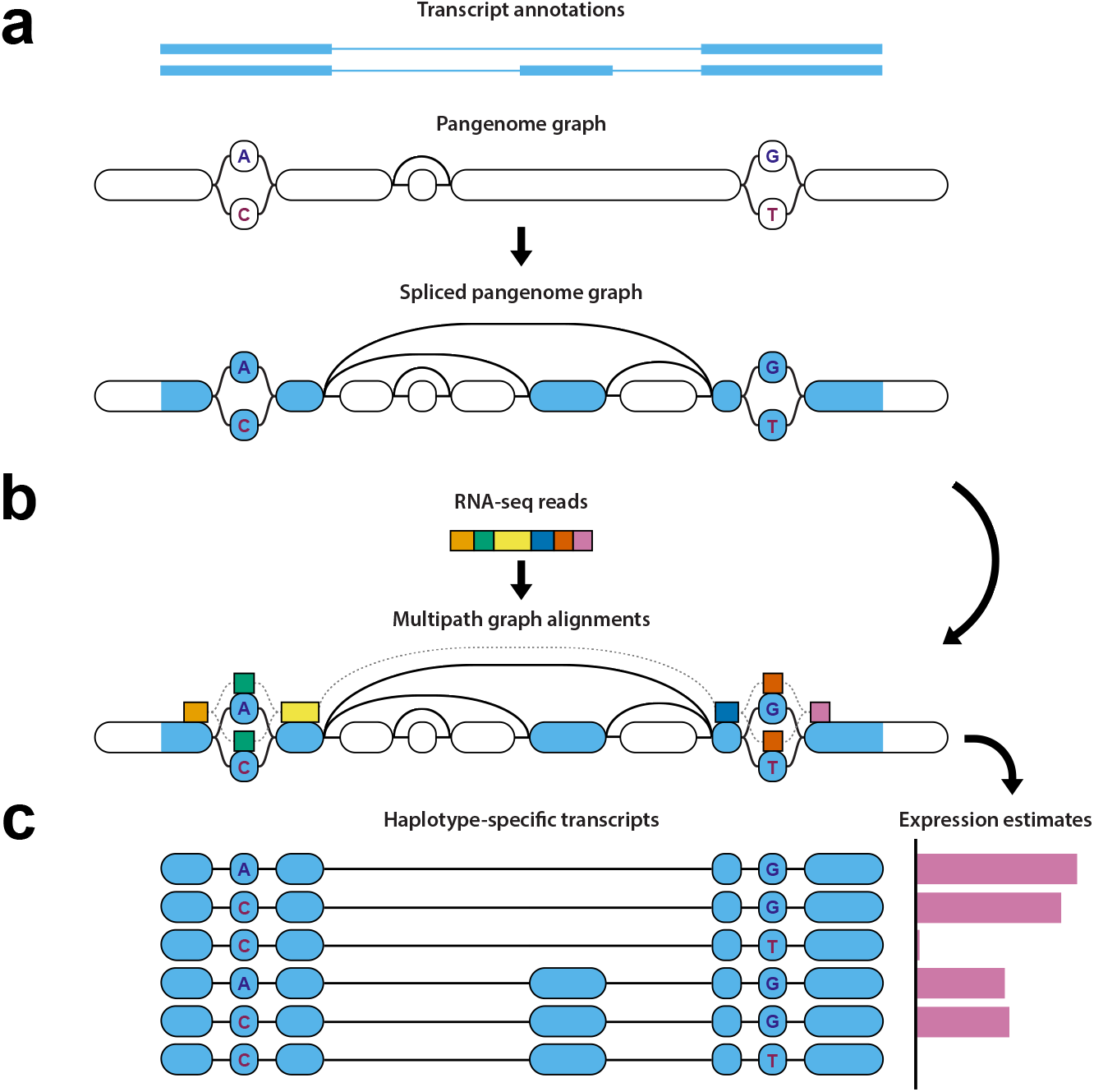
Diagram of haplotype-aware transcriptome analysis pipeline. The three major steps in the pipeline. **a** vg rna adds splice junctions derived from a transcript annotation to a pangenome graph to create a spliced pangenome graph. It simultaneously creates a pantranscriptome composed of a set of haplotype-specific transcripts (HSTs) using a panel of known haplotypes (not shown). **b** vg mpmap aligns RNA-seq reads to subgraphs of the spliced pangenome graph represented as a multipath alignment. **c** rpvg uses the alignments from mpmap to estimate the expression of the HSTs in the pantranscriptome.

### RNA-seq mapping benchmark

We compared vg mpmap against three other mappers: STAR [2], HISAT2 [19] and vg map [8]. STAR and HISAT2 can both use splicing information to guide mapping. However, of the two, only HISAT2 is able to also utilize genomic variants. vg map is not a splice-aware mapper, but it is still able to map to spliced pangenome graphs, which contain both splicing and genomic variation edges.

We used two different references for the comparison: the standard reference genome with added splice junctions (spliced reference) and a spliced pangenome graph containing both splice junctions and variants (spliced pangenome graph). For STAR, only the spliced reference was used. In addition, to assess alignment across unannotated splice junctions, we constructed references with a random 20% of transcripts removed before construction. This number was based on recent estimates of the fraction of novel transcripts in a sample using long reads [32]. For all of the tools besides STAR, this reference included variation (80% spliced graph), whereas STAR’s reference did not (80% spliced reference).

#### Simulated sequencing data

Paired-end reads were simulated from HSTs derived from the GENCODE transcript annotation set [33] and the NA12878 haplotypes from the 1000 Genomes Project (1000GP) [34]. vg sim was used to simulate the reads using reads from the ENCODE project (ENCSR000AED replicate 1) to parameterize the noise model [35, 36]. Fragment length distribution parameters used in the simulation were estimated from the same reads using RSEM [1]. The CEU population was excluded from the spliced pangenome graph as NA12878 is from that population, and we wanted to estimate performance for a new individual, who may not be as closely-related to the 1000GP populations. We simulated the HSTs with uniform expression across transcripts rather than trying to match a previous expression profile, which could bias expression towards easily-mappable transcripts.

Using the set of simulated reads we first compared the overall mapping performance of each method. Figure 2a shows the mapping error (1 – precision) and recall for different mapping quality thresholds. Reads are considered correctly mapped if one of their multi-alignments covers 90% of the true reference sequence alignment. The maximum number of reported multi-alignments per read was set to 10 for each method. The graph alignments from vg map and vg mpmap were projected to the reference sequence for this comparison. For vg mpmap, the group mapping quality value of the multi-alignments (see Methods) is used to stratify the results. The other tools are stratified by the mapping quality value of the alignment with the highest overlap, or the highest mapping quality if multiple alignments have the same best overlap. As can be seen in Figure 2a, vg mpmap achieves both a low error and high recall, while the other methods either had a high error or low recall. The same pattern is observed among primary alignments, ignoring multi-alignments (Supplementary Figure 2). The results also show that the spliced pangenome graph generally improves mapping performance. In addition, vg map and vg mpmap show substantially better calibration in their estimated mapping qualities, especially among the most confidently mapped reads (Supplementary Figure 3).

**Figure 2:**
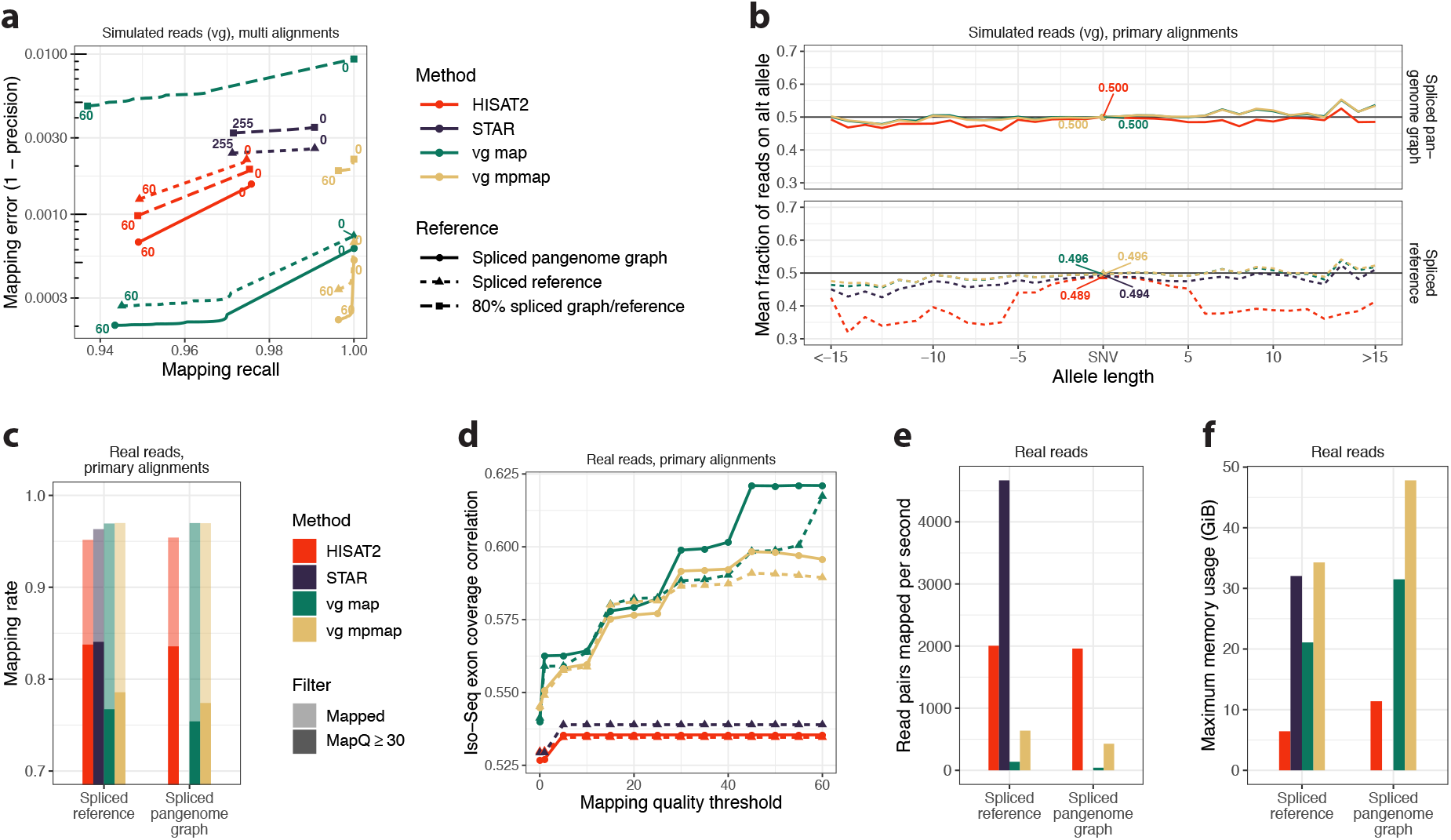
Mapping benchmark using RNA-seq data from NA12878. RNA-seq mapping results comparing vg mpmap and three other methods using simulated and real Illumina data (“vg sim (ENC, uniform)”, *N* = 50, 000, 000 read pairs, and “ENCSR000AED”, *N* = 97, 548, 052 read pairs, in Supplementary Table 5 and 4, respectively). Solid lines show results for spliced pangenome graph (“1000GP (all, excl. CEU)” in Supplementary Table 2), short dashed lines for spliced reference, and long dashed lines for a spliced reference (STAR) or spliced pangenome graph (all others) made with a random subset containing 80% of the transcripts. **a** Mapping error and recall for different mapping quality thresholds (colored numbers) using simulated data. Reads are considered correctly mapped if one of their multi-alignments covers 90% of the true reference sequence alignment. **b** Mean fraction of mapped reads supporting the non-reference allele for variants of different lengths in simulated data. Calculated from primary alignments with a mapping quality value of at least 30. Negative lengths correspond to deletions and positive to insertions. The colored numbers are the mean fraction for SNVs. **c** Mapping rate using real data. The solid bars show the mapping rate using a mapping quality threshold of 30. **d** Pearson correlation between Illumina and Iso-Seq exon coverage using real data as a function of mapping quality threshold. Exons were defined by the Iso-Seq alignments and each exon’s coverage was normalized by its length. **e** Number of read pairs mapped per second per thread using real data. The mapping times were estimated using 16 threads on a AWS m5.4xlarge instance. **f** Maximum memory usage for mapping in gigabytes using real data.

On the 80% spliced references, all of the tools’ performance decreases relative to the corresponding reference constructed with the full transcript set. As expected, vg map’s performance decreases dramatically, since it can only align over splice junctions represented in the graph. vg mpmap’s reduction in performance is larger than STAR and HISAT2’s. This reduction is concentrated on reads containing unannotated splice junctions (Supplementary Figure 4), but vg mpmap’s performance is still competitive with both of the other tools in the aggregate read set.

Next, we looked at whether using a variant-aware approach reduces reference bias. Figure 2b shows the mean fraction of reads mapped to the alternative allele for different allele lengths. Negative values correspond to deletions and positive values to insertions. When using the spliced reference genome, all methods exhibit a bias towards the reference allele, with vg map and mpmap showing less bias than the other methods. Using the spliced pangenome graph results in substantially reduced bias for all methods. We also analyzed the mapping error and recall on reads stratified by the number of variants they contain (Supplementary Figure 5). This analysis corroborates the allele bias results; vg mpmap and vg map retain high recall in the presence of variants, whereas HISAT2 and STAR’s performance decreases substantially, especially in the presence of indels.

Previous research has pointed out that allelic bias can also result from differential uniqueness between two alleles [24]. The WASP tool combats this bias by filtering out some reads. We compared allelic bias between the four mapping tools and a pipeline consisting of WASP and STAR. Using simulated data with no allelic bias, we measured 1) the proportion of variant sites with a statistically significant allele skew (two-sided binomial test) and 2) the number of sites with coverage at least 20 (Supplementary Figure 6). Both HISAT2 and STAR show an increase in falsely significant tests above the nominal false positive rate of *α* = 0.01, especially for insertions and deletions. The WASP (STAR) pipeline, vg map, and vg mpmap all show approximately the expected rate of false positives for all variant types. In addition, compared to the WASP (STAR) pipeline, vg mpmap retains 5,670 more variant sites with high coverage (≥ 20 reads) when using the spliced personal graph.

The mapping results were corroborated by alternative evaluation methodologies. First, we used an alternate correctness criterion based on aligning within 100 bases of the correct position on the paths in the graph (Supplementary Figure 7), which gave qualitatively similar results. Second, we used RSEM as an alternative read simulator (Supplementary Figure 8). All mapping tools showed similar performance with the alternative simulator except HISAT2, which had relatively higher recall.

The set of simulated reads used for the mapping evaluation presented in Figure 2a,b was not used to optimize the algorithmic design or parameters of vg map and vg mpmap. Thus, these reads can be considered a “test set”. Supplementary Figure 9a-c shows the results on the “training set” that was used for optimizing the method, which used RNA-seq data from Tilgner, et al. [37] to parameterize the read simulation. The mapping error and recall estimates for vg mpmap are quite similar between the two data sets, whereas the performance of all the other methods was generally worse on the training data. Simulated data from RSEM was also used during the development of vg mpmap.

#### Real sequencing data

We used the same read set from the ENCODE project that was used to parametrize the simulations to benchmark the methods on real data. We first looked at the fraction of aligned reads for each method (Figure 2c). As can been seen in the figure, all methods have comparable overall mapping rates. When only looking at alignments with a mapping quality value of at least 30, both STAR and HISAT2 show noticeably higher rates compared to vg map and vg mpmap. However, it seems that the cost of these higher mapping rates is lower precision (Figure 2a) and poorly estimated mapping qualities (Supplementary Figure 3). This most likely results from vg mpmap and vg map finding high-scoring secondary alignments that the other mapping tools did not find.

Ground-truth alignments are not available for real data, so we use a proxy based on Pacific Biosciences (PacBio) Iso-Seq read mappings instead. Specifically, we compare to Iso-Seq read alignments generated by the ENCODE project (ENCSR706ANY) from the same cell line as the Illumina reads [35, 36]. Since the cell line is the same, we expect the transcript expression to be similar despite some technical biases due to the different sequencing protocols. Moreover, long reads can generally be mapped more confidently than short reads. Thus, higher correlation in coverage between short read mappings and the Iso-Seq mappings should be indicative of more accurate short read mappings in the aggregate. Figure 2d shows the estimated Pearson correlation in the coverage of each exon as a function of mapping quality threshold. As can be seen, both vg map and mpmap achieves higher correlation than STAR and HISAT2, with the spliced pangenome graph resulting in even higher correlation for both (see Supplementary Figure 10 for the full scatter plot). The graph alignments from vg map and mpmap were projected to the reference genome for this analysis.

Finally, we compared the methods’ mapping speed and memory usage. Figure 2e shows the number of read pairs mapped per second per thread. Conversion from SAM to BAM was included in the HISAT2 time estimate to be more comparable to the output type of the other methods. vg mpmap’s increase in accuracy does not come for free; it is between 3.1 and 4.6 times slower than HISAT2, depending on the graph. However, it is 10.3 times faster than vg map on the spliced pangenome graph. vg mpmap uses slightly more memory than STAR on the spliced reference and somewhat more memory than HISAT2 on the spliced pangenome graph (Figure 2f).

We also compared the mapping performance of the different methods on the real RNA-seq data from Tilgner, et al. [37] and CHM13 RNA-seq data from the T2T consortium [38] (Supplementary Figure 9d,e and 11). Similar overall patterns are observed using these data sets. It is important to mention that the CHM13 data was used during the development of vg mpmap, and the other set was used to optimize the parameters of vg map and vg mpmap.

### Haplotype-specific transcript quantification

We compared rpvg to three other transcript quantification methods: Kallisto [3], Salmon [4], and RSEM [1]. We stress that none of these methods were developed to work on pantranscriptomes with millions of HSTs. However, they serve as a point of reference for what accuracy is achievable without new methods development. rpvg’s inference model includes both a diplotype and HST expression, conditioned on the diplotype. However, to facilitate the comparison to other tools, we report here the marginal expression over all HSTs, which is directly comparable to the output of the other tools that lack a diplotype model.

The simulated data was generated by vg sim, largely as described for the mapping benchmark. The only difference was that, rather than simulating transcripts with uniform expression, we simulated according to an expression profile that was estimated by RSEM using the same ENCODE reads. Three different pantranscriptomes were generated for the evaluation using different sets of 1000GP haplotypes (Supplementary Table 3): 1) all European haplotypes excluding the CEU population (“Europe (excl. CEU)”, 2,515,408 HSTs) 2) all haplotypes excluding the CEU population (“Whole (excl. CEU)”, 11,626,948 HSTs) and 3) all haplotypes (“Whole”, 11,835,580 HSTs). The CEU population was excluded for the same reason as in the mapping benchmark: NA12878 is part of this population, and we wanted to imitate the realistic setting in which a sample is not as well represented by the haplotype panel. In addition, we created a personal sample-specific transcriptome consisting of NA12878 HSTs (“Personal (NA12878)”, 235,400 HSTs). This transcriptome corresponds to the ideal case where a sample’s haplotypes are known beforehand. For the real data, 104 mitochondrial and scaffold transcripts were added to the pantranscriptomes. For all pantranscriptomes, rpvg uses a Graph Burrows-Wheeler transform (GBWT) to store the HSTs [31]. For the other methods the sequences of the HSTs in FASTA format were provided as input. HSTs with a haplotype probability below 0.8 were filtered from the rpvg output (Supplementary Figure 12).

We first looked at the method’s ability to accurately predict whether an HST was expressed or not. Figure 3a shows the recall and precision using simulated data. The results were stratified by different expression thresholds up to a value of 10 TPM (transcripts per million). Note that we were not able to run RSEM on the two largest pan-transcriptomes used in the analysis. rpvg exhibits a much higher precision than the other tools for all pantranscriptomes. 96.9% of the HSTs with an expression value of at least 1 TPM are correctly predicted to be expressed by rpvg using the “Whole (excl. CEU)” pantranscriptome. This illustrates the importance of having a diplotyping model when inferring HST expression using a pantranscriptome reference, which is one of the major differences between rpvg and the other methods. Importantly, only a minor difference is observed between the pantranscriptomes without the CEU population (excl. CEU) and the whole pantranscriptome (“Whole”), which contains NA12878. This could be explained by the fact that less than 2% of HSTs are on average unique to a specific sample when compared to all samples in other populations using the 1000GP data (Supplementary Figure 13). This suggests that haplotype panels like the 1000GP are a good alternative when a sample’s haplotypes are not available, although there are always limits to panel diversity, and some samples will be less well-represented by a 1000GP pantranscriptome.

**Figure 3:**
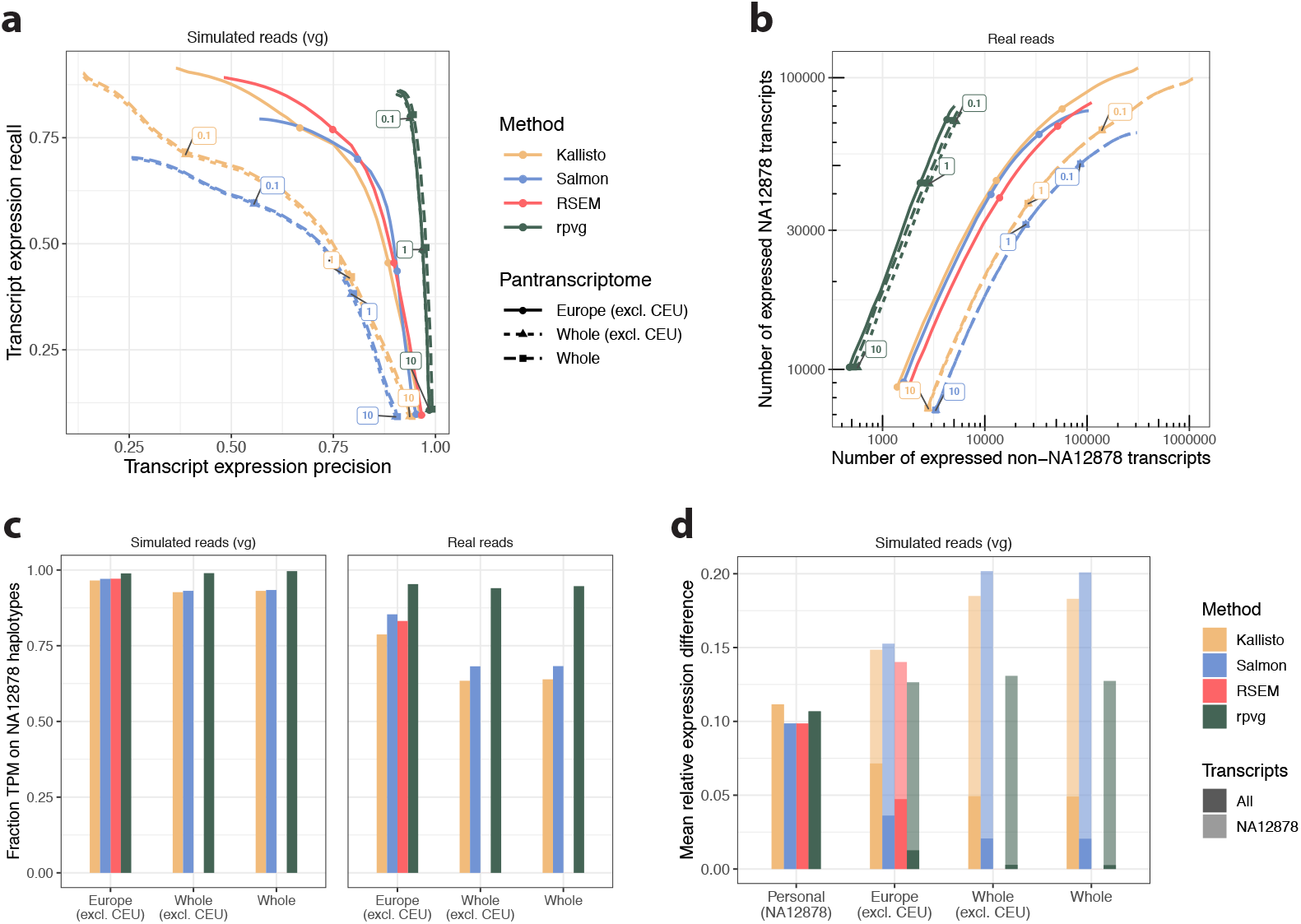
Haplotype-specific transcript quantification benchmark using RNA-seq data from NA12878. Haplotype-specific transcript (HST) quantification results comparing rpvg against three other methods using simulated and real Illumina data (“vg sim (ENC, RSEM)”, *N* = 50, 000, 000 read pairs, and “ENCSR000AED”, *N* = 97, 548, 052 read pairs, in Supplementary Table 5 and 4, respectively). The other methods were not designed for large pantranscriptomes, but they serve as a point of comparison for the performance that could be achieved without new methods development. Solid lines with circles are results using a pantranscriptome generated from 1000 Genomes Project (1000GP) European haplotypes excluding the CEU population. Dashed lines with triangles and squares are results using a pantranscriptome generated from all 1000GP haplotypes without and with the CEU population, respectively (Supplementary Table 3). **a** Recall and precision of whether a transcript is correctly assigned nonzero expression for different expression value thresholds (colored numbers for “Whole (excl. CEU)” pantranscriptome) using simulated data. Expression is measured in transcripts per million (TPM). **b** Number of expressed transcripts from NA12878 haplotypes shown against the number from non-NA12878 haplotypes for different expression value thresholds (colored numbers) using real data. **c** Fraction of transcript expression (in TPM) assigned to NA12878 haplotypes for different pantranscriptomes using simulated (left) and real (right) data. **d** Mean absolute relative expression difference (MARD) between simulated and estimated expression (in TPM) for different pantranscriptomes using simulated data. MARD was calculated using either all HSTs in the pantranscriptome (solid bars) or using only the NA12878 HSTs (shaded bars). “Personal (NA12878)” is a personal sample-specific transcriptome generated from 1000GP NA12878 haplotypes (Supplementary Table 3).

We also evaluated the accuracy of the HST expression estimation using real sequencing data (Figure 3b). Since we do not know which transcripts are expressed in real data, we focus instead on the haplotype estimation. Sample NA12878’s haplotypes are known to a reasonably high degree of certainty. Thus, we can indirectly measure accuracy by asking whether the HSTs that are estimated to be expressed are in fact from NA12878. Figure 3b shows that rpvg predicts markedly fewer HSTs from non-NA12878 haplotypes than the other methods. Also, we see again only a minor difference between pantranscriptomes. Next, we compared the fraction of transcript expression (in TPM) that was attributed to NA12878 haplotypes. This is shown in Figure 3c for both simulated (left bars) and real (right bars) data. We see that rpvg attributes more than 98.8% and 94.0% of the expression to NA12878 haplotypes when using simulated and real data, respectively. Furthermore, the figure shows that rpvg’s prediction accuracy only decreases slightly when the size of the pantranscriptome increases from 2.5M HSTs in “Europe (excl. CEU)” to 11.6M in “Whole (excl. CEU)”.

We compared how well the different methods could predict the correct expression value. Figure 3d shows the mean absolute relative expression difference (MARD) between the expression values of the simulated reads and the estimated values. The solid bars are MARD values when using all HSTs in the pantranscriptomes, and the shaded bars are when comparing the NA12878 HSTs only. Note that these bars are the same for the personal sample-specific set, which consists of only NA12878’s HSTs. On the personal set, rpvg performs comparably to the other methods. However, as the size of the pantranscriptome grows, the MARD on the NA12878 transcript set only increases slightly for rpvg. Supplementary Figure 14 shows the full scatter plots of the simulated and estimated expression values for the NA12878 HSTs. The much lower values for all methods when using all HSTs can be explained by the large number of unexpressed HSTs in the full pantranscriptomes. We also compared the expression values using Spearman correlation (Supplementary Figure 15). This metric supported overall similar conclusions, albeit with Kallisto and RSEM performing comparably to rpvg when using the pantranscriptomes but restricting focus to NA12878’s haplotypes. This suggests that Kallisto and RSEM accurately rank these transcripts’ expression but do not accurately estimate the absolute quantity.

Using only the HSTs estimated to be expressed by each method we see similar results for MARD and Spearman correlation (Supplementary Figure 16). However, when looking at reference transcript-level expression estimates, the other methods exhibit overall better MARDs (Supplementary Figure 17). Reference transcript expression was calculated by adding together the estimated expression values of all HSTs from the same transcript without for rpvg filtering HSTs with a low haplotype probability.

Similarly to the mapping benchmark, we also evaluated rpvg on NA12878 RNA-seq data from Tilgner et al. [37] and CHM13 data from the T2T consortium [38] (Supplementary Figure 18 and 19). Overall, similar conclusions can be drawn using these data. It is important to mention that both data sets were used to optimize the parameters of rpvg.

To show the advantage of the multipath alignment format when inferring HST expression we repeated the evaluation using single-path alignments from both vg map and vg mpmap as input to rpvg (Supplementary Figure 20). The vg mpmap single-path alignments were created by finding the best scoring path in each multipath alignment. For all pantranscriptomes and data sets, rpvg gave the best results using the multipath alignments.

rpvg can optionally run a Gibbs sampling step to quantify the uncertainty in the expression estimates. To evaluate the HST expression values estimated by Gibbs sampling we calculated the 90% credibility intervals from a 1000 samples and compared them to the simulated HST expression values (Supplementary Figure 21). 86.4% of the intervals contained the simulated expression value, which is close to the expected proportion.

To gauge the vg mpmap-rpvg pipeline’s ability to estimate allele-specific expression (ASE), we converted the simulated HST expression values and rpvg estimates to allele-specific read counts. In addition, we inferred allele-specific read counts using WASP [24] with STAR [2] alignments. Using both simulated and inferred allele-specific read counts, we identified heterozygotic variants that showed significant ASE using a two-sided binomial test with p-values adjusted using the Benjamini-Hochberg procedure and a False Discovery Rate (FDR) *α* = 0.1. Note, we only used WASP for bias correction and allele-specific read counting, and not its downstream inference method. Supplementary Figure 22 shows the results of comparing the significant variants between the simulation and methods. The vg mpmap-rpvg pipeline achieves markedly higher true positive rate with the same false positive rate as the WASP (STAR) pipeline. Moreover, similar rates were observed for insertions and deletion when using the vg mpmap-rpvg pipeline, whereas these variants were excluded by the WASP (STAR) pipeline.

To assess the vg mpmap-rpvg pipeline’s robustness to samples from individuals with admixed ancestry, we applied it to two samples from a recent study [39]: one of European American ancestry and one of African American ancestry (Supplementary Figure 23). We expect that the African American individual has a more admixed ancestry due to the higher genomic diversity present in Africa and the United States’ history of widespread slave rape by slave owners of European ancestry [40]. As a proxy for accuracy, we quantify how frequently rpvg can confidently identify two or fewer HSTs for a transcript. If none of the HSTs in the pantranscriptome match the individual, the posterior over HSTs will tend to diffuse onto multiple similar HSTs. Consistent with expectations from Supplementary Figure 13, we see somewhat lower accuracy on the African American individual’s sample, but the difference is small.

### Evaluating HLA typing

To evaluate the vg mpmap-rpvg pipeline’s ability to infer diplotypes for genes in the highly-polymorphic human leukocyte antigen (HLA) region, we created two HLA-specific pantranscriptomes using alleles from the IPD-IMGT/HLA database [41] (see Supplementary Table 3). We ran the pipeline on RNA-seq data for three trios from the 1000GP sequenced as part of the Human Genome Structural Variation Consortium (HGSVC) [42] (see Supplementary Table 4). Figure 4a shows the number of predicted expressed transcripts for each child and whether the inferred transcript diplotype is concordant with the inferred diplotype of the parents, according to Mendelian inheritance. Also shown is the fraction of expression assigned to concordant transcripts. For most of the genes, almost all of the expression is assigned to the concordant transcript, with the exception of B and DRB5. DRB5 is classified as a paralog which might explain the bad performance for this gene specifically. However, for B it is currently unclear why the typing performance is not as good as the other genes.

**Figure 4:**
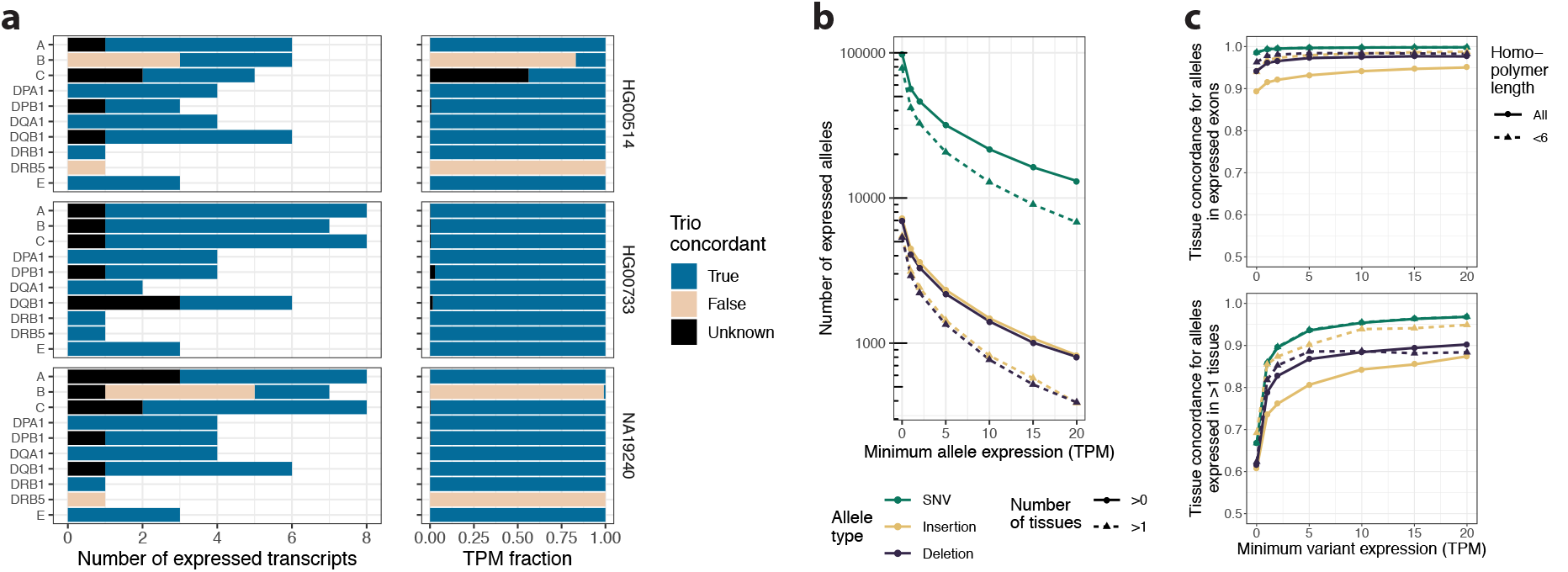
HLA typing and allele concordance evaluation using RNA-seq data from trios and different tissues. **a** HLA typing results from the vg mpmap-rpvg pipeline using real Illumina data from 3 trios (see Supplementary Table 4). The evaluation used the “HLA (10)” pantranscriptome containing ten HLA genes all of which had at least 100 haplotypes (Supplementary Table 3). The plot on the left shows the number of transcripts predicted to be expressed for each child and gene, and the right shows the fraction of trio concordant expression. Expression is measured in transcripts per million (TPM). The colors represent whether the predicted diplotype for the child is concordant with the prediction in the parents according to Mendelian inheritance. The concordance is labeled unknown when a transcript is not expressed in one of the parents. **b** & **c** Variant genotyping analyses using the vg mpmap-rpvg pipeline on real Illumina data from 5 tissues from the same individual (see Supplementary Table 4). The analyses used the pantranscriptome generated from all 1000 Genomes Project (1000GP) haplotypes (“Whole” in Supplementary Table 3). Colors represent different allele types. **b** Number of variant alleles predicted to be expressed in at least one (solid lines) or two tissues (dashed lines) for different expression (in TPM) thresholds. **c** Fraction of alleles predicted to be concordant across tissues for alleles in all expressed exons (including unexpressed alleles of expressed variants; top) and alleles expressed in at least two tissues (bottom). An allele is defined as concordant if it is either expressed or not expressed in all tissues that have the corresponding variant expressed. The results are shown for different variant expression (sum of expression across all alleles in TPM) thresholds and homopolymer lengths (solid and dashed lines).

We also ran the pipeline on ten randomly selected CEU samples from Geuvadis [43] for which HLA typing results are available from other studies of genomic sequencing data [44, 45]. Using this data gave similar results compared to the trio data, with typing of A, DQB1 and DRB1 being correct in all samples and typing of B being incorrect in some of the samples (Supplementary Figure 24). In addition, typing of C was also incorrect in some of the samples for this data.

While the presented results look promising, other HLA typing methods have shown similar or somewhat higher accuracy dependent on the gene, but the small sample size used here makes it challenging to determine the exact difference in performance [46]. However, one of the major advantages of the vg mpmap-rpvg pipeline compared to these methods is that it also provides HST expression estimates in addition to the typing.

### Investigating variant genotyping and effect prediction

To illustrate the vg mpmap-rpvg pipeline’s ability to genotype variants in a pantranscriptome from RNA-seq data, we ran the pipeline on five different tissues. All tissues were from the same individual and downloaded from the ENCODE project [35, 36] (see Supplementary Table 4). Figure 4b shows the number of expressed variant alleles for different expression thresholds. As expected, markedly more SNVs are predicted to be expressed than indels. Among indels, a similar number of insertions and deletions are predicted to be expressed.

Next, we looked at whether the expression of alleles were concordant across tissues. An allele was defined as concordant if it was either expressed or not expressed in all tissues for which the corresponding variant is expressed (see Supplementary Figure 25). To account for allelic dropout for lowly expressed exons, we calculated the concordance for different thresholds of variant expression (sum of expression across all alleles). Figure 4c shows the results of this analysis for alleles in all expressed exons (including unexpressed alleles of expressed variants; top) and alleles expressed in at least two tissues (bottom). When looking at alleles in all expressed exons, we see that the concordance rate reaches 0.95 for insertion, with higher values for deletions and SNVs. For alleles expressed in at least two tissues the rates are lower, but still over 0.95 for SNVs and 0.85 for indels. Long homopolymers are known to increase error rates in Illumina sequencing, which we have observed to be problematic for the vg mpmap-rpvg in some cases. To illustrate this, we removed all variants in homopolymers longer than five and repeated the concordance analysis. Using this set, the performance for insertions did indeed improve, whereas performance for deletions were surprisingly largely unchanged.

Finally, we investigated the effect of the predicted variants on functional elements. This was done using the Ensembl Variant Effect Predictor (VEP) toolset [47]. The results for expressed variant alleles in exons with a TPM of at least five are shown in Supplementary Figure 26. The number of predicted protein truncating variants (frameshift, splice donor, splice acceptor and stop gain) is comparable to or lower than what have previously been described dependent on the study [48, 49]. A lower number is not surprising since unexpressed variants are missed when using RNA-seq. The number of homozygotic protein truncating variants were higher than expected. Most of them were predicted to be expressed in more than one tissue.

### Assaying isoform-specific genomic imprinting

To demonstrate the utility of the vg mpmap-rpvg pipeline on a biological problem, we performed an exploratory analysis of genomic imprinting in a human sample. Genomic imprinting is the phenomenon in mammals in which some genes are expressed only from the copy inherited from a specific parent. That is, either the maternal or paternal copy is silenced, regardless of the genomic sequence of that haplotype. This is accomplished through mitotically heritable epigenetic modifications that are established during early development [50].

Several previous studies have studied imprinting genome-wide by quantifying ASE in RNA-seq data. These studies have demonstrated that imprinting varies substantially across tissues [51, 52] and varies in intensity across genes, with many genes showing biased expression away from one parent-of-origin copy but not complete silencing [21, 52, 53]. In addition, a handful of genes have been identified in which the polarity of imprinting depends on the isoform: some isoforms of the same gene are biased toward the paternal copy and others toward the maternal copy [21].

The previous genome-wide studies have methodological limitations that diminish their ability to detect isoform-level imprinting. Some have aggregated ASE across all isoforms of the gene, which precludes isoform-level analysis a priori [51–53]. The largest study, by Zink, et al. [21], performed all tests of imprinting on individual SNVs. This method can sometimes detect isoform-level differences if the isoforms have some unshared exons. However, in shared exons, the ASE signal from the highest-expressed isoforms can drown out the signal of lower-expressed isoforms. Depending on the configuration of exons in the isoforms, this can make it very challenging to identify imprinting of opposite polarity within the same gene.

Figure 5 shows results from our exploratory demonstration of isoform-level imprinting analysis using vg mpmap and rpvg. We ran the entire pipeline using RNA-seq data from a lymphoblastoid cell line derived from 1000GP sample NA12878, which was sequenced as part of the ENCODE project [35]. As a confirmatory analysis, we first looked for ASE in genes that have previously been identified as imprinted. In particular, we looked at the 20 genes with the most significant *p*-values from Zink’s, et al. study [21]. To mirror the methods used in this paper, we derived variant-specific ASE values by aggregating the expression across all HSTs that contain a given allele for a variant. Figure 5a shows that the vg mpmap-rpvg pipeline detects ASE at heterozygous variants in these imprinted genes at a markedly higher rate than in background across all genes.

**Figure 5:**
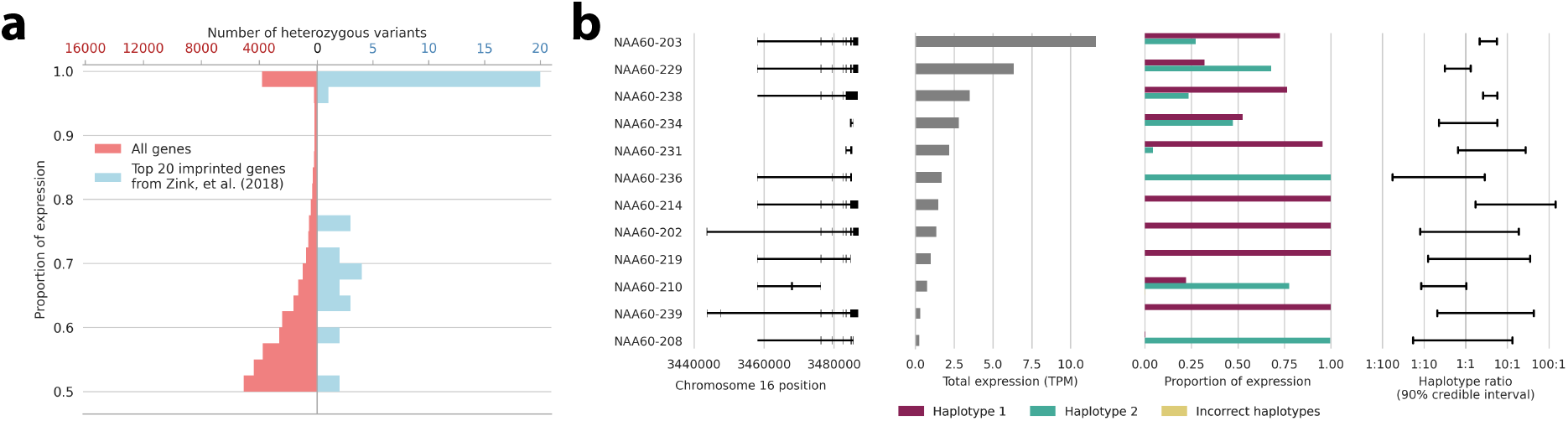
Exploratory demonstration of analyzing genomic imprinting using data from NA12878 lymphoblastoid cell line. Results of the vg mpmap-rpvg pipeline on RNA-seq data from a lymphoblastoid cell line from the ENCODE Project, focusing on genes previously identified as imprinted in blood (“ENCSR000AED” in Supplementary Table 4). The analyses used the pantranscriptome generated from all 1000GP haplotypes except haplotypes from the CEU population (“Whole (excl. CEU)” in Supplementary Table 3) **a** The proportion of expression attributed to the higher-expressed allele of heterozygous variants among the 20 most significantly imprinted genes from Zink’s, et al. study [21] compared to all genes. The axes are scaled so that both histograms have the same area. **b** Isoform-level haplotype-specific expression in NAA60, which was identified as imprinted but not as having isoform-dependent reversals in the polarity of imprinting. Isoforms with expression less than 0.25 transcripts per million (TPM) are not shown.

The vg mpmap-rpvg pipeline is also capable of detecting isoform-dependent genomic imprinting. Figure 5b shows an illustrative example in the gene NAA60, which codes for an enzyme that acetylates the N-terminus of proteins in the Golgi apparatus. The isoforms show a complex pattern of imprinting polarity, which does not correlate strongly with exons or start sites in any particular genomic region. Given the large differences in expression of these isoforms, the SNV-based analysis would have had difficulty identifying imprinting in the more lowly-expressed isoforms, and indeed this gene was reported as imprinted (in fact, it is among the 20 most significantly genes referred to above) but not as having isoform-dependent imprinting [21]. However, this gene has been identified as having isoform-dependent imprinting using RT-PCR in patients with uniparental disomies [54]. Nevertheless, it should be emphasized that this exploratory analysis of a single sample, while suggestive, is insufficient to conclusively demonstrate isoform-dependent imprinting in NAA60. Doing so would require further biological replicates and more rigorous controls for cis-regulation and cell line clonality [53].

## Discussion

The pace of development in the field of eukaryotic pangenomics has surged in recent years. Improvements in sequencing technology have made it practical to characterize the genomes of increasingly many samples. As a result, pangenomes made from tens to hundreds of reference-quality genome assemblies have been constructed for several agricultural organisms [55–57], and similar efforts are underway for humans by the Human Pangenome Reference Consortium and others [58]. Simultaneously, the bioinformatics tools to do pangenomic analyses have matured to the point of practicality for many applications [11, 59, 60]. Moving forward, we anticipate that pangenomic methods will continue to expand to inform increasingly many areas of genomics [61].

In this work, we have presented one step in this expansion: generalizing transcriptomics into pantranscriptomics. Our novel bioinformatics pipeline provides a full stack of tools for pantranscriptomic analysis. It can construct pantranscriptomes, map RNA-seq reads to these pantranscriptomes, and quantify transcription with haplotype-resolution. The construction takes advantage of efficient pangenome data structures, the mapping achieves a desirable balance of accuracy and speed, and the quantification can infer haplotype-specific transcript expression even when the sample’s haplotypes are not known beforehand.

Some downstream applications are already apparent. For one, the pipeline can be used to study causes of haplotype-specific differential expression. We demonstrated its capabilities on one such example: genomic imprinting. The demonstration showed suggestive evidence of complex patterns of imprinting at the isoform level, which would have been difficult or impossible to detect with previous genomewide methodologies. The pipeline could be similarly used to study other sources of haplotype-specific differential expression, such as nonsense-mediated decay and cis-regulation.

Another application is characterizing genotypes and haplotypes in coding regions from RNA-seq data. We demonstrated this capability by calling genotypes and HLA diplotypes. However, work is still needed to improve computational efficiency and accuracy in the HLA region. One of the major complications is that the dense variation in this region produces complicated graph topologies that lead to significant uncertainty in local alignments.

For all of these applications, the vg mpmap-rpvg pipeline increases the information that is available from RNA-seq data without paired genomic sequencing. This will enable low-cost study designs and deeper reanalyses of existing data.

Of course, our pipeline also has limitations. We have developed it to have good performance on pantranscriptomes constructed from phased variant calls. This is presently the most available data resource for constructing pangenomes. However, as increasingly many haplotype-resolved assemblies are produced, we predict that the emphasis in pangenomics will shift to pangenome graphs constructed from whole genome alignments. Constructing these graphs is currently an area of active research [62, 63]. Such graphs have more complicated topologies, often involving complex cyclic motifs. Experience leads us to believe that pantranscriptomic tools will require further methods development to use these data resources effectively.

Additional work on downstream analyses will be necessary to fully utilize haplotype-specific transcript expression inference. For example, current differential expression methods rely on being able to align transcript counts between transcripts of different individuals [64]. This is usually trivial at the transcript level, but it is difficult at the HST level, since different individuals may not share a haplotype. While HST expression estimates can always be marginalized to produce allele or transcript expression estimates, more general statistical frameworks will need to be developed to avoid information loss between these steps in transcriptomic pipelines.

Our pipeline is optimized for short-read RNA-seq data. The higher-error long-read RNA-seq technologies developed more recently require specifically-tailored algorithms for efficient analysis [32, 65]. Pantranscriptomic analyses of long-read RNA-seq data will likewise require further development, although the pipeline described here could serve as a platform for this development. Nevertheless, the cost-effectiveness of short-read sequencing virtually ensures that it will remain an important part of the sequencing landscape into the near future.

Finally, our pipeline also relies on having a comprehensive pantranscriptome that contains many of the sample’s haplotype-specific transcripts. The pantranscriptomes used in this study (based on the 1000 Genome Project) provided good results in the three samples analyzed, but this performance may not extend to samples from other populations. Here—and throughout pangenomics—there is a compelling case to improve the completeness of data resources through more diverse sampling.

## Acknowledgements

Research reported in this publication was supported by the National Human Genome Research Institute of the National Institutes of Health under Award Numbers U01HG010961, R01HG010485, U41HG010972, U24HG011853 and OT2OD026682. The content is solely the responsibility of the authors and does not necessarily represent the official views of the National Institutes of Health. The work of JAS was supported by the Carlsberg Foundation. We thank the ENCODE Consortium, the Thomas Gingeras Laboratory (Cold Spring Harbor Laboratory), the Ali Mortazavi Laboratory (University of California Irvine) and the Joe Ecker Laboratory (Salk Institute for Biological Studies) for generating and sharing the ENCODE data used in this study. We would also like to thank Megan Dennis (University of California Davis) for generating and providing access to the CHM13 RNA-seq data on behalf of the T2T consortium. Finally, we would like to thank Jean Monlong and Glenn Hickey for feedback on the manuscript, and everybody else in the VG Team.

## Methods

### Sequencing data, transcript annotations and variation databases

GENCODE v29 (primary assembly) was used as a transcript annotation set [33]. All transcripts with either the mRNA_start_NF or mRNA_end_NF tag were removed in order to only keep confirmed full-length transcripts. Furthermore, a transcript subset containing 80% of the GENCODE transcripts was created by randomly removing 34,490 of the 172,449 transcripts in the annotation. The fraction removed was based on recent estimates of the fraction of novel transcripts in a sample using long reads [32].

Genomic variants on GRCh38 from the 1000 Genomes Project (1000GP) were downloaded from EBI (http://ftp.1000genomes.ebi.ac.uk/vol1/ftp/release/20130502/supporting/GRCh38_positions/) [34]. The variants were first normalized using bcftools [66] and four different sets containing variants from differently-sized collections of samples were created (Supplementary Table 1). Two of these sets were constructed so as to not include variants unique to the CEU population. This was because we benchmarked the pipeline on NA12878, who is from this population, and we wanted our evaluations to cover one of the intended use-cases for the pipeline: sequencing a new sample from a population that is not represented in the reference haplotype panel. For all of the variant sets except the personal sample-specific set (where allele frequency was not relevant), the intronic and intergenic variants were further filtered using bcftools, keeping only variants with an alternative allele frequency of at least 0.002 or 0.001 depending on the set. This was done to decrease the complexity of the graph in regions where fewer reads are expected to map. The GRCh38 (primary assembly) reference genome used throughout the study was downloaded from Ensembl (ftp://ftp.ensembl.org/pub/release-94/fasta/homo_sapiens/dna/).

A list of all sequencing data used can be found in Supplementary Table 4.

### Spliced pangenome graph construction

We developed a method in the vg toolkit, vg rna, for constructing spliced pangenome graphs from a transcript annotation and an existing pangenome graph. vg rna begins by identifying the path in the graph that corresponds to each exon in the annotation. This process is facilitated by indexes in the vg toolkit that can efficiently query graph locations of positions on the linear reference. These exon paths can start or end internally in a node rather than only at boundaries between nodes, as with other paths in vg. Next, vg rna divides nodes as necessary to expose exon boundaries as node boundaries and then adds edges (splice-junctions) to the graph connecting adjacent exons within each transcript. The transcript paths are then labeled in the resulting spliced pangenome graph. Lastly, the spliced pangenome graph’s node ID space is compacted and reordered in topological order to make graph compression more efficient [67].

Different combinations of transcript annotations (full and an 80% random subset) and variant sets (Supplementary Table 1) were used as input to create the graphs used in the mapping and expression inference evaluation.

### Pantranscriptome construction

In addition to constructing spliced pangenome graphs, vg rna can simultaneously generate pantranscriptomes consisting of haplotype-specific transcripts (HSTs) created from transcript and haplotype annotations. It creates pantranscriptomes by projecting the reference transcript paths onto haplotypes paths that are either labeled in the graph or indexed using the Graph Burrows-Wheeler transform (GBWT) [31]. The GBWT is a succinct data structure for efficiently storing thousands of paths in a graph, such as haplotypes or transcripts. If nodes are split during the spliced pangenome graph construction (see above), vg rna first updates the haplotypes in the input GBWT. Next, the flanking positions of the exon boundaries on the reference chromosome path are located in the graph. These positions are used as anchors for projecting exons between the reference and haplotype paths. Anchoring on the positions adjacent to exon boundaries allows for genomic variation at the distal ends of exons.

Depending on whether the haplotype paths are labeled in the graph or stored in a GBWT, the projection is performed differently. For haplotype paths that have been labeled in the graph, we first locate all paths that contain both anchor nodes for each exon in a transcript. Next, for each located exon anchor pair we then follow the haplotype path between the two anchors to create the projected haplotype-specific (HS) exon path. HST paths are then created by combining all HS exon paths that are projected to the same haplotype. Only complete transcripts where all exons are successfully projected are kept. A projection will fail if there is variation at the anchor position in the target haplotype. Finally, HST transcripts that are identical are collapsed, producing a set of unique HSTs for each reference transcript. Since the number of pre-collapsed HSTs can be as large as the number of haplotypes, the algorithm is not suitable for large haplotype sets. For these, the GBWT-based algorithm, described below, is a better choice.

A broadly similar approach is used when the haplotype paths are stored in a GBWT. However, it differs in how the projected exons paths are constructed and combined. To find all possible haplotype paths between two exon anchors, we use a depth-first search (DFS). The search is initialized at the start anchor and traverses all possible paths in the graph starting from that anchor. Each explored exon path in the DFS (branch) is queried against the GBWT index and is terminated if it is not a subpath of any haplotypes in the index. Furthermore, a branch is also terminated if it is not possible to reach the end anchor node by any of the haplotypes consistent with the exon path. This is determined by examining whether any of the haplotypes containing the exon path also contain the end anchor node. The output from the search is a list of unique projected HS exon paths and the set of haplotypes consistent with each of them. The final HST paths are constructed one exon at a time by connecting HS exon paths that share at least one haplotype for each transcript. Because all the HS exon paths are unique this procedure will always result in a set of only unique HST paths and thus it is not necessary to collapse identical paths. This attribute makes the approach using the GBWT scale well with the number of haplotypes, as it can take advantage of the fact that haplotypes are often identical locally.

A list of all pantranscriptomes created for this study including the transcript annotations and variant sets used as input can be seen in Supplementary Table 3. The HSTs were written both as nucleotide sequences in FASTA format and as paths to a GBWT. A bidirectional GBWT, where each path is stored in both directions, was also created. rpvg uses this index to decrease computation time when reads are not strand-specific. For each GBWT, a corresponding *r*-index was further constructed. This index, based on the original *r*-index by Gagie et al., significantly decreases the computation time it takes to query path IDs in the GBWT [68].

### Read simulation model

Most of the simulated reads were generated using vg sim, a read simulator in the vg toolkit that is designed primarily for next-generation sequencing (NGS) reads. Its model consists of three components: a Markov model for base quality strings, a path frequency model, and a fragment length model (when sampling paired-end reads).

The model for base quality strings is fit to replicate the base quality strings in a user-provided FASTQ. A separate Markov transition distribution is fit for each base position in the read. The state of each Markov distribution consists of two components: the Phred base quality at that base and whether that base is an N. If a paired-end FASTQ is provided, vg sim will fit a separate model for each read end. In addition, the first states of each read in the pair are modeled with a single joint distribution, which allows for some dependence between the quality of both reads in the pair. The probabilities of the Markov transitions and the initial states are estimated by their empirical frequency in the FASTQ.

vg sim determines the base sequence of each read by following random walks through the pangenome graph. These walks may optionally be restricted to specific paths through the graph. Importantly for this study, the simulation can be restricted to the paths of transcripts in a spliced pangenome graph. The sampling frequency of a transcript path is proportional to the product of its effective length [4] and its expression value measured in transcripts per million (TPM), as determined by a user-provided expression profile. Once the path has been chosen, the starting location of the read is selected uniformly at random along the transcript. The sequence of the walk is then extracted, and sequencing errors are introduced according to the probability distribution implied by the base quality string. A user-specified fraction of these errors are produced as indel errors rather than substitution errors.

When simulating paired-end sequencing, the fragment length is modeled with a normal distribution. The user provides the mean and standard deviation for this distribution. For a given path, the normal distribution is truncated to between 1 and the path length. Both reads are sampled from a single walk through the graph with length equal to the sampled fragment length. If this length is shorter than the read length, the read is truncated to the fragment length.

### Simulating RNA-seq reads from haplotype-specific transcripts

Reads were simulated from haplotype-specific transcript (HST) paths derived from the haplotypes of NA12878 in the 1000 Genomes Project (1000GP) and the GENCODE transcript annotation. The corresponding spliced pangenome graph (including the paths) was created using vg rna. Identical HSTs were not collapsed, so that reads could be simulated from each haplotype independently.

In total, we created five different simulated read sets (Supplementary Table 5): four using vg sim and one using RSEM. Two different read sets were used to fit the simulation model: SRR1153470 and ENCSR000AED. For both real read sets, we used vg sim to create two simulated read sets. One set of reads was simulated with an expression profile derived from the real data, and the other set was simulated with uniform expression across transcripts. The single RSEM simulation used the uniform approach. The simulated read sets with uniform expression were used to evaluate mapping, whereas the sets with data-based expression were used to evaluate expression inference.

To ensure balanced expression between the two haplotypes for all transcripts, only transcripts that were successfully projected to both haplotypes were given a positive expression for the uniform expression set. For the simulated read sets with data-derived expression, the reads were first mapped using Bowtie2 [14] with default parameters and then quantified using RSEM [1], also with default parameters. The fragment length distribution mean and standard deviation estimated by RSEM were used to parameterize the fragment length distribution in vg sim. For all five read sets, we simulated 25M 101 base-pair read pairs from each haplotype. For vg sim we used an indel probability error of 0.001 and the base quality distribution was trained on 10M randomly sampled read-pairs of the training data. The read-pairs were sampled using seqtk [69]. RSEM was given the estimated training data model file and the background noise fraction was set to zero.

### Mapping and multipath alignment with vg mpmap

Like most read mappers, vg mpmap’s mapping algorithm is designed using the “seed-cluster-extend” paradigm. First, it locates exact matches “seeds” between the read and the graph. Next, the seeds are “clustered” together to identify regions of the graph that the read could align to. Finally, the seeds are “extended” into an alignment of the entire read. Because these operations occur in the context of a pangenome graph, they use several specialized algorithms and indexes.

#### Seeding

vg mpmap seeds alignments with maximal exact matches (MEMs) against the graph, which it finds using a GCSA2 index [70]. MEMs are exact matches between an interval of the read and a walk in the graph such that the match cannot be extended further in either direction at that location in the graph. The MEMs are found using a two-stage algorithm, which has also been described previously [8].

In the first stage, the algorithm finds super-maximal exact matches (SMEMs), which are MEMs for which the read interval is not contained within the read interval of any other MEM (Supplementary Algorithm 1). This algorithm also relies on a longest common prefix (LCP) array, which allows navigation upward in the implicit suffix tree that the GCSA2 encodes. The second stage of the algorithm finds the minimally-more-frequent MEMs of each SMEM, subject to a minimum length (Supplementary Algorithm 3). These are the longest MEMs that are shorter than the SMEM but have their read interval contained in the SMEM’s read interval. This stage also relies on the GCSA2 index.

#### Clustering

The clustering algorithm in vg mpmap is built around the distance index described in [71]. In brief, this index can query the minimum distance between two positions in the pangenome graph by expressing the distance as the sum of a small number of precomputed distances. This is accomplished by taking advantage of the common topological features of pangenome graphs, namely that they tend to contain long chains of bubble-like motifs that result from genomic variation. These features are captured in the graph’s “snarl decomposition”, in which a snarl is one of these bubble-like motifs [72].

The clustering algorithm begins by constructing a directed acyclic graph (DAG) in which the nodes correspond to MEM seeds. The edges are added whenever 1) there is a path connecting two seeds in the graph, and 2) the seeds are collinear along the read. Note that the collinearity criterion guarantees acyclicity. We use the distance index to determine the existence of a path that connects the seeds in the graph, and the edges are also labeled by the distance. Edges that are much longer than the read length are not added; this avoids treating distal elements on the same chromosome as part of the same cluster. In addition, we accelerate this process using Algorithm 3 from [71], which partitions seeds into equivalence classes based on the distance between them. The equivalence relation is the transitive closure of the relation of being connected by a path of length less than *d*, which is a tunable parameter. By choosing *d* correctly, we can ensure that all of the edges we would include occur between seeds in the same equivalence class. This significantly reduces the number of distance queries we need to perform.

Once the DAG of seeds has been constructed, we approximate the contribution of each seed and edge to the score of an alignment that contains them. Seeds are scored as if they are a short alignment of matches, and edges between seeds may be scored as an insertion or deletion if the distance in the graph does not match the distance on the read. These values serve as node and edge weights. We then use dynamic programming to compute the heaviest path defined by the node and edge weights within each connected component of the DAG and take the seeds along this path as a candidate cluster. Clusters are passed through to the next stage of the algorithm if their weight is within a prespecified amount of the heaviest-weight cluster, subject to a hard limit on the total number of clusters.

#### Multipath alignments

Most existing sequence-to-graph aligners, including vg map, produce an alignment of the sequence to a particular path through the graph. vg mpmap uses a different alignment formalism, which we call a multipath alignment. In a multipath alignment, the sequence can diverge and reconverge along different paths through the graph (Supplementary Figure 1). Thus, the read can align to a full subgraph rather than to a single path. This allows the alignment object to carry within itself the alignment uncertainty at known variants or splice-junctions. This information can be used in downstream inference applications, including rpvg.

More formally, a multipath alignment of read *R* is itself a digraph with the following properties:

1. Each node corresponds to an alignment of some substring of *R* to a path in the pangenome
2. An edge between *u* and *υ* exists only if *u* and *υ* align adjacent substrings of R to adjacent paths in the pangenome.
3. Every source-to-sink path through the multipath alignment can be concatenated into a complete, valid alignment of R to a path in the pangenome.

It is worth noting that multipath alignments are acyclic by construction, since the nodes can be partially ordered by the read interval that they align. vg mpmap additionally annotates each node’s partial alignment with its alignment score. The alignment score of any particular sequence-to-path alignment expressed in the multipath alignment can be computed efficiently by simply adding the partial alignments scores along the path.

While sequence alignments have well-established optimization criteria, there is no such criterion for optimizing the topology of a multipath alignment. In lieu of one, we adopt heuristics that are motivated by the common topological features of pangenome graphs. Our high-level strategy is to use exact match seeds to anchor alignments. We then use dynamic programming to align between seeds and within sites of variation in the graph, which we identify using the snarl decomposition of the pangenome graph. Using a multiple-traceback algorithm, we can then obtain alignments to different paths through the graph as necessary.

#### Anchoring alignments

To use a cluster of exact match seeds to anchor a multipath alignment, it is first necessary to compute the reachability relationships between the seeds. This is a non-trivial problem.

We begin by converting the local graph around a cluster into a directed acyclic graph using an algorithm that has been described previously [8]. In brief, we identify small feedback arc sets within each strongly-connected component using the Eades-Lin-Smyth algorithm [73], and then we duplicate the strongly-connected component with the feedback arcs linking successive copies. Using dynamic programming over the DAG as we construct it, we can preserve all cyclic walks up to some prespecified length, which is based on the read length.

After creating the DAG, we inject the seeds into the new graph. Since the DAG conversion algorithm can expand the node space of the original graph, seeds can now correspond to multiple locations in the DAG. In this case, we duplicate the seeds to all of the corresponding locations in the DAG. We then use a three-stage algorithm that computes the transitive reduction of a graph in which the nodes correspond to seeds, and two seeds have an edge between them if they are collinear along the read and reachable within the pangenome graph (Supplementary Algorithm 4).

1. Compute the reachability relationships between the seeds, ignoring collinearity on the read.
2. Rewire the reachability edges between the seeds to respect collinearity on the read.
3. Compute the transitive reduction of the resulting graph.

This algorithm is designed to have linear run time in the number of seeds and the size of the DAG, but only in the typical case where the seeds line up along a walk through the pangenome graph. In the general case, the run time can be quadratic.

#### Dynamic programming with multiple traceback

The alignments between anchors (i.e. the vertices in the transitively reduced DAG) are computed using a banded implementation of partial order alignment [74]. The alignments of the read tails past the end of anchors are computed using a SIMD-accelerated POA implementation from the gssw library.

We use a specialized traceback algorithm to obtain the alignments to multiple paths through the pangenome graph from a single dynamic programming problem (Supplementary Algorithm 8). Instead of the optimal alignment, the algorithm returns the *k* highest-scoring alignments. We choose *k* to be the number of paths through the subgraph we are aligning to, subject to a hard maximum. The key insight behind the algorithm is that the next highest-scoring traceback can be determined by checking local properties of the dynamic programming matrix while computing the highest-scoring traceback. In addition, for each anchor that crosses a snarl, we remove the interior of snarl before performing alignments. This way, the multiple traceback algorithm can align to multiple paths at sites of variation.

#### Quantifying mapping uncertainty

The method that vg mpmap uses to compute mapping quality is largely shared with vg map (see [8] Supplementary Note). As in vg map, base qualities are incorporated into alignment scores (essentially downweighting low-quality bases), and the alignment scores are subsequently used to compute a mapping quality. The formulas used to compute mapping quality rely on the conversion of alignment scores into the log-likelihood of a hidden Markov model (HMM), as described in [75] and [76].

vg mpmap also uses a concept of a mapping’s “multiplicity” to model errors introduced by the mapping algorithm itself. In particular, at certain points in the algorithm, we enforce hard caps on certain algorithmic behaviors, such as the number of alignments that will be attempted, in order to prevent excessive run time. If we run up against these hard caps, we expect that not all high-scoring alignments will be found. We incorporate this information into the mapping quality formula by treating alignments as if multiple equivalent alignments actually were found. For example, if we attempted alignments for 10 of 30 promising clusters and found 1 high-scoring alignment, we would estimate its multiplicity to be 3. That is, we estimate that there are a total of 3 alignments that are equally high-scoring, including the ones that we did not find. We then compute the mapping quality as if 2 additional copies of the alignment had been found.

Multiplicities allow vg mpmap to aggregate information about sources of algorithmic inaccuracy over different steps in the algorithm. The central entities in each step of the mapping algorithm (seeds, clusters, alignments, and pairs) are each associated with a multiplicity. These multiplicities follow particular combining rules between successive steps of the algorithm. When combining orthogonal pieces of information (seeds in a cluster, or single-end alignments in a paired alignment), the new entity receives the minimum of its constituents’ multiplicities. When layering on a new source of algorithmic uncertainty (typically a further hard cap), an entity’s multiplicity is multiplied by its estimated multiplicity in that step of the algorithm.

#### Determining statistical significance

vg mpmap uses a frequentist hypothesis test to assess the statistical significance of a read alignment. The test statistic that we use is the alignment score. The null hypothesis is that the alignment score was obtained by a uniform random sequence of the same length as the read. By default, we set the type-I error rate to 0.0001. If an alignment score’s *p*-value is not significant at this level, the read is reported as unmapped.

Modeling the null hypothesis of the test is not entirely straightforward. In general, we expect higher local alignment scores from longer reads or larger pangenome graphs. However, there are subtleties. A large pangenome graph may consist of many repeats of the same sequence so that its effective size is smaller than its total sequence length. Alternatively, a small pangenome graph may have a complex topology that admits a combinatorially large set of walks. For these reasons, we take an empirical approach that fits a model to match the pangenome graph. At the start of every mapping run, we map a sample of uniform random sequences of varying lengths. The resulting alignment scores are used to fit the parameters of a distribution using maximum likelihood, and those parameters are regressed against the read length. The regression allows us to query the *p*-value for a read of any length.

The parametric distribution we use can be derived as the maximum of *ν* independent, identically distributed (i.i.d.) exponential variables with rate λ. This distribution has the following probability density function:

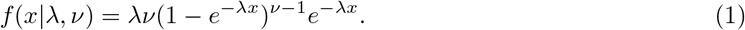

The fitting algorithm alternates between maximizing the likelihood with respect to each of the two parameters with the other fixed until convergence. *ν* is fit using the Newton-Raphson method, and λ is fit using golden-section search.

The motivation for this model is that the length of the match starting at position of a uniform random sequence (the read) and position of a fixed sequence (the reference) is approximately Geometric(1/4), assuming the two sequences are relatively long. The optimal local alignment score is closely related to the longest match at any position on the read sequence to any position on the pangenome graph. Moreover, most of these matches have only weak dependence on each other, so the i.i.d. approximation is reasonable. We use an exponential distribution because it closely approximates a geometric distribution and is easier to fit.

#### Paired-end mapping

vg mpmap has several features designed to take advantage of the paired-end sequencing reads produced by Illumina sequencers. At the beginning of each paired-end mapping run, vg mpmap uses a sample of the first 3,000 uniquely mapped pairs to fit parameters of a fragment length distribution. The distance between the reads in each pair is computed with the distance index. Non-uniquely mapped pairs are buffered and then remapped after the fragment length distribution has been fit.

The fragment distribution is modeled as a normal random variable with mean *μ* and variance *σ*^2^. We use a method of moments estimator for a truncated normal distribution so that the parameter estimation is robust to possible mismappings. In particular, we discard the largest and smallest 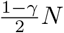 fragment length measurements (default *γ* = 0.95). This procedure makes the estimator insensitive to a sufficiently small fraction of outliers that can result from mismapping or mapping across unannotated splice junctions. The remaining *γN* measurements correspond to a sample from a truncated normal distribution with the same *μ* and *σ*^2^. The following estimators can be derived using method of moments on this truncated normal distribution:

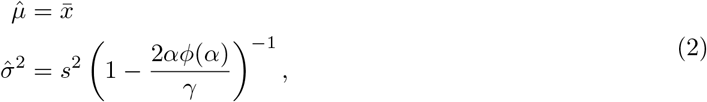

where 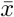 and *s*^2^ are the empirical mean and variance among the retained *γN* measurements, and 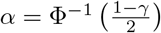 is the left truncation point on a standard normal distribution.

When mapping paired-end reads, the clustering stage of the algorithm adds an additional step. First, each read in the pair’s seeds are clustered as in the single-end algorithm. Next, the clusters from the two reads are paired by checking which pairs imply a fragment length within 10 standard deviations of the mean, as estimated by the algorithm in the previous section. The implied fragment length connecting two clusters is estimated using the distance index, with the position of a cluster taken to be the position of its longest seeds. Pairs of clusters are prioritized by a sum of an estimated alignment score (interpreted as a log-likelihood) and the log-likelihood of the normal distribution that we model the fragment length distribution with.

The heuristics used for read mapping inevitably fail in some cases. When mapping paired-end reads, it sometimes happens that the heuristics fail on only one of the two reads of a fragment. When this occurs, it is sometimes possible to “rescue” the alignment of the other read by aligning it to the region of the pangenome graph where we expect to find it relative to the mapped read.

vg mpmap employs this strategy whenever the pair clustering procedure fails to produce a pair of clusters consistent with the fragment length distribution, or when all of the clustered alignment pairs have at least one end without a statistically significant alignment. We also perform a limited number of rescues even when a consistent cluster pair is found, provided that there are clusters of at least one of the ends that are equally as promising as the one in the cluster pair. This helps improve the calibration of mapping qualities. We place a hard cap on the number of rescues performed to control run time. The fraction of eligible rescues that were actually performed becomes a component in the multiplicity of an alignment, as described previously.

The multipath alignment algorithm is slightly different when computing rescue alignments. This is necessary because there are no exact match seeds to use as anchors. Instead, we first perform a single path alignment using gssw. Then we remove any sections of the alignment that lie inside snarls, and realign those segments of the read as when connecting anchors in the standard multipath alignment algorithm.

#### Spliced alignment

Because spliced pangenome graphs include annotated splicing events as edges, it is usually unnecessary to use specialized alignment algorithms to obtain spliced alignments. However, even for well-annotated genomes, transcript annotations are incomplete, especially for lowly-expressed transcripts, so it is still important to be able to produce spliced alignments. vg mpmap includes a spliced alignment algorithm but applies it conservatively: only when an alignment includes a moderately long soft-clip on at least one end. A long soft-clip is suggestive that the clipped end of the read might align to a part of the graph that was too distant to be included in the primary seed cluster, as would be expected with an unannotated splice event. One common exception to this pattern are adapter sequences, which can be captured in a read when the sequenced fragment is shorter than the read length. To avoid the computational burden of attempting to find nonexistent spliced alignments for these cases, common adapter sequences are specifically excluded from this subroutine.

The spliced alignment algorithm begins by finding candidate regions to align the clipped read end to. These regions are selected by scanning over secondary mappings, unaligned seed clusters, and unclustered seeds. For paired reads, spliced alignments can also be found by rescuing the soft-clipped portion of the read from the other read in the pair. This is only possible when the soft-clip is on the side of the read that faces inward on the fragment. In addition, spliced alignment rescue is only attempted when none of the other spliced alignment candidates yields a statistically significant spliced alignment.

Spliced alignment candidates must pass several filters to be included in the read mapping. Candidates are filtered out unless they roughly correspond to the clipped end of the read. They must also be reachable from the primary alignment by some path in the graph, which is determined using the minimum distance index. The final filter that a spliced alignment must pass is a significance test. This test has three components: 1) the increase in alignment score that results from aligning the additional bases, 2) the bias against the splice site motifs in the intron, and 3) the bias against the intron length. Splice site motifs are penalized by their log-frequency, as given by Burset, et al. [77]. The bias against the intron length is determined using the log-likelihood of a log-normal mixture model fit to the human intron length distribution. The three components are combined into a joint log-likelihood and tested against a critical value.

To compute the quantities needed for this test, the spliced alignment algorithm identifies the splice motifs near the ends of a pair of splice candidates. If any pair of canonical splice site dinucleotides are found on any path from the two ends, the intervening sequence is aligned as if the two splice sites were joined by an edge in the graph. In addition, the intron length is measured between these positions in the graph using distance along the reference path.

#### Multimapping reads

Reads with multiple high-scoring alignments can be reported in two different ways. First, separate alignments can be reported up to a user-specified maximum number. Second, a single multipath alignment can be reported that includes all high-scoring alignments. This multipath alignment may be disconnected. In the first option, all of the reads are annotated with a collective “group mapping quality” that quantifies the probability that all of the reported alignments are incorrect. In the latter option, the main mapping quality annotation is equivalent to the group mapping quality.

### RNA-seq mapping evaluation

We compared vg mpmap’s performance at mapping RNA-seq data against the vg toolkit’s existing graph alignment method vg map [8] and two state-of-the-art RNA-seq mapping tools, HISAT2 [19] and STAR [2]. Graph indexes and genomes were created for each tool using default parameters, with mpmap and map sharing the XG and GCSA index. The mapping compute and memory usage of each tool were estimated using 16 threads on an m5.4xlarge AWS instance. All mappers were run with default or recommended parameters for RNA-seq data. For the simulated data the maximum number of reported multi-alignments per read was set to same value for each method. A value of 10 was used since it is the default value of both HISAT2 and STAR. The SRR1153470 and CHM13 data were used to optimize the parameters of vg map and vg mpmap.

We evaluated mapping accuracy on simulated reads using two different methodologies to ensure the robustness of our conclusions. One methodology was based on basewise overlaps along the linear reference genome, and the other was based on distances along transcript and reference paths in the graph.

For the overlap-based evaluation the graph alignments were first projected to the reference paths using vg surject in spliced alignment mode. Briefly, surject takes a set of graph-aligned reads and re-aligns them to all nearby reference paths in the graph, producing a BAM file with the reads aligned to the reference sequences. The re-alignment is only performed on the parts of the alignment that do not already follow the reference paths. A read was considered correctly mapped if 90% of the bases of the simulated true reference alignment were covered by one of its estimated multi-alignments. The true reference alignments were generated using the transcript position of each read provided by vg sim or RSEM, and the NA12878 haplotype-specific transcript reference alignments. The latter were created by projecting the transcript paths to the reference sequences using vg surject in spliced alignment mode.

Due to sequencing artifacts, the ends of reads will occasionally consist of such low-quality bases as to be practically random. Our simulation framework recapitulates this feature of real sequencing data. However, in real data these read ends do not correspond to any underlying genomic sequence, whereas the simulation assigns them a true genomic alignment. Aligners that softclip these uninformative bases would be penalized in this evaluation, even though this is the correct decision for real data. We therefore decided to trim all bases at both ends of an alignment (including the true alignments) that had a Phred base quality score below 3 for the purpose of computing the overlap. All alignments for which more than half of the sequence was trimmed were discarded from the evaluation so that the percent overlap could be estimated more confidently.

To classify whether a read contained any novel (unannotated) splice-junctions we looked at all deletions and reference skips in the true reference alignment with a length of at least 20 bp. These were compared to the transcript annotation that was used to build the graph or reference, and defined as novel if it was not possible to find a splice-junction in the annotation that was within 5 bp at both ends.

We used the vg gampcompare tool for the distance-based evaluation. The truth set in this evaluation was the true graph alignments produced by vg sim. In short, vg gampcompare finds the minimum possible distance between the start position of an estimated alignment and the true alignment across all reference and transcript paths in the graph. Before running GAMPCOMPARE, HISAT2 and STAR’s BAM format alignments were converted into graph alignments (GAM format) using vg inject, which translates linear reference alignments into alignments against the path of the reference in a graph. An alignment was considered correct if its start position was within 100 bp of the start position of the true alignment along the path of the reference or any transcript path.

Reference bias was quantified using simulated reads, by counting the number of reads that overlapped variants with a mapping quality value of at least 30. For this analysis we used the reference-based alignments (projected alignments for vg map and vg mpmap). In order to treat different variant types and lengths equally, we computed the read count for each variant allele as the average read count across the allele two breakpoints. Reads simulated from each haplotype were counted separately and only variants with at least 20 reads across both alleles combined were used to quantify reference bias. Complex variants that were not classified as SNVs, simple deletions or simple insertions were skipped.

To further evaluate allelic bias, we counted the reads supporting each allele of heterozygous variants among mapped simulated reads for each of the mappers using the approach described above. We also added a pipeline consisting of STAR followed by read filtering with WASP to this comparison. We found that WASP was computationally infeasible using the full CEU population, so we instead gave it a variant database consisting of only NA12878’s own variants. Therefore, to have a better comparison to WASP, we also created personal sample-specific (NA12878) references for vg mpmap, vg map, and HISAT2, and we report results for these references as well. We estimated the observed rate of false positives by testing for allelic skew in mapped reads on heterozygotic variants using a two-sided binomial test (*α* = 0.01). All significant *p*-values are false positives, since the reads were simulated without an allelic bias.

When benchmarking using real reads, truth alignments are not available. Instead, we used a proxy measure of aggregate mapping accuracy based on long read mappings from the same cell line. The long reads are easier to map confidently, and we expect the cell line to have similar transcript expression across replicates. Thus, higher correlation between the coverage of short read mappings and the coverage of long read mappings is indirect evidence of higher accuracy. For long read data, we used NA12878 PacBio Iso-Seq alignments generated by the ENCODE project (Supplementary Table 4). The cleaned Iso-Seq alignments of four replicates were first merged and secondary alignments and alignments with a quality below 30 were filtered using samtools [78]. These filtered alignments were then compared to the short-read RNA-seq alignments by calculating the Pearson correlation of the average exon read coverage between the two. Exons were defined using the Iso-Seq alignments by first converting them to BED format and then merging overlapping regions using bedtools [79].

We measured memory and compute time for all mappers using the Unix time utility. The reads per second statistic was computed by dividing the number of reads by the product of the wall clock time and the number of threads. This is a somewhat biased measurement, since it includes the one-time start up computation that does not scale with the number of reads. However, the magnitude of this bias is small, and it tends to disfavor vg mpmap, which has the longest start up of the tools we evaluated.

Reference alignments in BAM format were sorted and indexed, also using samtools. The SeqLib library was used in the evaluation scripts to parse the alignments and calculate overlaps [80].

### Haplotype-specific transcript quantification

We developed rpvg as a general tool for inferring the most likely paths and their abundance from a set of mapped sequencing reads. In this study we used rpvg to quantify the expression of haplotype-specific transcripts (HSTs) in a pantranscriptome. rpvg’s algorithm consist of four main steps:

1. Find read alignment paths that align to HST paths
2. Cluster alignment paths and HST paths
3. Calculate alignment path probabilities
4. Infer haplotypes and expression from probabilities

A graphical overview can be seen in Supplementary Figure 27.

#### Finding alignment paths

The first step of rpvg is to parse each alignment and find all alignment paths that align to (i.e. follow) at least one HST path in the pantranscriptome GBWT index (Supplementary Figure 27a). An alignment path is the set of nodes a read alignment follows in the graph. For single-path alignments there is only one alignment path, but for multipath alignments there can be many. We will here focus on multipath alignments, since a single-path alignment is merely the simpler case when a multipath alignment only contains a single path.

Multipath alignments are represented as a graph, and thus the objective is to find all paths through this graph that also exist as subpaths in the GBWT. In other words we want to find all possible alignments that the read can have to all HST in the pantranscriptome. This search would normally scale linearly in the number of HSTs overlapping the read, but the GBWT allows us to query all HSTs that contain the same subpath. Therefore, HSTs that are locally identical will be queried together, taking advantage of the fact that haplotypes are markedly more similar locally than globally.

rpvg uses a depth-first-search (DFS) through the multipath alignment graph to find all alignment paths. A branch in the search is terminated if its alignment path is not present as a subpath in the GBWT. A DFS is initialised at each source node in the alignment graph. We terminate any alignment path early where it is not possible to reach a score of 20 below the current highest scoring path, assuming perfect scoring for the remainder of the alignment.

The topology of the multipath alignment graphs is determined by heuristics. In some cases these heuristics fail, resulting in multipath alignments that do not cover all possible alignment paths. This can result in incorrect downstream expression estimates as a read might be missing an alignment path to the correct HST. To overcome this, rpvg allows alignment paths to be shortened in order to be made consistent with an HST path. More specifically, the DFS can start and end up to four bases inside the read (excluding soft-clipped bases). The score of partial alignment paths are penalized proportionally to the number of non-matched bases at each end, adjusted for their quality. The longest possible alignment path to a HST is selected as the best alignment.

The output from the DFS is one set of alignment paths for each multipath alignment. Next, rpvg labels a set as low scoring if the highest scoring alignment path in the set is less than 0.9 times the optimal quality-adjusted alignment score, which is the score an alignment would get if it consisted of only matches. The sets labeled as low scoring are treated as being incorrect; they may be misalignments, or originate from an HST not in the input pantranscriptome. The labeled sets are later used when calculating the noise probability.

For paired-end reads, one additional step is needed: combining the alignment paths of each read to create a set of alignment paths for the whole fragment. First a set of alignment paths is generated for each alignment in the pair as described above. Next, rpvg attempts to combine each start (first read) alignment path with each of the end (second read) alignment paths. If the fragments are not strand-specific and the pantranscriptome GBWT is not bidirectional, rpvg then repeats the process using the reverse complement of the fragment.

The procedure to combine the two alignment paths differs depending on whether they overlap or not. If they do overlap, a single combined alignment path is created for the fragment by merging the two while requiring that the path of overlapping portions matches perfectly. If they are separated by an insert, the start alignment path is extended using a depth-first search following the HST paths. If the search reaches one of the start nodes for an end alignment path, a new fragment alignment path is created by merging the search and end alignment path. The new fragment alignment path is only kept if it follows at least one HST path in the pantrancriptome. The search is terminated if all start nodes in the end alignment paths have been visited and they are not part of a cycle. An alignment path is discarded if its length is above *μ* + 10*σ*, where *μ* and *σ* are the mean and standard deviation of the fragment length distribution. These parameters are either supplied by the user or parsed from the input alignments (the vg aligners write the parameters they estimated to the alignment file). The score of the resulting fragment alignment path is calculated as the sum of the scores of the two read alignment paths. The mapping quality is calculated as the minimum across the two reads.

The final output from the search is a set of alignment paths and the HSTs that each path aligns to for each read or fragment. For simplicity, in the following, we will use the term “fragment” to denote both a single-end read and a set of paired-end reads.

#### Clustering transcript paths

HST paths that do not share any fragments are independent, and therefore their expression can be inferred separately. In contrast, the expression of HST paths that share alignments must be inferred jointly. Accordingly, rpvg identifies clusters of HST paths that share alignment paths from the same fragment. By dividing the inference problem into these smaller, independent clusters, computation and memory can be considerably reduced.

The clustering algorithm works by first constructing an undirected graph where vertices correspond to HST paths and edges correspond to HST paths being observed in the same set of fragment alignment paths. All connected components in this graph (clusters) are then located using breadth-first-search. All fragments are assigned to their respective clusters based on the HST paths that their alignment paths align to.

#### Calculating alignment path probabilities

For each fragment, the probability of it originating from each of the HSTs in its cluster is calculated by rpvg using the alignment path scores, lengths and mapping quality (Supplementary Figure 27b). First the probability *ϵ* that the fragment was not from any of the HST in the cluster is calculated using the mapping quality *q*:

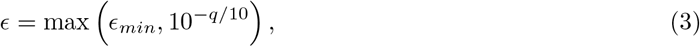

where *ϵ_min_* is the minimum noise probability. The motivation behind having a minimum is that mapping qualities are generally less reliable at higher values. The minimum noise probability is 10^−4^ for all fragments except those that were labeled as low scoring, for which it is 1. Now, let *A* be the set of alignment paths (i.e. alignments) for this fragment. For each alignment path *a* ∈ *A*, the likelihood of it being the correct path is calculated using its score *s_a_* and length *ℓ_a_*:

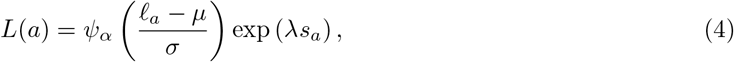

where λ is a scaling factor that converts the alignment score into the log-likelihood of a pair-HMM [76], *ψ_α_*(*x*) = 2*ϕ*(*x*)Φ(*αx*) is the density of a skew-normal distribution [81] (with *ϕ* and Φ the density and distribution function of a standard normal distribution, respectively), and *μ*, *σ*, and *α* are the location, scale, and shape parameters of the fragment length distribution modeled as a skew-normal distribution. For paired reads, these parameters are estimated from the alignment path lengths across all fragments that have 1) a mapping quality of at least 30, and 2) the same length for all alignment paths. The fragment length distribution is omitted from the equation when the fragments are single-end reads. With this likelihood, we can compute the posterior probability that the fragment originated from a given HST. Let the set of all HST paths in the cluster be denoted by *T*, and let the set of HST paths an alignment path *a* is consistent with be denoted by *T_a_*. The probability that the fragment (or alignment *A*) originated from an HST is calculated as:

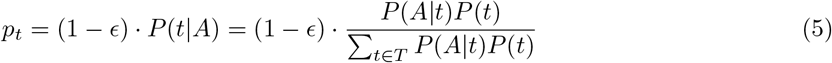

with

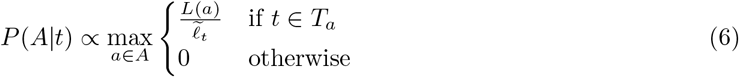

Here, 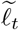 is the effective transcript length for *t* calculated as 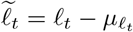 [3, 4]. In turn, *μ_ℓ_t__* is the mean of the fragment length distribution truncated to [1, *ℓ_t_*], computed using a published formula [82]. The effective transcript length accounts for the fact that fragments cannot be sequenced from all positions due to the size of the fragment. If the fragments are single-end reads, the fragment length distribution parameters used to calculate the effective length must be supplied by the user. The prior over HSTs *P*(*t*) is taken to be uniform. If the HST probability *p_t_* is below 10^−8^, it is truncated to 0 to reduce storage.

We denote the set of all fragment probabilities in a cluster as *F* and the probabilities for a fragment *i* as *F_i_* = (*ϵ*, **p**), where **p** is the vector of probabilities over all T HSTs in the cluster. Many fragments will have very similar probabilities and can thus be collapsed to save computation resources and memory [4, 83]. To do this we collapse two fragment probabilities *F_i_* and *F_j_* if they satisfy both of:

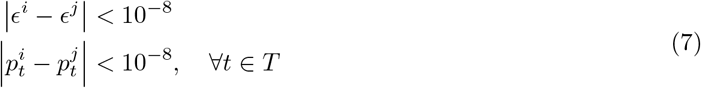

We also associate each set of collapsed fragments with *c*, the number of collapsed fragments in the set. The resulting set *E* of tuples (*ϵ*, **p**, *c*) is subsequently used to infer the expression of the HSTs in the pantranscriptome.

#### Inferring haplotype-specific transcript expression

RPVG quantifies the expression of the HSTs in the pantranscriptome using a nested inference scheme (Supplementary Figure 27c). This is done independently for each cluster. First, the distribution over haplotype combinations (i.e. diplotypes) is inferred. The most probable haplotype combinations are then selected from this distribution and expression is inferred conditioned on the haplotypes. In the following, we will assume the sample is diploid, but the equations and algorithms generalize to any ploidy.

The marginal distribution over diplotypes is approximated by assuming the haplotypes are identical for all transcripts in a cluster. The motivation behind this approximation is that most clusters cover only a small region (typically a gene) of the genome. However, this approximation can break down when there are partial haplotypes or recombination events in the cluster. Using the transcript and haplotype origin table provided by vg rna, the HSTs in the cluster are first grouped by their haplotype origin. Note that since an HST can be consistent with more than one haplotype it can also belong to multiple groups. Next, groups with the same set of HSTs are collapsed, resulting in a set of unique haplotype groups.

Now let us denote the set of haplotype groups as *H*, with each group *h* ∈ *H* consisting of a set of HSTs. The objective is to infer the distribution over diplotypes *d* = {*h*_1_, *h*_2_} conditioned on the set of collapsed fragment probabilities *E*. The probability of a diplotype is defined as:

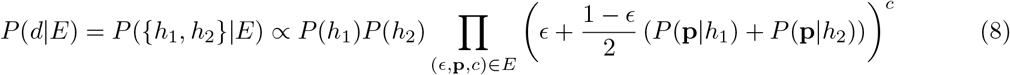

and

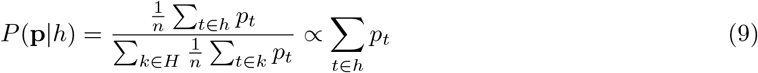

where the prior probability of each haplotype group *P*(*h*) is proportional to the number of haplotypes in the group, and *n* is the number of transcripts in the cluster (the factors of 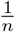 and 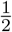 amount to an approximation that expression is uniform across all transcripts and the two haplotypes, respectively). This model is inspired by similar haplotyping models used in Platypus and other genotypers [84–86].

The distribution over diplotypes is inferred by calculating *P*(*d|E*) for all pairs of haplotype groups *h* ∈ *H*. To reduce the space of haplotype combinations that need to be evaluated, rpvg uses a branch-and-bound-like algorithm, where diplotypes containing an improbable haplotype group are not evaluated. Instead, the probability of all diplotypes containing an improbable haplotype group is set to 0. A haplotype group *h* is labeled to be improbable if its optimal diplotype probability *P*({*h*, *h_o_*}|*E*) is 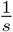 times lower than the current highest evaluated probability, where *s* is the minimum diplotype posterior probability threshold used in the next step in the inference. The optimal diplotype probability is defined as

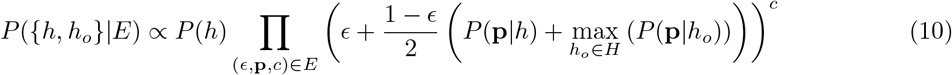

This value serves as an upper bound on the probability of any diplotype containing *h*.

The expression of the HSTs in the cluster is inferred using the inferred distribution over diplotypes. First, the set of diplotypes with a posterior probability of at least *s* = 10^−3^ is selected from the distribution *P*(*d|E*). HST expression is inferred for each of the diplotypes in this set.

The following is repeated for each diplotype in the set. First, all HSTs that are consistent with at least one of the haplotypes in the diplotype are collected. We denote this HST subset *T_s_* ⊆ *T* and define the likelihood over the relative expression values ***α*** as

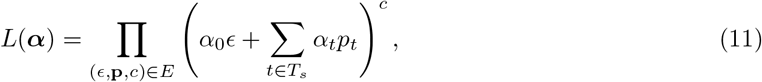

where *α*_0_ is the expression value of an artificial “noise transcript” that accounts for the possibility of mismapping. An expectation maximization (EM) algorithm is used to find the (local) maximum likelihood estimate of the expression values. The algorithm iterates between assigning fractional fragment counts to the HSTs and the noise transcript, and updating the expression values. This is a well known algorithm that is used by many other transcript quantification tools [1, 3, 4, 83]. The expression values are initialized uniformly and the EM algorithm is run until convergence or for a maximum of 10,000 iterations. The algorithm is considered converged if

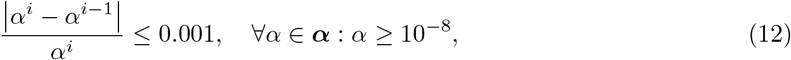

for 10 consecutive iterations, where *i* is the index of the current iteration. This criteria is inspired by the one used by Kallisto [3] and Salmon [4]. For the final maximum likelihood estimate, we truncate all the relative expression values below 10^−8^ to 0.

After the EM step, rpvg can optionally run a Gibbs sampling step to quantify the uncertainty in the expression estimates. The Gibbs algorithm samples the assignment of each fragment to a HST (or the noise transcript), and the expression values ***α***, which are given a symmetric uniform Dirichlet prior with a concentration parameter of one. A similar algorithm is described in Li et al. [1] and Patro et al. [4]. First, 1000 diplotypes are sampled from *P*(*d|E*), with a Gibbs sampler being run for each unique sampled diplotype. Each sampler is initialized on the maximum likelihood estimate from the EM algorithm and the number of samples of expression values collected is equal to the number of times the diplotype was sampled. This results in a total of a 1000 collected Gibbs samples of expression values for each cluster. In addition, we thin each Gibbs chain and only collects a sample of expression values at every 25th Gibbs iteration. This is done to reduce autocorrelation between samples.

rpvg provides both a joint and marginal output of the inferred probabilities and expression values. The joint output contains the inferred posterior probabilities over HST combinations (i.e. diplotypes) and their corresponding estimated expression values. Only combinations with a probability of at least 10^−8^ are written to this output. The marginal output is more similar to outputs of other quantification methods and contains the haplotype probability and estimated expression value for each HST in the pantranscriptome. The haplotype probability is calculated as the sum of posterior probabilities over all diplotypes that includes the HST. The HST expression reported is similarly calculated as the sum of the estimated expression values over all diplotypes that include the HST, weighted by their posterior probability. The expression of the noise transcript is aggregated across all clusters into a single artificial transcript called “Unknown”.

### Transcript quantification evaluation

We compared rpvg’s quantification accuracy against three other transcript quantification tools: Kallisto, Salmon and RSEM. Haplotype-specific transcript indexes for Kallisto, Salmon and RSEM were built from the HST sequence FASTA files generated by vg rna. For the real data, the 104 full-length mitochondrial and scaffold transcripts in the GENCODE v29 annotation were added to the pantran-scriptomes. Salmon indexing was run with duplicates kept and, on the real data, the reference genome was given as a decoy. The Bowtie2 mapper was used in RSEM with the maximum number of alignments per read increased to 1,000. The transcript expression was estimated using default parameters for all methods, except for the real data where strand-specific inference was enabled. Kallisto and Salmon were run without bias correction as it did not provide a clear advantage on the “Europe (excl. CEU)” pantranscriptome using the SRR1153470 reads (data not shown). RSEM was only run on the NA12878 personal sample-specific transcriptome and the “Europe (excl. CEU)” pantranscriptome, as it did not scale to the two largest pantranscriptomes.

rpvg was run using default parameters and with three different types of alignments inputs: the standard multipath alignments from vg mpmap and single-path alignments from vg map and vg mpmap. The vg mpmap single-path alignments were generated by finding the best scoring path in the multipath alignments using vg view. The fragment length distribution parameters estimated by vg mpmap were given as input to rpvg when using the vg map alignments. rpvg was run with a ploidy of 2 for all read sets, including CHM13. All HSTs with a haplotype probability below 0.8 were filtered from the rpvg output. The SRR1153470 and CHM13 read data was used to optimize the parameters of rpvg.

For the SRR1153470 and ENCSR000AED data, which are both NA12878 cell lines, we compared the quantified HSTs to the NA12878’s haplotypes from the 1000GP data. We considered an HST consistent with these haplotypes if it matched the sequence of one of the two possible NA12878 haplotype versions of the transcript. The haplotyping performance of each method was then estimated by comparing the number and fraction of quantified HSTs with positive expression that were consistent.

We used transcripts per million (TPM) to measure expression. For the simulated data we re-calculated the TPM value for all methods. The reason was that we wanted to ensure that there was no bias towards RSEM, which was used to estimate the expression profile employed by vg sim to parameterize the HST expression values. The TPM value depends on the effective transcript length, which is not calculated in the same manner for each method. Therefore, if this is not corrected, methods that estimate the effective transcript length more similarly to RSEM will have an advantage that does not depend on their ability to predict correct expression values. The true fragment length distribution parameters and the effective transcript length approach employed by rpvg (similar to Kallisto and Salmon) was used when re-calculating the TPM values.

The method’s ability to predict the correct expression value was evaluated using the simulated data for which the true expression is known. The true expression values were calculated from a table provided by vg sim, which indicates the transcript of origin for each read. The simulated TPM values were calculated in the same manner as described above. We used both Spearman correlation and mean absolute relative difference (MARD) to quantify concordance between estimated and true expression.

The CHM13 cell line is effectively haploid, so only a single HST is expected to exist for each transcript. We used this feature of the data to measure the haplotype inference performance of each method on the T2T CHM13 data. We defined each HST as either major or minor. Major HSTs were defined as the highest expressed haplotype for each transcript; the rest were defined as minor. The fraction of expression from minor HSTs is a lower bound on the fraction of incorrectly inferred transcript expression. Accordingly, we used the number of major and minor transcripts that each method predicted to be expressed to compare their haplotype inference performance.

To evaluate allele-specific expression (ASE) estimation, we converted the simulated and estimated HST expression values to allele-specific read counts for the NA12878 variants. These were calculated by dividing the expression values with the corresponding transcript length and multiplying by twice the read length (to account for paired-end sequencing). In addition, we inferred allele-specific read counts for the same NA12878 variants using the WASP [24] pipeline with STAR [2] alignments. Using both simulated and inferred allele-specific read counts, we next labeled heterozygotic variants with at least one read in the simulated data as showing significant ASE using a two-sided binomial test with p-values adjusted using the Benjamini-Hochberg procedure and a False Discovery Rate (FDR) *α* = 0.1. We took the hypothesis tests of the true simulated read counts to be the truth labels. The sets of labeled heterozygotic variants were lastly compared between simulation and methods to produce ASE true and false positive rates.

We assessed the rpvg’s robustness to admixture on real data using an indirect proxy. We applied the mpmap-rpvg pipeline to two samples from the same study: one of European American ancestry and the other of African American ancestry. We expect the African American individual to be more highly admixed. We then measured the proportion of marginal posterior expression assigned to the two most highly-expressed HSTs by summing over diplotypes. This is a lower bound on error, since for the majority of genes without a copy number alteration, there can only be two copies of the gene. We then computed the proportion of transcripts for which the two highest-expressed HSTs accounted for at least a given threshold proportion of the total expression. We repeated this analysis for different threshold values and stratified the results by minimum transcript expression.

### HLA pantranscriptome construction and typing

We evaluated the vg mpmap-rpvg pipeline’s ability to type HLA alleles. To start, we constructed a set of HLA haplotypes using gene allele sequences from the IPD-IMGT/HLA database (release 3.43.0) [41]. Many of the alleles in the database are partial and do not cover the corresponding entire gene, with a large fraction of them only covering the coding sequence or just the antigen recognition site (ARS) exons. Since haplotypes covering whole genes including introns are needed to construct a pantranscriptome using the vg rna pipeline, we first imputed the missing coding sequence for all the partial alleles using hlaseqlib [30]. This library contains a method which finds the closest complete allele for each partial allele and uses it to impute the missing sequence.

Next we used the reference to extend the alleles into full haplotypes. We padded each allele with 10,000 reference bases on both sides using the corresponding genes coding start and end location in the GENCODE v29 transcript annotation to ensure that the allele sequences would align to the correct genes. The padded HLA alleles were then aligned to a spliced pangenome graph using vg mpmap in long read, single-path mode. The resulting alignments were projected to the reference genome using vg surject and used to determine the location of splice-junctions in the allele sequences. The reference sequences of the corresponding introns were added to the allele to produce haplotype sequences covering the whole gene. The intron sequences were only added for junctions that were within 2 bases of an annotated splice-junction to ensure that genomic deletion were not mistakenly interpreted as splice-junctions.

These haplotypes were then used to create HLA pantranscriptomes. First, the haplotypes were mapped against the same spliced variation graph using vg mpmap in long read, single-path mode. The resulting alignments were used to update the graph with the variation in the haplotypes using vg augment. Using these haplotypes and vg rna, we created two HLA pantranscriptomes (see Supplementary Table 3): “HLA (main)” consisting of five of the main and most variable HLA genes and “HLA (10)” consisting of 10 HLA genes of which all had at least a 100 haplotypes. HLA Null alleles that have been shown not to be expressed were not included in the construction of the pantranscriptomes. In addition, transcript *HLA-B-258* were also not used as it was covering both *HLA-B* and *HLA-C*. This transcript have been removed in later versions of the GENCODE annotation. “HLA (main)” was built using the “1000GP (all, excl. CEU)” spliced variation graph, whereas “HLA (10)” was built using the “1000GP (all)” graph.

Using the “HLA (main)” pantranscriptome, we first optimized the default parameters of rpvg for HLA typing using six RNA-seq samples from the Geuvadis data set: NA07051, NA11832, NA11840, NA11930, NA12287 & NA12775 [43] (see Supplementary Table 4). All samples were from the CEU population, which were excluded in the variation graph construction. We compared the inferred alleles to the typing results from Gourraud et al. [44] and Abi-Rached et al. [45], both of which are available on the 1000GP homepage (https://www.internationalgenome.org/category/hla/). Similar to the quantification evaluation, HSTs with a haplotype probability below 0.8 were filtered before evaluation. An expressed HST were regarded as correct if its corresponding HLA allele matched one of the two studies. When evaluating diplotyping performance both HLA alleles needed to be correct and also match both ground truth alleles in the same study. The maximum number of standard deviations from the mean allowed for the fragment length, the maximum allowed score difference to the best alignment and the threshold for filtering alignments compared to the optimal score were adjusted to improve speed or typing accuracy (data not shown).

Next, we ran rpvg using the optimized parameters on two different sets of RNA-seq data: ten randomly selected CEU samples from Geuvadis that were not used in the optimization and 3 trios from the 1000GP sequenced as part of the Human Genome Structural Variation Consortium (HGSVC) [42] (see Supplementary Table 4). The Geuvadis and HGSVC data sets were run on the “HLA (main)” and “HLA (10)” pantranscriptomes, respectively. For the Geuvadis data, we used the same two studies as described above to determine typing accuracy, whereas for the HGSVC data trio concordance was used. For the Geuvadis data sets, typing accuracy for three different levels of HLA resolution were evaluated: 1 field, 2 field and G groups. G groups are defined as alleles that have identical nucleotide sequences across the ARS exons and were used to distinguish ambiguous alleles in the Gourraud et al. study based on Sanger sequencing of these exons.

### Variant genotyping and effect prediction

We used RNA-seq data from five randomly selected tissues from the same individual to estimate allele concordance across data sets. We also demonstrate the ability to genotype variants with potential effects on functional elements. All five data sets are available from the ENCODE project [35, 36] (see Supplementary Table 4). For each RNA-seq data set, all technical replicates were combined and the vg mpmap-rpvg pipeline was run using the “Whole” pantranscriptome (see Supplementary Table 3). The pipeline was run with default parameters except for rpvg, for which it was specified that the RNA-seq data was strand-specific.

The rpvg HST expression estimates were converted to variant allele-based expression values for the downstream analyses. To do this, all exonic variants were first annotated with the transcripts overlapping them using bcftools. These annotated variants were then used together with the original haplotypes to translate each HST to its corresponding set of variant alleles. Using this translation, we computed the expression of each variant allele as the sum of the expression over HSTs that contained the allele.

The haplotype probability values were similarly computed as a sum over the HSTs that contained the allele. However, for diplotypes where the alleles were called homozygous, only the probability of one of the haplotypes was added. This ensured that the corresponding alleles were only counted once per diplotype sample similar to the haplotype probabilities.

Using these results for each of the five tissues, we estimated the number of expressed variant alleles and the allele concordance between the tissues. For this analysis, we filtered all alleles with a probability below 0.8. An allele was defined to be expressed if it had nonzero expression in at least one tissue, and a variant was defined as expressed if at least one of its alleles, including the reference, had nonzero expression. Next, we estimated the consistency in whether an allele was expressed or not between tissues. To account for alternative expression and splicing across tissues, we only considered tissues for which the corresponding variant was expressed. An allele was then said to be concordant across tissues if it was either expressed or not expressed across all tissues that had the corresponding variant expressed (see Supplementary Figure 25). Variants that were only expressed in a single tissue were excluded for the concordance estimation. Since both alleles might not have been sampled in the sequencer by chance for lowly expressed exons, we repeated the analyses for different thresholds of variant expression. Finally, the homopolymer length of each variant was calculated by counting the maximum number of consecutive identical bases in each direction from the variant start site.

Next, we predicted the effect of the expressed variants on functional elements, such as transcript and protein sequences. This was done using the Ensembl Variant Effect Predictor (VEP) toolset [47]. VEP was run on all expressed variant alleles with a probability of at least 0.8 and a variant expression value of at least 5 TPM. The reason for the latter threshold was that we wanted to show the effects for exons with a non-negligible expression and also minimize genotyping error from allelic dropout. The GENCODE v29 transcript annotation was used for VEP with conversion to minimal variant representation enabled. In addition, multiple different columns of information were added to the output besides the default. Only predicted consequences with an impact rating of moderate or high were kept. For variants with both moderate and high impact consequences, only the high impact ones were shown.

### Demonstration of analyzing genomic imprinting

We obtained RNA-seq data sets from samples NA11832, NA11930, NA12775, and NA12889 from the Geuvadis data set [43] and ran them through the vg mpmap-rpvg pipeline (see Supplementary Table 4). Each sample had two accessions, which were combined into one data set. We also obtained and analyzed data from sample NA12878 from the ENCODE Project (ENCSR000AED replicate 1) [35, 36]. These samples are all unrelated. All parameters used were identical to those used in the real data evaluations of vg mpmap and only difference for rpvg was that Gibbs sampling was enabled. The Geuvadis samples were used to troubleshoot the analysis and identify potentially interesting genes to highlight in the demonstration. The analyses were then repeated on the ENCODE sample. This design reduces the risk of identifying noise as signal. Only the results final analysis are the ones reported in the Results section, but results were broadly consistent across all samples.

To confirm that the pipeline could detect previously known ASE, we looked for signatures of imprinting in the 20 genes with the most statistically significant parent-of-origin ASE in the study by Zink, et al. (from Supplementary Table 6) [21]. One of these genes, *RP11-69E11.4*, had since been removed from the GENCODE database, so we excluded it from the analysis. Zink’s, et al. study analyzed ASE on individual SNVs. To make our results comparable to theirs, we translated rpvg’s HST-based expression quantification into a corresponding variant allele-based expression quantification using the same approach as described above. The expression of each allele was computed as the sum of the expression of each HST that contained the allele.

We decided to highlight the haplotype-specific expression of the NAA60 gene in depth because it consistently showed monoallelic expression for both haplotypes across different isoforms in the initial exploratory data sets. To identify the haplotype of origin for different HSTs, we compared the variants associated with each HST (using the table from vg rna) to the sample’s haplotypes from the 1000GP VCF. Equal-tailed credible intervals were approximated using rpvg’s Gibbs sampling method.

### Code availability

The source code for vg and rpvg is publicly available at https://github.com/vgteam/vg [87] and https://github.com/jonassibbesen/rpvg [88] respectively. Both tools are licensed under the MIT License. A full list of the versions of all computational tools used is available in Supplementary Table 6. All bash scripts with exact command-lines used to generate the results are available at https://github.com/jonassibbesen/vgrna-project-paper [89]. This repository includes references to Docker containers and log files from the analyses. All custom C++, Python, and R scripts used for analysis and plotting are available at both the above repository and https://github.com/jonassibbesen/vgrna-project-scripts [90].

### Data availability

All data used in this study are available at https://github.com/jonassibbesen/vgrna-project-paper. Data that are available from public repositories are provided as web links only. Accession numbers are included when relevant, and accession numbers for sequencing data are also listed in Supplementary Table 4. The repository also includes all spliced pangenome graphs and pantranscriptome haplotype-specific transcript sets, which may be freely used in other projects.

