## Supplementary material for "Haplotype-aware pantranscriptome analyses using spliced pangenome graphs"

### Supplementary information for haplotype-aware pantranscriptome analyses using spliced pangenome graphs

#### Supplementary Figures

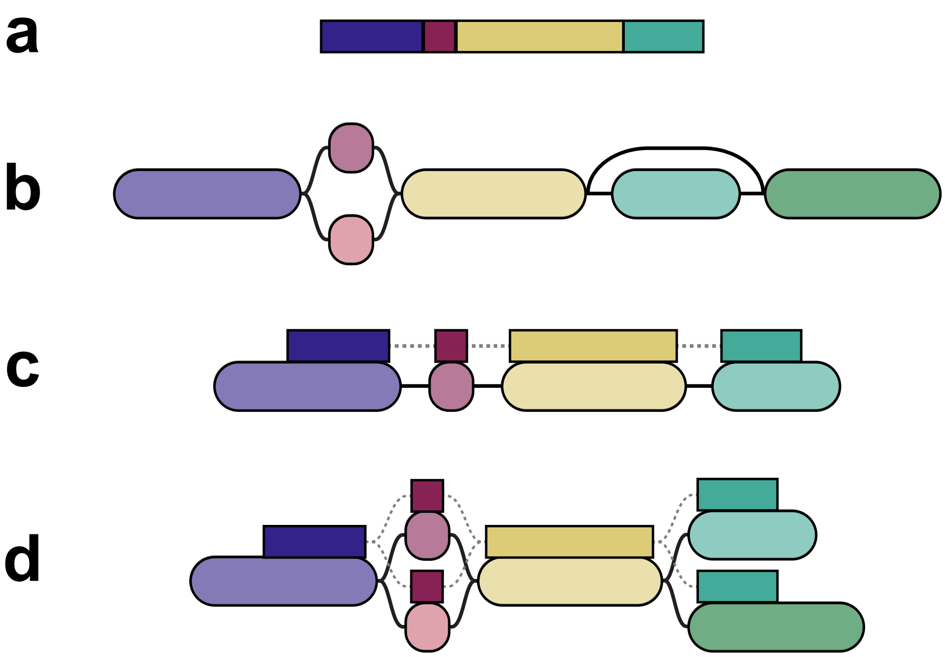


**Supplementary Figure 1: Diagram of a multipath alignment**
A diagrammatic comparison between the multipath alignment output of VG MPMAP and the single-path alignment output of other graph aligners (such as VG MAP). **a** A read and **b** a sequence graph, which have been colored to indicate which parts of the read could plausibly align to which parts of the graph. **c** A single-path alignment. The read sequence is aligned to one path from the graph. **d** A multipath alignment. The alignment can split and rejoin to express the alignment uncertainty to different paths in the graph.

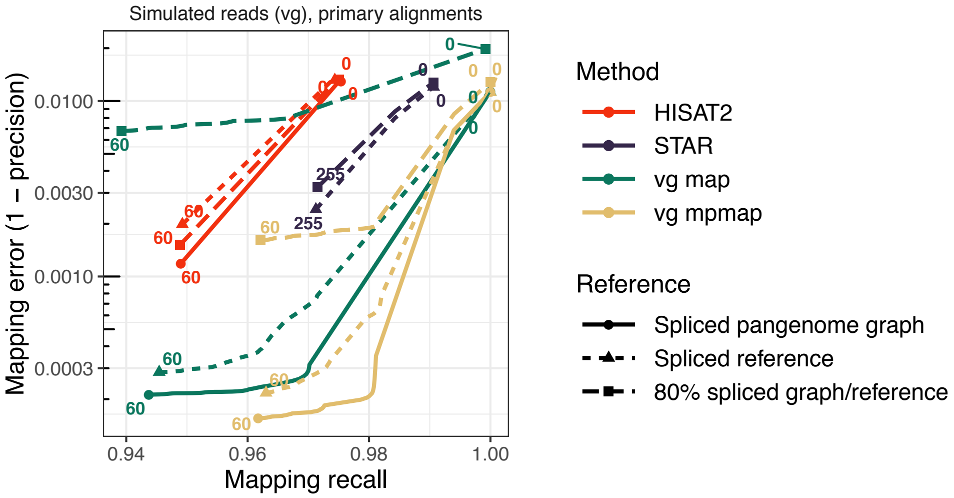


**Supplementary Figure 2: Mapping benchmark for primary alignments using RNA-seq data from NA12878**
Mapping error and recall for vg mpmap and three other methods using simulated Illumina data ("vg sim (ENC, uniform)" in Supplementary Table 5). Colored numbers indicate different mapping quality thresholds. Reads are considered correctly mapped if their primary alignments covers 90% of the true reference sequence alignment. Solid lines show results for spliced pangenome graph ("1000GP (all, excl. CEU)" in Supplementary Table 2), short dashed lines for spliced reference, and long dashed lines for a spliced reference (STAR) or spliced pangenome graph (all others) made with a random subset containing 80% of the transcripts.

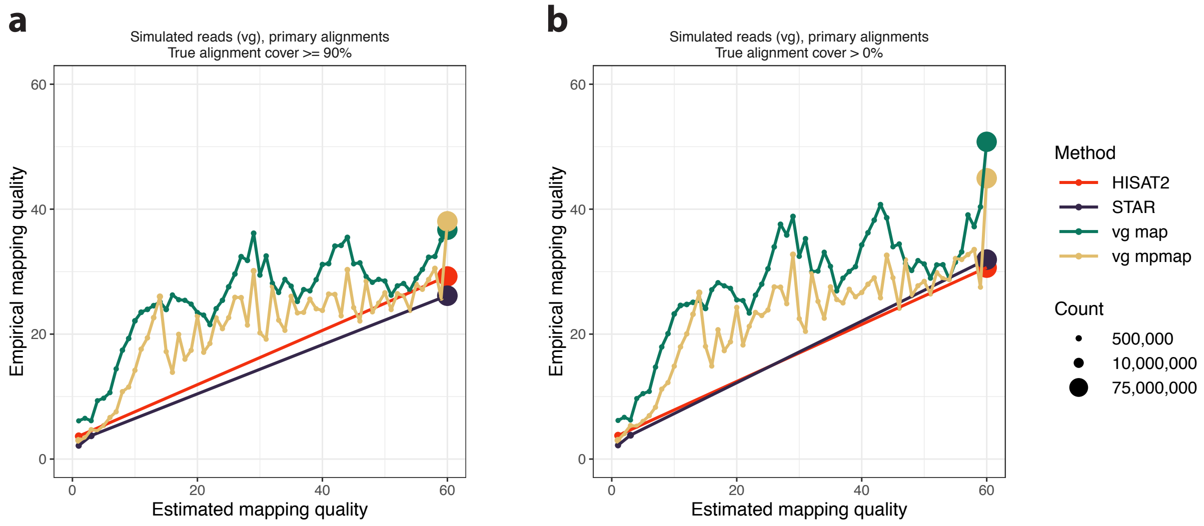


**Supplementary Figure 3: Benchmark of mapping quality calibration using RNA-seq data from NA12878**
Quantile-quantile (QQ) plot of the reported probability of mapping error (using the probabilistic interpretation of mapping quality) and the observed rate of mapping error for vg mpmap and three other methods. The reads used are simulated Illumina data ("vg sim (ENC, uniform)" in Supplementary Table 5). Correctness is determined by **a** 90% cover of the true alignment, and **b** any overlap with the true alignment. Results are shown using a spliced reference for STAR and for the other methods a spliced pangenome graph ("1000GP (all, excl. CEU)" in Supplementary Table 2).

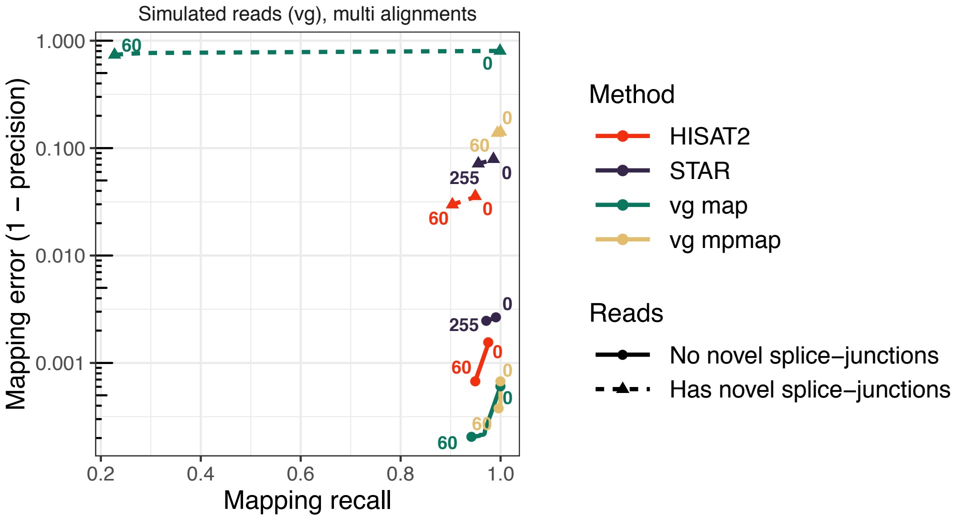


**Supplementary Figure 4: Mapping benchmark stratified by novel splice-junctions using RNA-seq data from NA12878**

Mapping error and recall for vg mpmap and three other methods using simulated Illumina data ("vg sim (ENC, uniform)" in Supplementary Table 5). Colored numbers indicate different mapping quality thresholds. Reads are considered correctly mapped if one of their multi-alignments covers 90% of the true reference sequence alignment. Solid and dashed lines show the results for reads containing no novel splice-junctions and at least one novel splice-junction, respectively. Results are shown using a spliced reference (STAR) or spliced pangenome graph (all others) made with a random subset containing 80% of the transcripts.

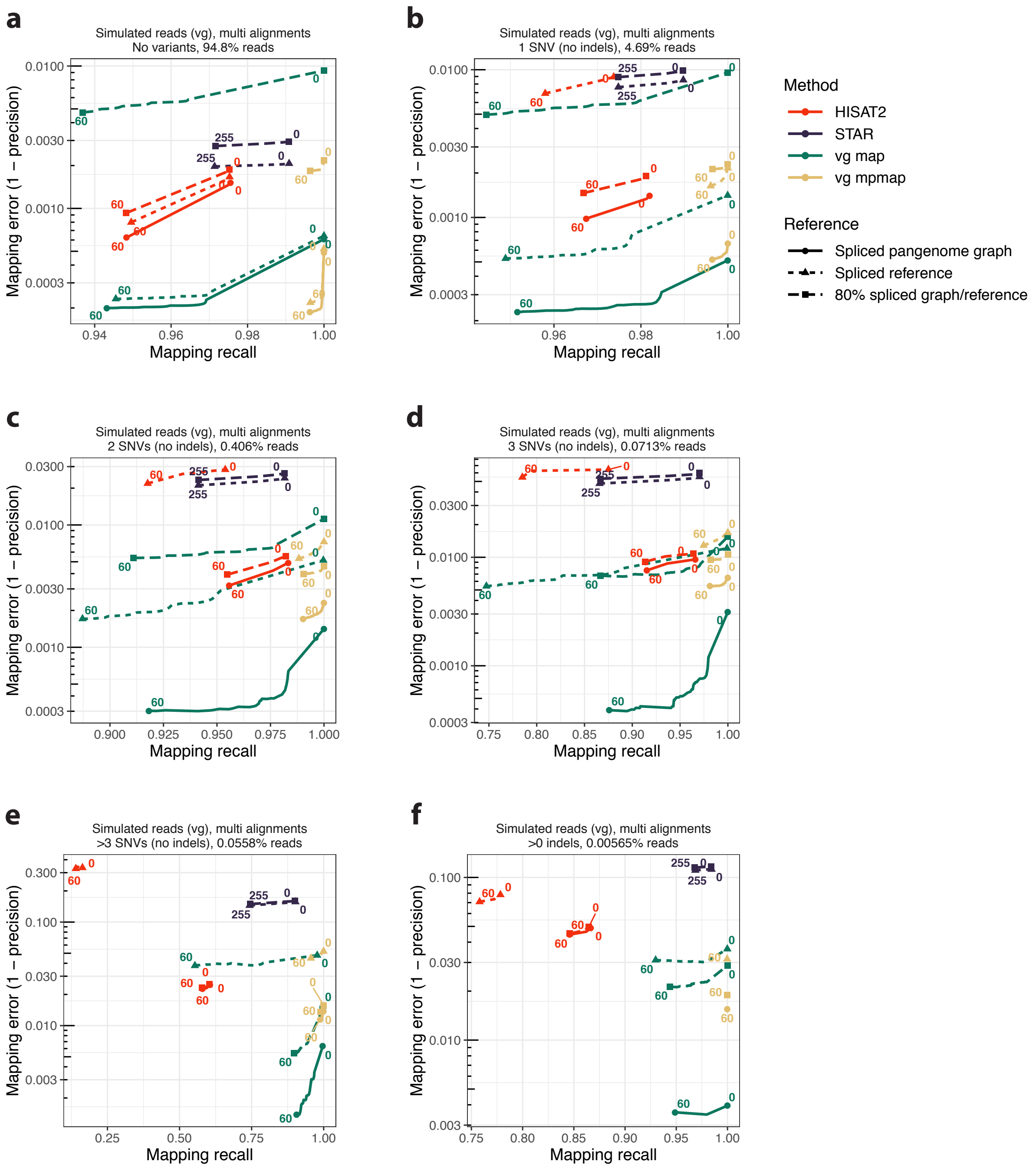


**Supplementary Figure 5: Mapping benchmark stratified by non-reference variants using RNA-seq data from NA12878**

Mapping error and recall for VG MPMAP and three other methods using simulated Illumina data ("vg sim (ENC, uniform)" in Supplementary Table 5). Colored numbers indicate different mapping quality thresholds. Reads are considered correctly mapped if one of their multi-alignments covers 90% of the true reference sequence alignment. Solid lines show results for spliced pangenome graph ("1000GP (all, excl. CEU)" in Supplementary Table 2), short dashed lines for spliced reference, and long dashed lines for a spliced reference (STAR) or spliced pangenome graph (all others) made with a random subset containing 80% of the transcripts. Reads are stratified into those that **a** contain no variants, **b** contain no insertions or deletions (indels) and one single nucleotide variant (SNV), **c** contain no indels and two SNVs, **d** contain no indels and three SNVs, **e** contain no indels and more than three SNVs, and **f** contain any indels.

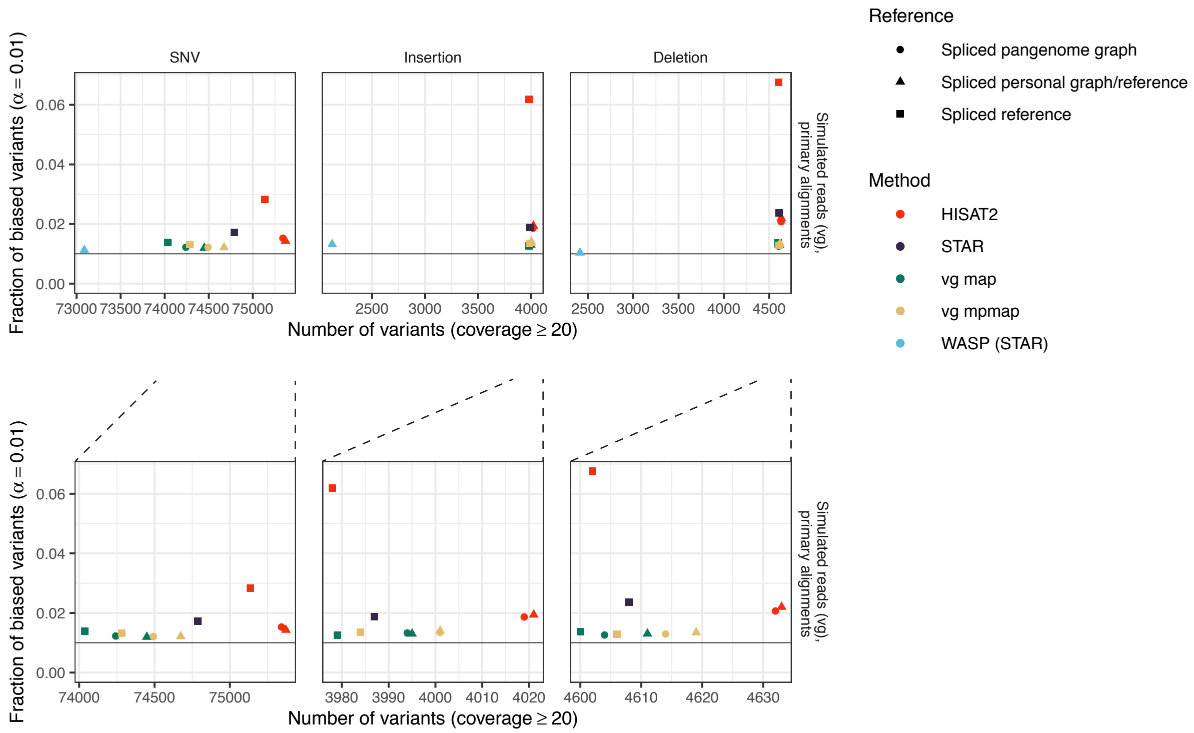


**Supplementary Figure 6: Allelic bias benchmark using RNA-seq data from NA12878**
Allelic mapping bias for VG MPMAP and four other methods using simulated Illumina RNA-seq reads ("vg sim (ENC, uniform)" in Supplementary Table 5), which were simulated without allelic bias. STAR was used as the aligner for the WASP pipeline. Circles and squares show the results using a spliced pangenome graph ("1000GP (all, excl. CEU)" in Supplementary Table 2) and spliced reference, respectively. Triangles shows the results using a spliced personal reference or graph containing only the 1000 Genomes Project (1000GP) NA12878 variants ("1000GP (NA12878)" in Supplementary Table 2). The WASP (STAR) pipeline were provided the 1000GP NA12878 haplotypes as input. The number of variant sites with coverage at least 20 is plotted against the observed rate of false positive hypothesis tests of allelic skew (two-sided binomial test, $\alpha = 0.01$). Coverage was calculated from primary alignments with a mapping quality value of at least 30. The bottom row shows a zoomed view without WASP (STAR).

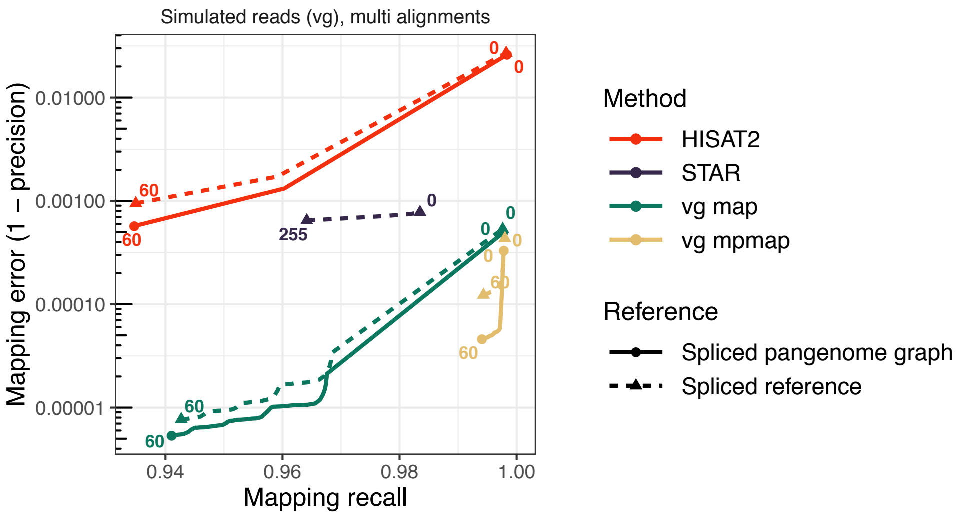


**Supplementary Figure 7: Graph-based mapping benchmark using RNA-seq data from NA12878**

Mapping error and recall for VG MPMAP and three other methods using simulated Illumina data ("vg sim (ENC, uniform)" in Supplementary Table 5). Colored numbers indicate different mapping quality thresholds. An alignment is considered correct if its start position is within 100 bases from the start position of the true alignment measured using any labeled transcript path in the graph or the linear reference sequence. Solid and dashed lines show the results using a spliced pangenome graph ("1000GP (all, excl. CEU)" in Supplementary Table 2) and spliced reference, respectively.

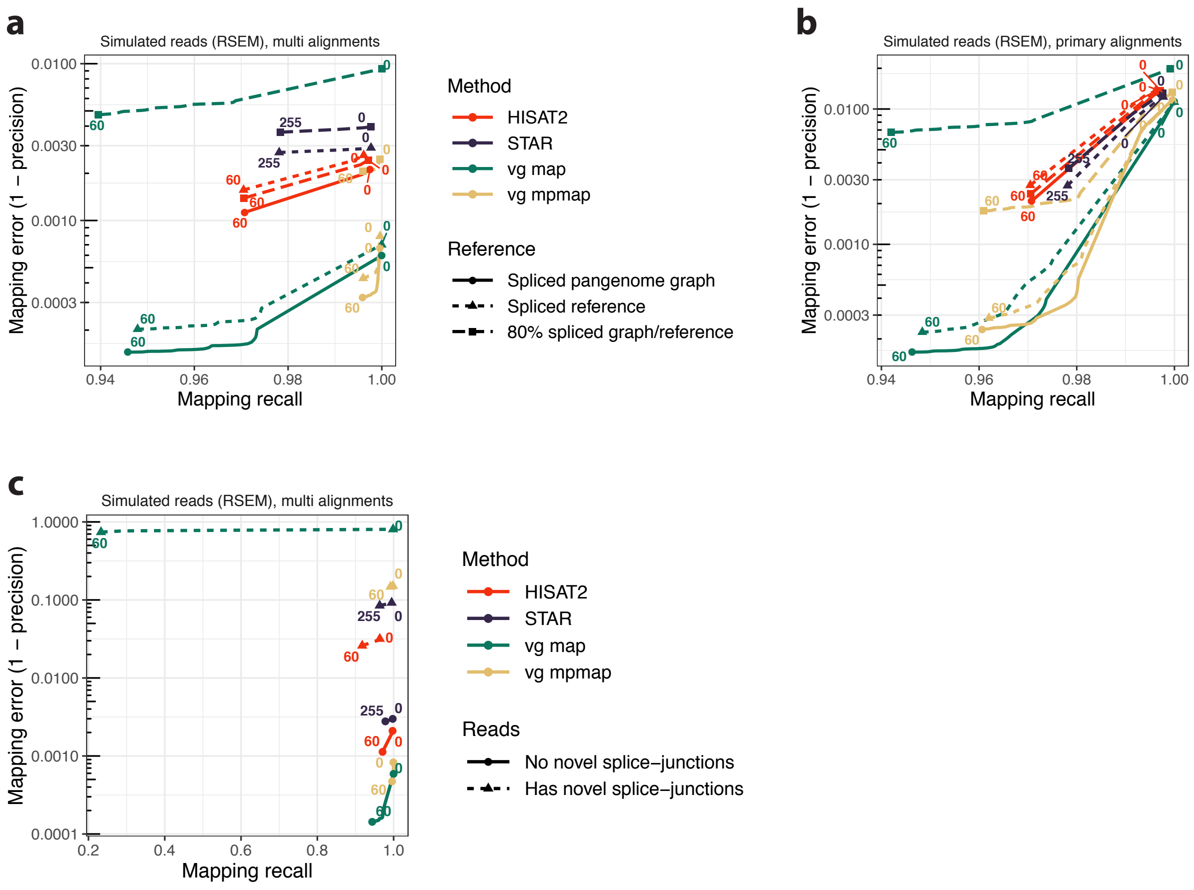


**Supplementary Figure 8: Mapping benchmark using RSEM-simulated RNA-seq data from NA12878**

Mapping error and recall for VG MPMAP and three other methods using simulated Illumina data ("RSEM (ENC, uniform)" in Supplementary Table 5). Colored numbers indicate different mapping quality thresholds. Reads are considered correctly mapped if one of their multi-alignments (**a**,**c**) or their primary alignments (**b**) cover 90% of the true reference sequence alignment. **a**,**b** Solid lines show results for spliced pangenome graph ("1000GP (all, excl. CEU)" in Supplementary Table 2), short dashed lines for spliced reference, and long dashed lines for a spliced reference (STAR) or spliced pangenome graph (all others) made with a random subset containing 80% of the transcripts. **c** Shows reads stratified into those that contain no (solid lines) and at least one (dashed lines) novel splice-junction. Results are shown using a reference/graph made with a random subset containing 80% of the transcripts.

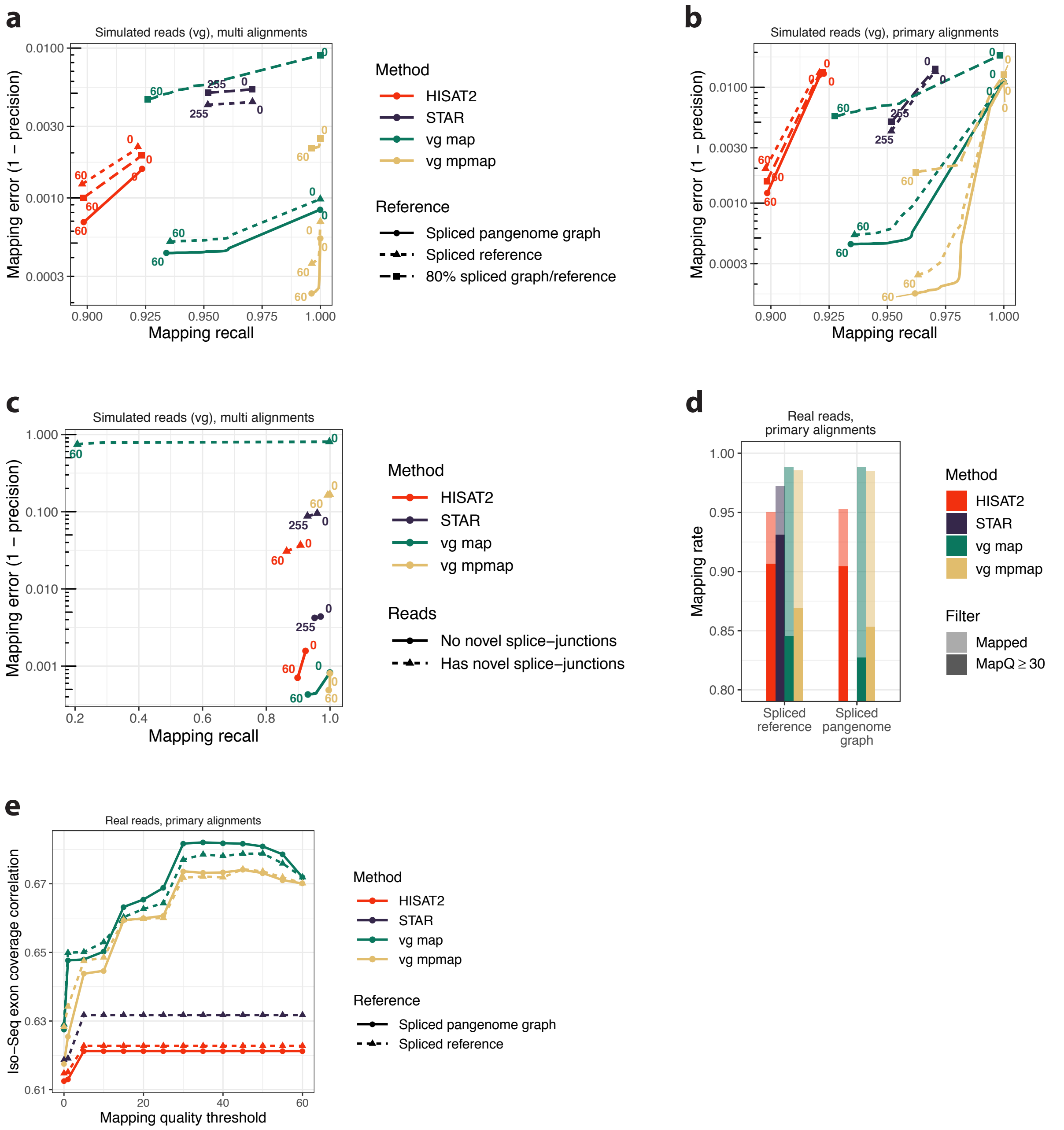


**Supplementary Figure 9: Mapping benchmark using RNA-seq training data from NA12878** RNA-seq mapping results comparing VG MPMAP and three other methods using the simulated and real Illumina data that was used in the optimization of VG MAP and VG MPMAP ("vg sim (SRR, uniform)" and "SRR1153470" in Supplementary Table 5 and 4, respectively). **a-c** Mapping error and recall for different mapping quality thresholds (colored numbers) using simulated data. Reads are considered correctly mapped if one of their multi-alignments (**a**,**c**) or their primary alignments (**b**) cover 90% of the true reference sequence alignment. **a**,**b**,**e** Solid lines show results for spliced pangenome graph ("1000GP (all, excl. CEU)" in Supplementary Table 2), short dashed lines for spliced reference, and long dashed lines for a spliced reference (STAR) or spliced pangenome graph (all others) made with a random subset containing 80% of the transcripts. **c** Shows reads stratified into those that contain no (solid lines) and at least one (dashed lines) novel splice-junction. Results are shown using a reference/graph made with a random subset containing 80% of the transcripts. **d** Mapping rate using real data. The solid bars show the mapping rate using a mapping quality threshold of 30. **e** Pearson correlation between Illumina and Iso-Seq exon coverage using real data as a function of mapping quality threshold. Exons were defined by the Iso-Seq alignments and each exon's coverage was normalized by its length.

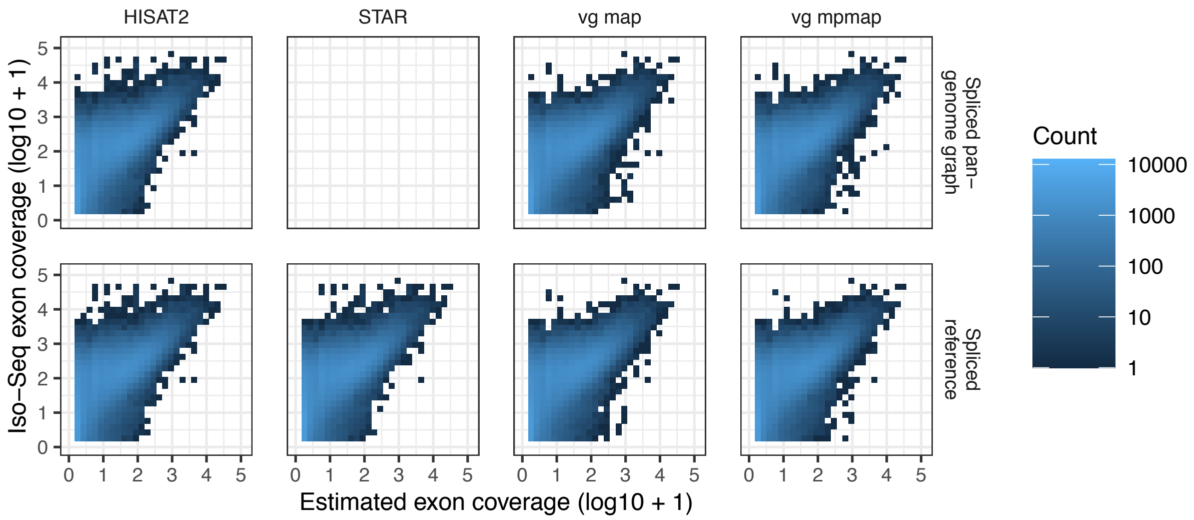


**Supplementary Figure 10: Exon coverage comparison using Iso-Seq and Illumina RNA-seq data from NA12878**
Comparison of exon-level coverage estimated using Iso-Seq alignments ("ENCSR706ANY" in Supplementary Table 4) and real Illumina RNA-seq reads ("ENCSR000AED" in Supplementary Table 4) mapped by VG MPMAP and three other methods. Exons were defined by the Iso-Seq alignments and each exon's coverage was normalized by its length. Top and bottom rows show the results using a spliced pangenome graph ("1000GP (all, excl. CEU)" in Supplementary Table 2) and spliced reference, respectively.

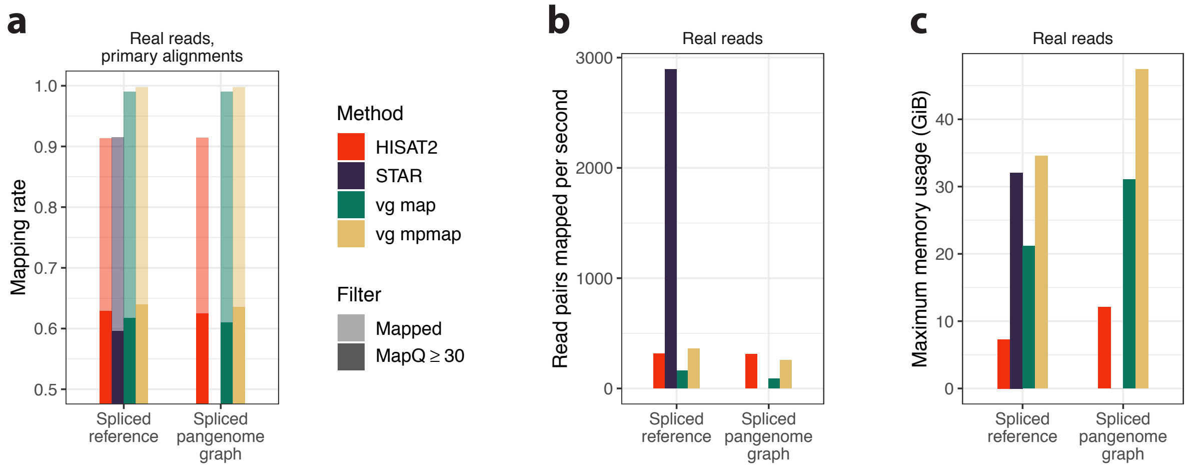


**Supplementary Figure 11: Mapping benchmark using RNA-seq training data from CHM13**
RNA-seq mapping results comparing VG MPMAP against three other methods using real Illumina data that was used in the optimization of VG MPMAP ("CHM13" in Supplementary Table 4). Reads are mapped both to a spliced reference and a spliced pangenome graph ("1000GP (all)" in Supplementary Table 2). **a** Mapping rate. The solid bars show the mapping rate using a mapping quality threshold of 30. **b** Number of read pairs mapped per second per thread. The mapping times were measured using 16 threads on a AWS m5.4xlarge instance. **c** Maximum memory usage for mapping in gigabytes.

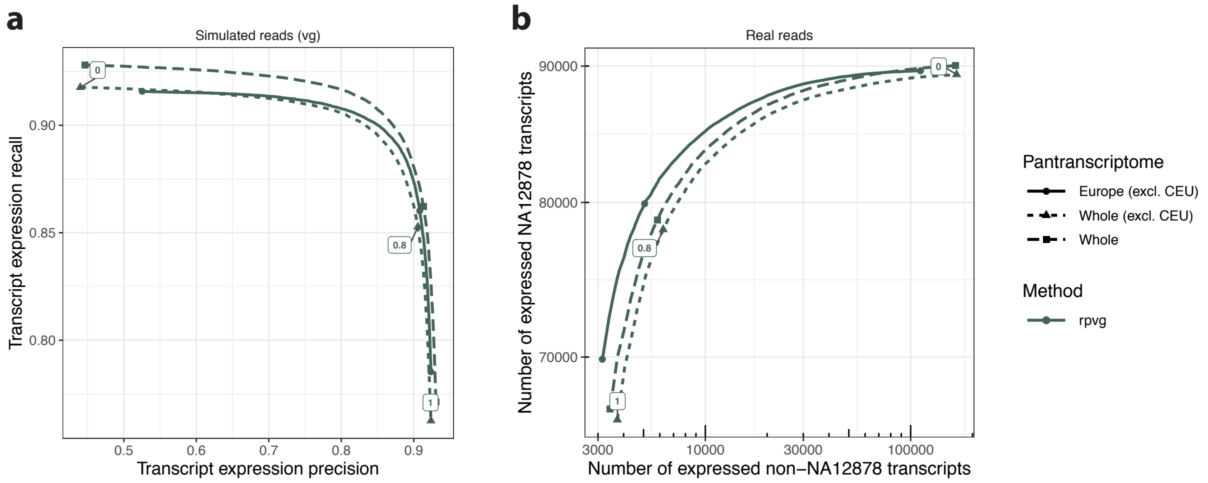


**Supplementary Figure 12: Haplotype probability evaluation using RNA-seq data from NA12878**

Evaluation of the haplotype probability estimated for each haplotype-specific transcript (HST) by RPVG using simulated and real Illumina data ("vg sim (ENC, RSEM)" and "ENCSR000AED" in Supplementary Table 5 and 4, respectively). Solid lines with circles are results using a pantranscriptome generated from 1000 Genomes Project (1000GP) European haplotypes excluding the CEU population. Dashed lines with triangles and squares are results using a pantranscriptome generated from all 1000GP haplotypes without and with the CEU population, respectively (Supplementary Table 3). **a** Recall and precision for whether a transcript is correctly assigned nonzero expression for different haplotype probability thresholds (colored numbers for "Whole (excl. CEU)" pantranscriptome) using simulated data. **b** Number of expressed transcripts from NA12878 haplotypes shown against the number from non-NA12878 haplotypes for different haplotype probability thresholds (colored numbers) using real data. All HSTs with a haplotype probability below 0.8 were filtered from the RPVG output for the HST quantification benchmark.

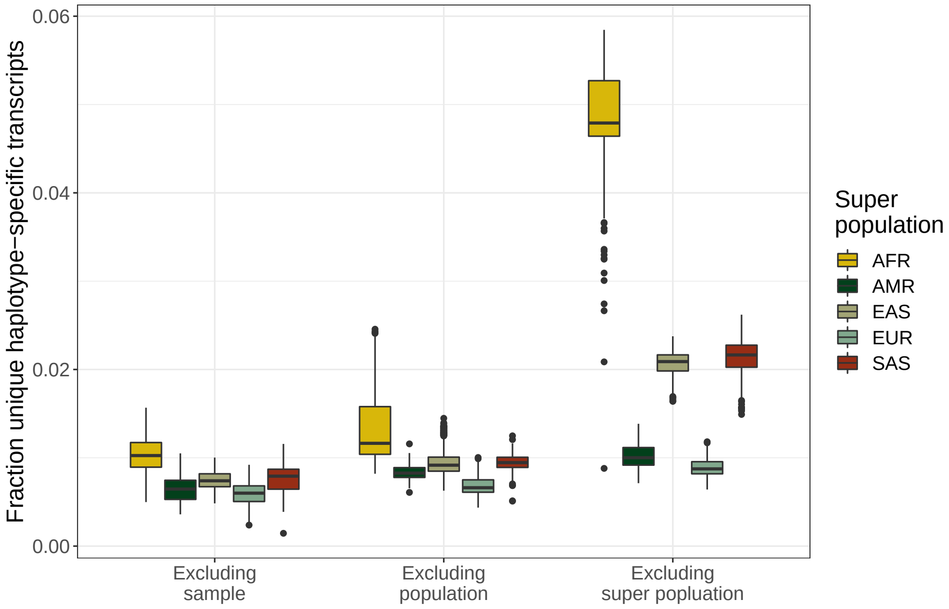


**Supplementary Figure 13: Haplotype-specific transcript uniqueness in a 1000 Genomes Project pantranscriptome**
The fraction of HSTs that are unique to each sample in the 1000 Genomes Project (1000GP) when compared to different subsets of samples in the 1000GP. Left box plots show the fraction unique when comparing to all other samples, middle box plots show the fraction unique when comparing to all other samples excluding the samples' population, and right box plots show the fraction unique when comparing to all other samples excluding the samples' super population. AFR: African, AMR: Admixed American, EAS: East Asian, EUR: European, SAS: South Asian.

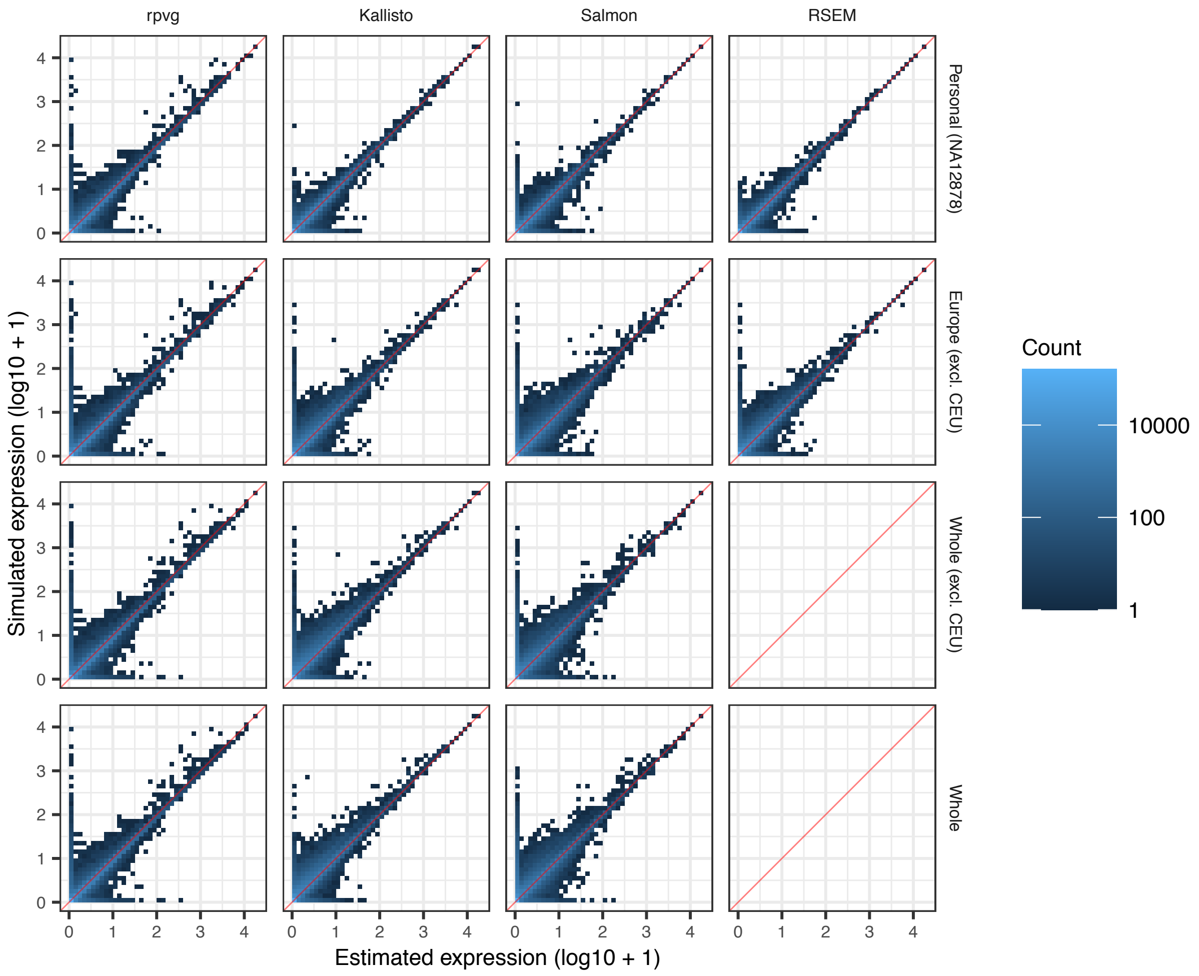


**Supplementary Figure 14: Haplotype-specific transcript expression comparison using RNA-seq data from NA12878**
Comparison of estimated and simulated NA12878 haplotype-specific transcript (HST) expression values (in transcripts per million (TPM)) for RPVG and three other methods using simulated Illumina data ("vg sim (ENC, RSEM)" in Supplementary Table 5). "Personal (NA12878)" is a personal sample-specific transcriptome generated from 1000 Genomes Project (1000GP) NA12878 haplotypes. "Europe (excl. CEU)" is a pantranscriptome generated from European 1000GP haplotypes excluding the CEU population. "Whole (excl. CEU)" and "Whole" are pantranscriptomes generated from all 1000GP haplotypes without and with the CEU population, respectively (Supplementary Table 3).

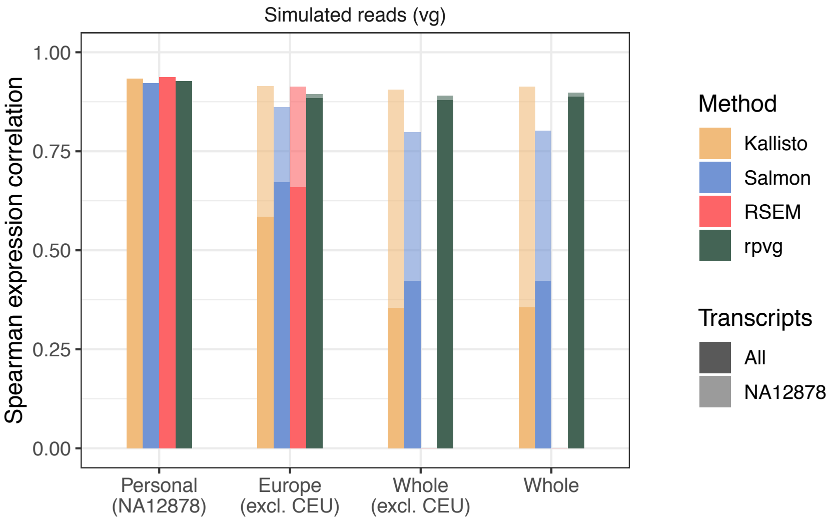


**Supplementary Figure 15: Haplotype-specific transcript expression correlation benchmark using RNA-seq data from NA12878**
Haplotype-specific transcript (HST) quantification results comparing RPVG and three other methods using simulated Illumina data ("vg sim (ENC, RSEM)" in Supplementary Table 5). Shows Spearman correlation between simulated and estimated expression (in transcripts per million (TPM)) for different pantranscriptomes. Correlation was calculated using either all HSTs in the pantranscriptome (solid bars) or using only the NA12878 HSTs (shaded bars). "Personal (NA12878)" is a personal sample-specific transcriptome generated from 1000 Genomes Project (1000GP) NA12878 haplotypes. "Europe (excl. CEU)" is a pantranscriptome generated from European 1000GP haplotypes excluding the CEU population. "Whole (excl. CEU)" and "Whole" are pantranscriptomes generated from all 1000GP haplotypes without and with the CEU population, respectively (Supplementary Table 3).

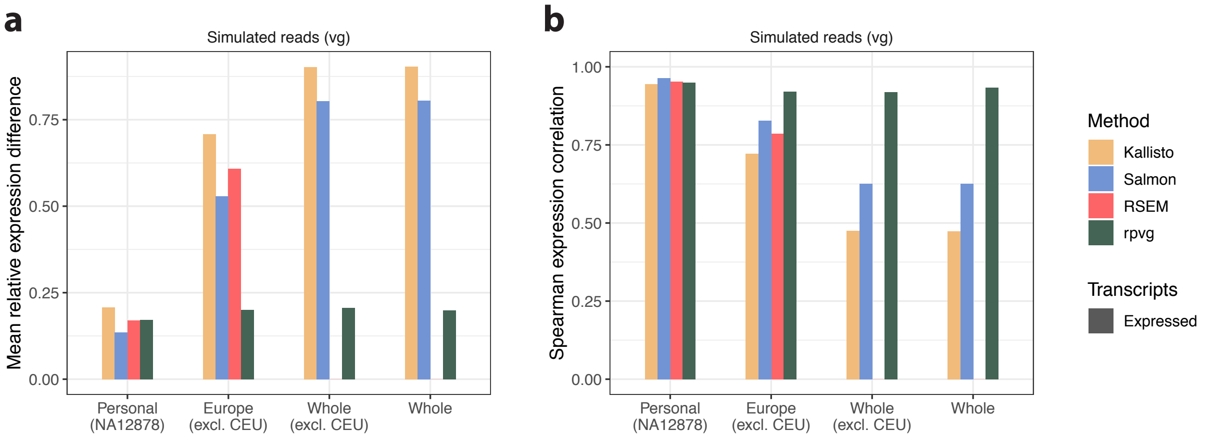


**Supplementary Figure 16: Expression benchmark of expressed haplotype-specific transcripts using RNA-seq data from NA12878**
Haplotype-specific transcript (HST) quantification results comparing RPVG and three other methods using simulated Illumina data ("vg sim (ENC, RSEM)" in Supplementary Table 5). "Personal (NA12878)" is a personal sample-specific transcriptome generated from 1000 Genomes Project (1000GP) NA12878 haplotypes. "Europe (excl. CEU)" is a pantranscriptome generated from European 1000GP haplotypes excluding the CEU population. "Whole (excl. CEU)" and "Whole" are pantranscriptomes generated from all 1000GP haplotypes without and with the CEU population, respectively (Supplementary Table 3). **a** Mean absolute relative expression difference (MARD) and **b** Spearman correlation between simulated and estimated expression (in transcripts per million (TPM)) for different pantranscriptomes using simulated data. MARD and correlation were calculated using only HSTs estimated to be expressed by each method.

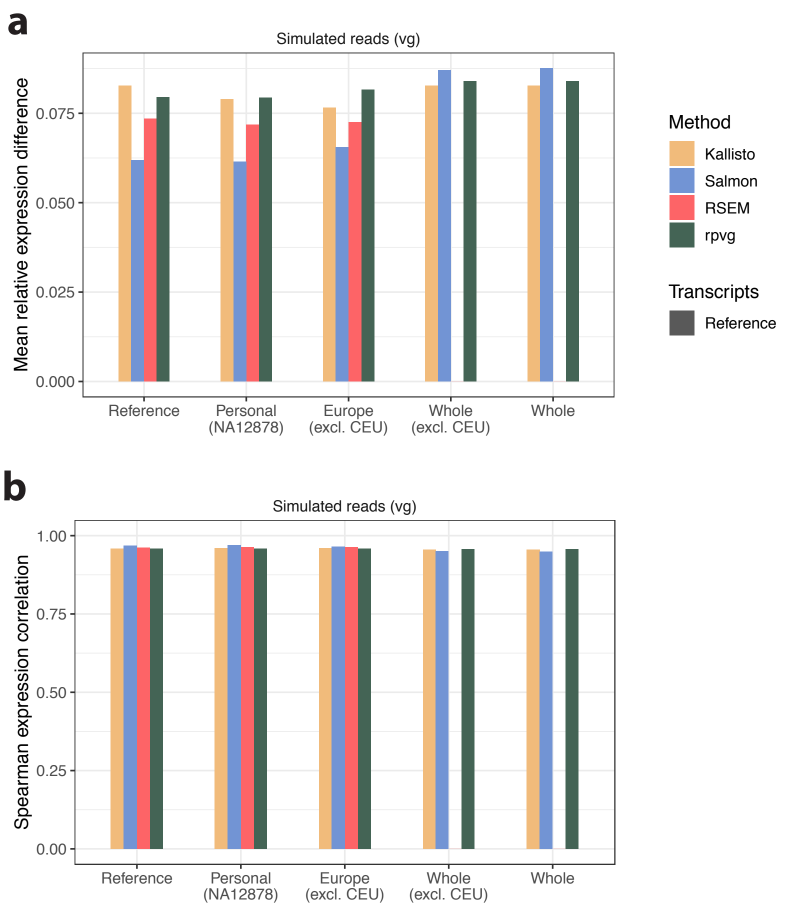


**Supplementary Figure 17: Transcript expression benchmark using RNA-seq data from NA12878**

Transcript quantification results comparing RPVG and three other methods using simulated Illumina data ("vg sim (ENC, RSEM)" in Supplementary Table 5). "Reference" is the regular reference-based transcript annotation. "Personal (NA12878)" is a personal sample-specific transcriptome generated from 1000 Genomes Project (1000GP) NA12878 haplotypes. "Europe (excl. CEU)" is a pantranscriptome generated from European 1000GP haplotypes excluding the CEU population. "Whole (excl. CEU)" and "Whole" are pantranscriptomes generated from all 1000GP haplotypes without and with the CEU population, respectively (Supplementary Table 3). **a** Mean absolute relative expression difference (MARD) and **b** Spearman correlation between simulated and estimated expression (in transcripts per million (TPM)) for the reference set and different pantranscriptomes using simulated data. For the pantranscriptomes, transcript expression were calculated by adding together the estimated expression values of all haplotype-specific transcripts from the same transcript.

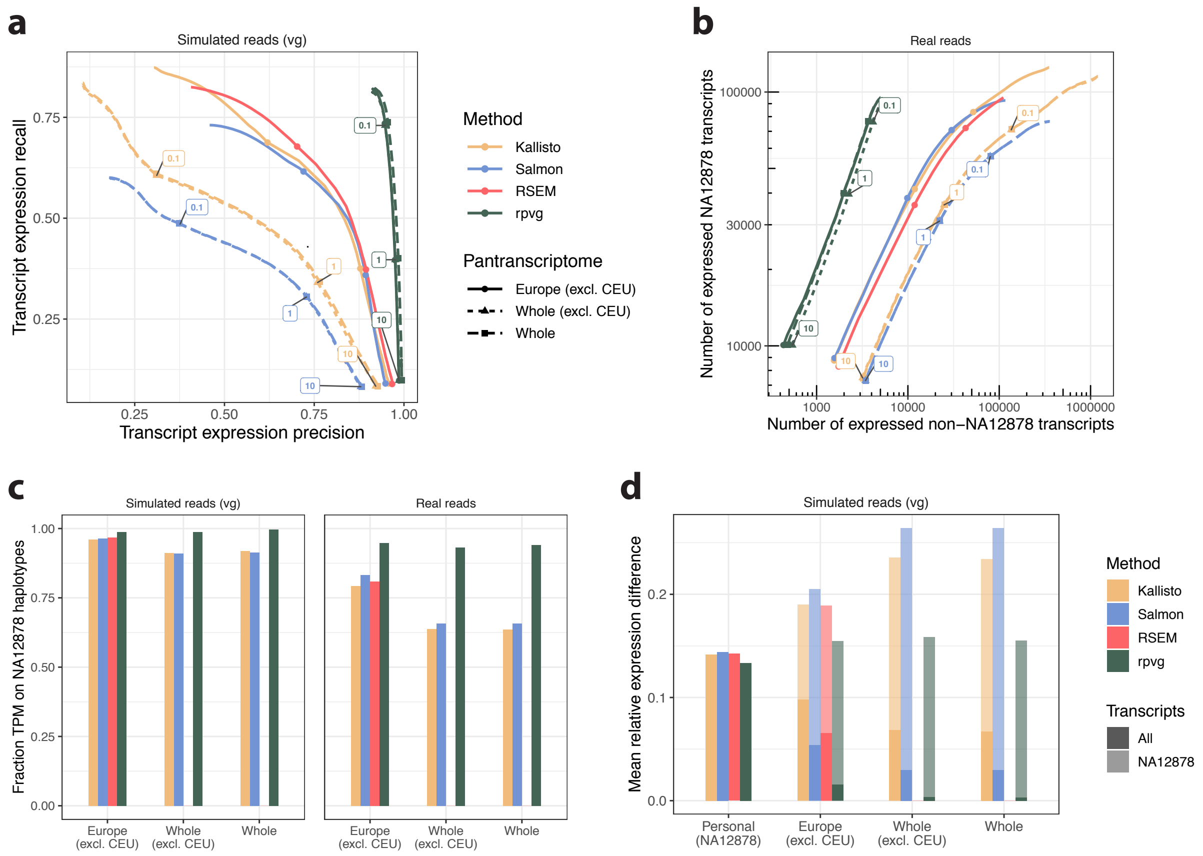


**Supplementary Figure 18: Haplotype-specific transcript quantification benchmark using RNA-seq training data from NA12878**

Haplotype-specific transcript (HST) quantification results comparing RPVG against three other methods using simulated and real Illumina data that was used in the optimization of RPVG ("vg sim (SRR, RSEM)" and "SRR1153470" in Supplementary Table 5 and 4, respectively). Solid lines with circles are results using a pantranscriptome generated from 1000 Genomes Project (1000GP) European haplotypes excluding the CEU population. Dashed lines with triangles and squares are results using a pantranscriptome generated from all 1000GP haplotypes without and with the CEU population, respectively (Supplementary Table 3). **a** Recall and precision of whether a transcript is correctly assigned nonzero expression for different expression value thresholds (colored numbers for "Whole (excl. CEU)" pantranscriptome) using simulated data. Expression is measured in transcripts per million (TPM). **b** Number of expressed transcripts from NA12878 haplotypes shown against the number from non-NA12878 haplotypes for different expression value thresholds (colored numbers) using real data. **c** Fraction of transcript expression (in TPM) assigned to NA12878 haplotypes for different pantranscriptomes using simulated (left) and real (right) data. **d** Mean absolute relative expression difference (MARD) between simulated and estimated expression (in TPM) for different pantranscriptomes using simulated data. MARD was calculated using either all HSTs in the pantranscriptome (solid bars) or using only the NA12878 HSTs (shaded bars). "Personal (NA12878)" is a personal sample-specific transcriptome generated from 1000GP NA12878 haplotypes (Supplementary Table 3).

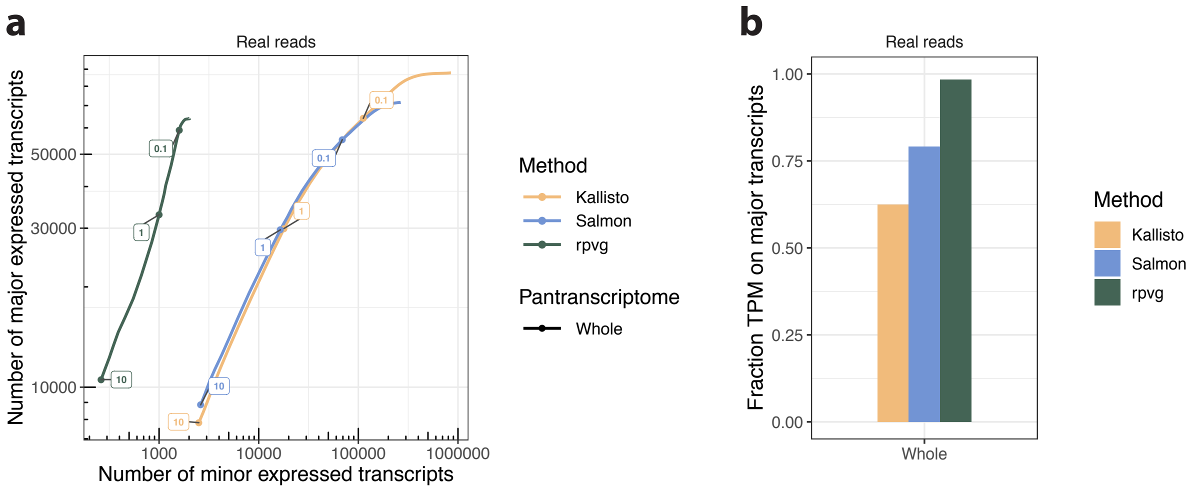


**Supplementary Figure 19: Haplotype-specific transcript quantification benchmark using RNA-seq training data from CHM13**

Haplotype-specific transcript (HST) quantification results comparing RPVG against two other methods using real Illumina data that was used in the optimization of RPVG ("CHM13" in Supplementary Table 4). A pantranscriptome generated from all 1000 Genomes Project haplotypes were used for the benchmark ("Whole" in Supplementary Table 3). Each HST is either classified as major or minor. Major HSTs are defined as the highest expressed haplotype for each transcript; the rest are defined as minor. As CHM13 is effectively haploid, the fraction of expression from minor HSTs is a lower bound on the fraction of incorrectly inferred transcript expression. **a** Number of major expressed transcripts against the number of minor expressed for different expression value thresholds (colored numbers). Expression is measured in transcripts per million (TPM). **b** Fraction of transcript expression (in TPM) assigned to major transcripts for different methods.

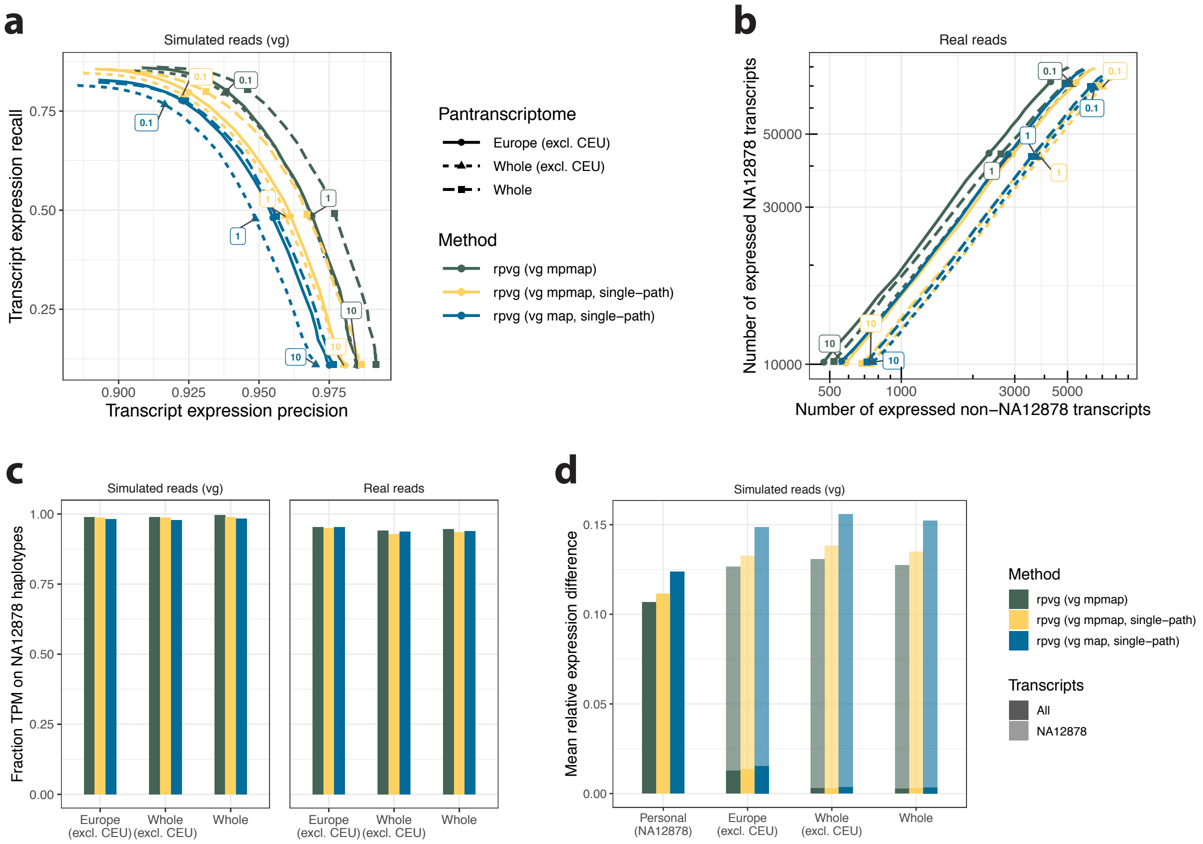


**Supplementary Figure 20: Multipath alignment benchmark using RNA-seq data from NA12878**

Haplotype-specific transcript (HST) quantification results comparing RPVG with single-path and multipath alignments from VG MPMAP and VG MAP as input using simulated and real Illumina data ("vg sim (ENC, RSEM)" and "ENCSR000AED" in Supplementary Table 5 and 4, respectively). Solid lines with circles are results using a pantranscriptome generated from 1000 Genomes Project (1000GP) European haplotypes excluding the CEU population. Dashed lines with triangles and squares are results using a pantranscriptome generated from all 1000GP haplotypes without and with the CEU population, respectively (Supplementary Table 3). The VG MPMAP single-path alignments were created by finding the best scoring path in each multipath alignment. **a** Recall and precision of whether a transcript is correctly assigned nonzero expression for different expression value thresholds (colored numbers for "Whole (excl. CEU)" pantranscriptome) using simulated data. Expression is measured in transcripts per million (TPM). **b** Number of expressed transcripts from NA12878 haplotypes shown against the number from non-NA12878 haplotypes for different expression value thresholds (colored numbers) using real data. **c** Fraction of transcript expression (in TPM) assigned to NA12878 haplotypes for different pantranscriptomes using simulated (left) and real (right) data. **d** Mean absolute relative expression difference (MARD) between simulated and estimated expression (in TPM) for different pantranscriptomes using simulated data. MARD was calculated using either all HSTs in the pantranscriptome (solid bars) or using only the NA12878 HSTs (shaded bars). "Personal (NA12878)" is a personal sample-specific transcriptome generated from 1000GP NA12878 haplotypes (Supplementary Table 3).

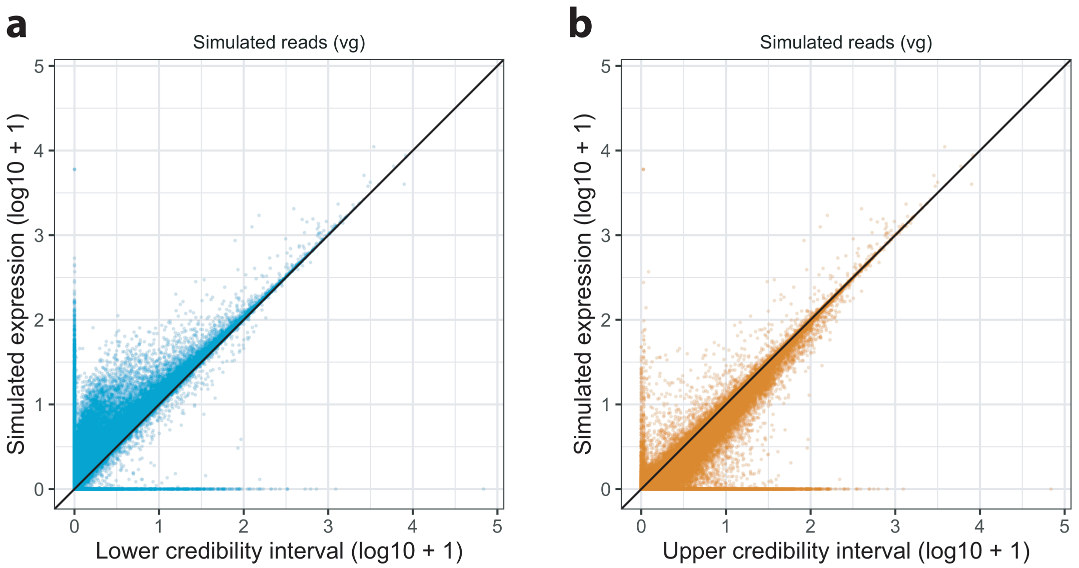


**Supplementary Figure 21: Gibbs sampling evaluation using RNA-seq data from NA12878** Evaluation of haplotype-specific transcript expression quantification uncertainty estimated by Gibbs sampling in RPVG using simulated data ("vg sim (ENC, RSEM)" in Supplementary Table 5). A pantranscriptome generated from all 1000GP haplotypes without the CEU population was used for the evaluation ("Whole (excl. CEU)" in Supplementary Table 3). **a** Lower and **b** upper bounds of equal-tailed 90% credibility intervals shown against simulated expression values. The credibility intervals were estimated from a 1000 Gibbs samples.

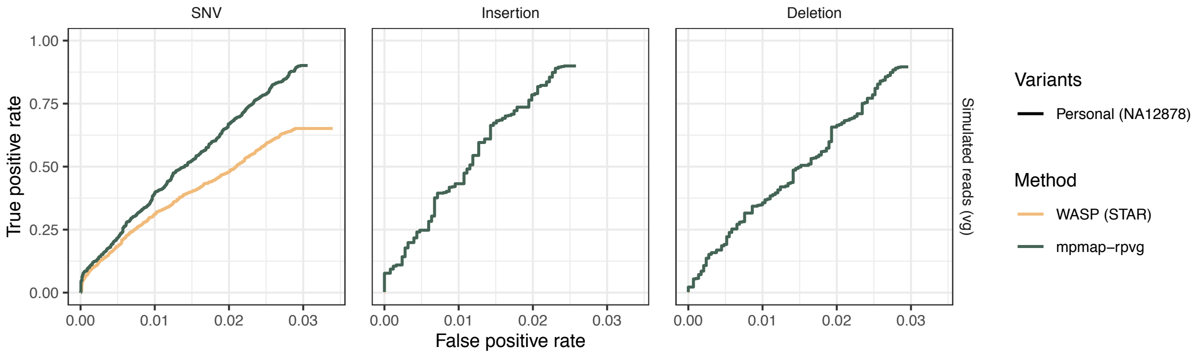


**Supplementary Figure 22: Allele-specific expression benchmark using RNA-seq data from NA12878**
Allele-specific expression (ASE) results comparing the MPMAP-RPVG pipeline against WASP (with STAR as the aligner) using simulated data ("vg sim (ENC, RSEM)" in Supplementary Table 5). Shows true positive rate and false positive rate of ASE significance for different thresholds of variant read count in the simulated data. Variants were defined as showing significant ASE using a two-sided binomial test of the allele-specific read counts with p-values adjusted using the Benjamini-Hochberg procedure and a False Discovery Rate (FDR) $\alpha = 0.1$. All heterozygotic NA12878 variants from the 1000 Genomes Project (1000GP) with at least one read in the simulated data were used for the benchmark. For the MPMAP-RPVG pipeline, we used the personal transcriptome generated from the 1000GP NA12878 haplotypes (Supplementary Table 3). WASP was provided the 1000GP NA12878 haplotypes as input. Note, we only used WASP for bias correction and allele-specific read counting, and not its downstream inference method.

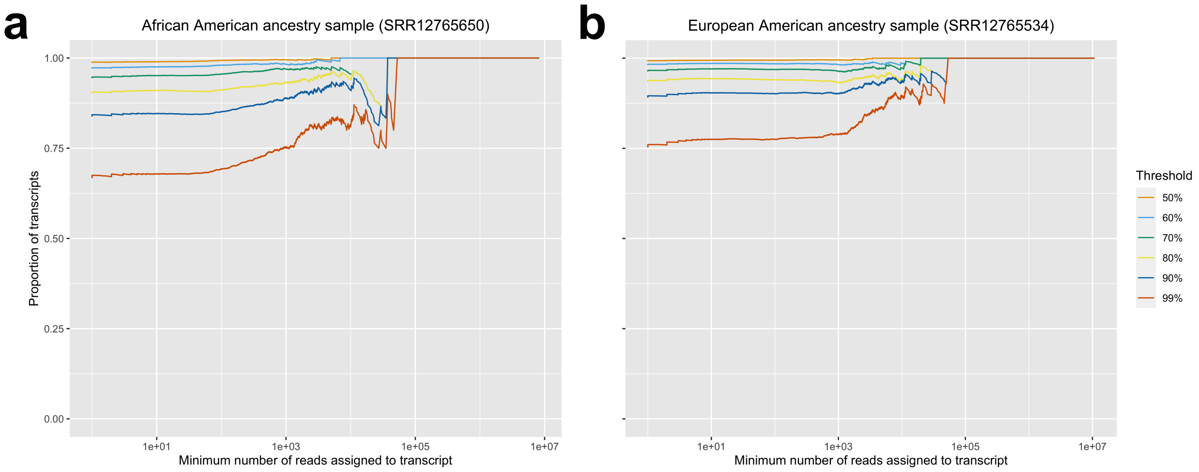


**Supplementary Figure 23: Proportion of marginal expression attributed to $\leq 2$ HSTs of a transcript**

For two samples from **a** an African American individual and **b** a European American individual, the proportion of transcripts for which the marginal expression has at least $X$ proportion assigned to $\leq 2$ HSTs, for various values of $X$ ("SRR12765650" and "SRR12765534" in Supplementary Table 4). Colors correspond to different thresholds on the proportion of marginal expression. A pantranscriptome generated from all 1000 Genomes Project haplotypes were used for the evaluation ("Whole" in Supplementary Table 3). Transcripts with fewer than 1 inferred read are omitted.

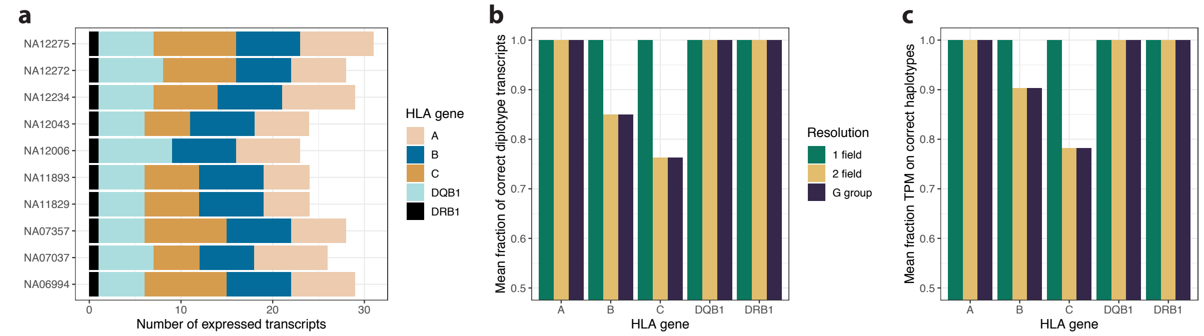


**Supplementary Figure 24: HLA typing evaluation using RNA-seq data from ten Geuvadis samples**

HLA typing results from the MPMAP-RPVG pipeline using real Illumina data from ten CEU Geuvadis samples (see Supplementary Table 4). The evaluation used the "HLA (main)" pantranscriptome containing five of the main HLA genes (Supplementary Table 3). **a** Number of transcripts predicted to be expressed for each sample and gene. **b** Mean fraction of transcripts with correctly predicted diplotypes across samples for 3 different levels of HLA allele resolution. 1 field: Identical allele group, 2 field: Identical protein sequence, G group: Identical nucleotide sequence across antigen recognition site exons. A haplotype-specific transcript (HST) is classified as correct if its corresponding HLA allele matches the prediction based on genomic sequencing data in one of two HLA studies [1, 2] **c** Mean fraction of transcript expression assigned to correct HLA alleles across samples for 3 different levels of HLA allele resolution. Expression is measured in transcripts per million (TPM).

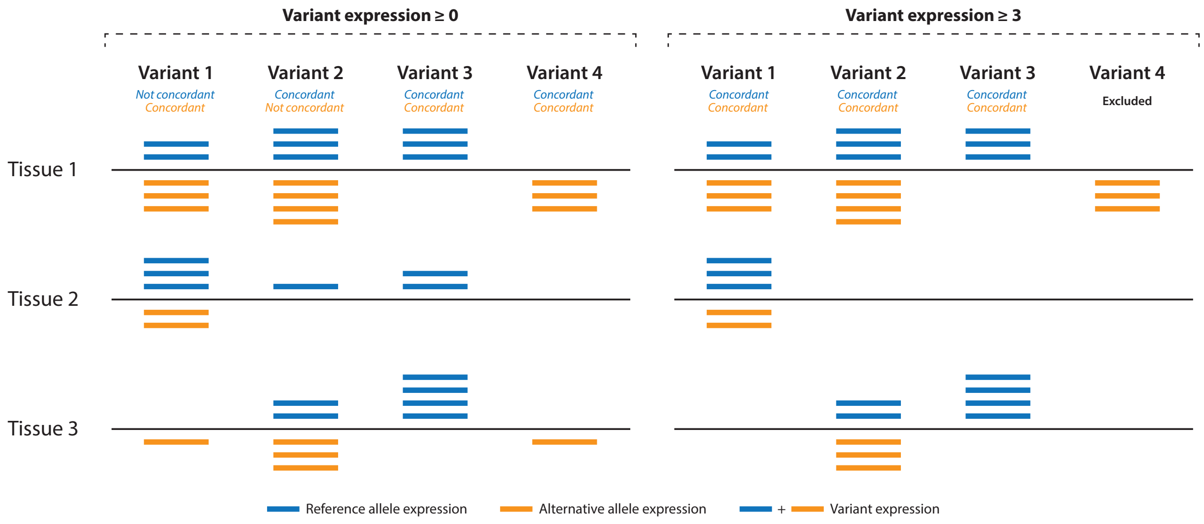


**Supplementary Figure 25: Examples of allele expression concordance across tissues**
A set of examples showing allele concordance across tissues using two different variant expression thresholds. Only three tissues are used in the example for simplicity. Blue and orange bars correspond to reference and alternative allele expression, respectively. Variant expression is calculated as the sum of the two alleles. An allele is defined as concordant if it is either expressed or not expressed in all tissues for which the corresponding variant is expressed. Using this definition all alternative alleles except for the allele in variant 2 are defined as concordant when the minimum variant expression threshold is set to 0. If the variant expression threshold is increased to 3, the alternative allele in variant 2 becomes concordant since tissue 2 will be filtered for this variant. Moreover, variant 4 will be excluded due to tissue 3 being filtered since at least two expressed tissues are needed to compute concordance.

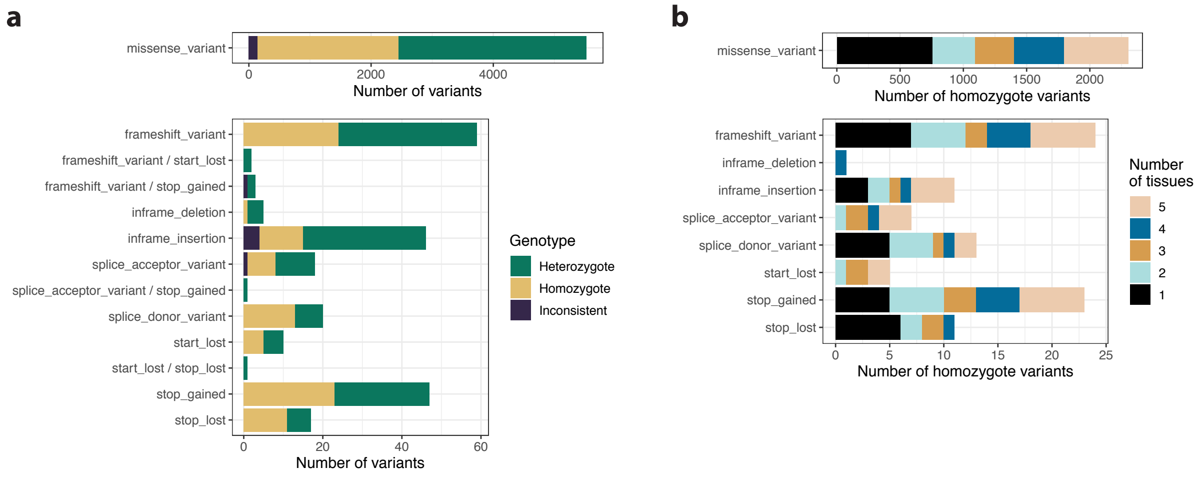


**Supplementary Figure 26: Effect prediction of expressed variants genotyped using RNA-seq data from five tissues**

Prediction of effects on functional elements for expressed variants genotyped with the MPMAP-RPVG pipeline using real Illumina data from five different tissues from the same individual (see Supplementary Table 4). The pipeline used a pantranscriptome generated from all 1000 Genomes Project (1000GP) haplotypes ("Whole" in Supplementary Table 3). The Ensembl Variant Effect Predictor (VEP) toolset [3] was used to predict functional consequences of all variants in exons with a TPM of at least five. **a** Number of expressed variants predicted to have an effect on functional elements for different Sequence Ontology (SO) consequence terms. The colors represent whether a variant is genotyped as heterozygote or homozygote across the tissues. Inconsistent is when the tissues do not agree. Note the large difference in numbers between "missense_variant" (top bar) and the other consequences. **b** Number of predicted homozygote variants for different SO consequence terms. The colors represent the number of tissues the variant is predicted to be expressed in.

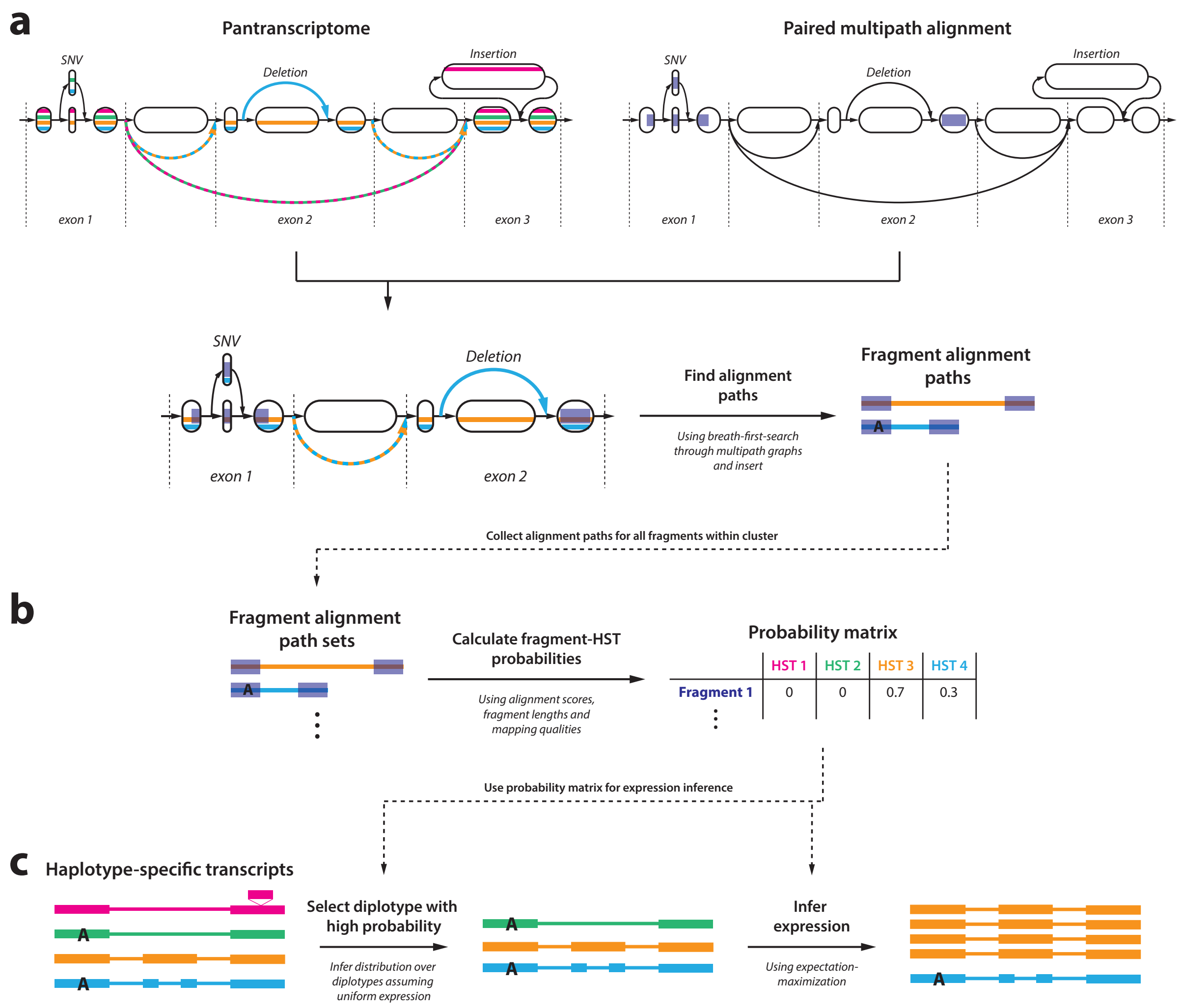


**Supplementary Figure 27: Diagram of haplotype-specific transcript quantification in rpvg** Diagram showing an overview of how RPVG infers expression of haplotype-specific transcripts (HSTs) in a pantranscriptome from a set of paired-end multipath alignments (see Supplementary Figure 1). The colored thin lines correspond to HST paths, and the blue transparent regions correspond to aligned read sequences. In the diagram we assume the sample is diploid, but the algorithm generalize to any ploidy. **a** For each fragment, all paths through the multipath alignment graphs are identified using a depth-first-search (DFS). For paired-end reads, the DFS also traverses the fragment insert creating alignment paths of the whole fragment. Only alignment paths that follow a HST path in the pantranscriptome are kept. **b** The probabilities that each fragment originated from each of the HSTs in a cluster are calculated using the score and length of the fragment alignment paths, and the mapping quality. **c** The fragment-HST probability matrix is used to infer the expression of the HSTs using a nested inference scheme. First, a distribution over diplotypes is inferred. Next, the most probable diplotypes are selected from this distribution and expression is inferred conditioned on the haplotypes using expectation-maximization. The inferred expression values are combined across the most probable diplotypes to produce the final HST expression estimates.

### Supplementary Tables

| Name | Description | Non-exonic variant filter | Number of samples | Number of exonic variants | Number of total variants |
|---|---|---|---|---|---|
| 1000GP (NA12878) | 1000GP variants for sample NA12878 | None | 1 | 152,968 | 4,266,678 |
| 1000GP (EUR, excl. CEU) | 1000GP variants for all samples with European (EUR) ancestry excluding the CEU population | AF $\geq$ 0.002 | 404 | 986,167 | 15,041,731 |
| 1000GP (all, excl. CEU) | 1000GP variants for all samples excluding the CEU population | AF $\geq$ 0.001 | 2,405 | 3,808,242 | 31,731,676 |
| 1000GP (all) | 1000GP variants for all samples | AF $\geq$ 0.001 | 2,504 | 3,873,100 | 29,970,512 |

**Supplementary Table 1: Genomic variant (haplotype) sets**
1000GP: 1000 Genomes Project

| Name | Transcript annotation | Number of transcripts | Variant set† |
|---|---|---|---|
| 1000GP (NA12878) | GENCODE v29 (full-length) | 172,449 | 1000GP (NA12878) |
| 1000GP (EUR, excl. CEU) | GENCODE v29 (full-length) | 172,449 | 1000GP (EUR, excl. CEU) |
| 1000GP (all, excl. CEU) | GENCODE v29 (full-length) | 172,449 | 1000GP (all, excl. CEU) |
| 1000GP (80%, all, excl. CEU) | 80% of GENCODE v29 (full-length) | 137,959 | 1000GP (all, excl. CEU) |
| 1000GP (all) | GENCODE v29 (full-length) | 172,449 | 1000GP (all) |

**Supplementary Table 2: Spliced pangenome graphs**
1000GP: 1000 Genomes Project
†See Supplementary Table 1 for more details on the 1000GP variant sets

| Name | Transcript annotation | Number of transcripts | Haplotype set† | Number of haplotypes | Number of haplotype-specific transcripts |
|---|---|---|---|---|---|
| Personal (NA12878) | GENCODE v29 (full-length) | 172,449 | 1000GP (NA12878) | 2 | 235,400 |
| Europe (excl. CEU) | GENCODE v29 (full-length) | 172,449 | 1000GP (EUR, excl. CEU) | 808 | 2,515,408 |
| Whole (excl. CEU) | GENCODE v29 (full-length) | 172,449 | 1000GP (all, excl. CEU) | 4,810 | 11,626,948 |
| Whole | GENCODE v29 (full-length) | 172,449 | 1000GP (all) | 5,008 | 11,835,580 |
| HLA (main) | GENCODE v29 (full-length) | 44 | IMGT/HLA (A, B, C, DQB1 & DRB1) | 21,386 (genes) | 122,637 |
| HLA (10) | GENCODE v29 (full-length) | 64 | IMGT/HLA (A, B, C, E, DPA1, DPB1, DQA1, DQB1, DRB1 & DRB5) | 23,330 (genes) | 129,031 |

**Supplementary Table 3: Pantranscriptomes**
†See Supplementary Table 1 for more details on the 1000 Genomes Project (1000GP) haplotype sets

| Name / Accession (source) | Reference | Replicate | Cell-line / Tissue | Sequencing type | Number of read-pairs or alignments |
|---|---|---|---|---|---|
| SRR1153470 (SRA) | [4] | NA | NA12878 | Strand-specific PE Illumina HiSeq 2000 | 115,359,773 read-pairs |
| ENCSR000AED (ENCODE) | [5, 6] | 1 | NA12878 | Strand-specific PE Illumina HiSeq 2000 | 97,548,052 read-pairs |
| ENCSR706ANY (ENCODE) | [5, 6] | 1-4 | NA12878 | PacBio Iso-Seq | 2,687,717 alignments |
| CHM13 (T2T†) | [7] | 1 | CHM13 | Strand-specific PE Illumina NovaSeq | 90,930,105 read-pairs |
| ENCSR146ZKR, ENCSR825GWD, ENCSR686JJB, ENCSR502OTI & ENCSR995BHD (ENCODE) | [5, 6] | NA | Adrenal gland, Sigmoid colon, Adipose tissue, Psoas muscle & Aorta | Strand-specific PE Illumina HiSeq 2000 | 109,405,824 - 251,137,848 read-pairs |
| NA07051, NA11832, NA11840, NA11930, NA12287, NA12775 & NA12889 (Geuvadis‡) | [8] | NA | NA07051, NA11832, NA11840, NA11930, NA12287, NA12775 & NA12889 | PE Illumina HiSeq 2000 | 30,371,135 - 62,803,070 read-pairs |
| NA06994, NA07037, NA07357, NA11829, NA11893, NA12006, NA12043, NA12234, NA12272 & NA12275 (Geuvadis‡) | [8] | NA | NA06994, NA07037, NA07357, NA11829, NA11893, NA12006, NA12043, NA12234, NA12272 & NA12275 | PE Illumina HiSeq 2000 | 33,436,056 - 56,561,356 read-pairs |
| ERR1050073, ERR1050074 & ERR1050075 (SRA) | [9] | NA | NA19238, NA19239 & NA19240 | Strand-specific PE Illumina HiSeq 2500 | 59,219,085 - 72,457,249 read-pairs |
| ERR1050076, ERR1050077 & ERR1050078 (SRA) | [9] | NA | HG00512, HG00513 & HG00514 | Strand-specific PE Illumina HiSeq 2500 | 58,601,893 - 69,073,722 read-pairs |
| ERR1050079, ERR1050080 & ERR1050081 (SRA) | [9] | NA | HG00731, HG00732 & HG00733 | Strand-specific PE Illumina HiSeq 2500 | 56,254,714 - 92,075,712 read-pairs |
| SRR12765650 & SRR12765534 (SRA) | [10] | NA | Rectum & Ileum | Strand-specific PE Illumina HiSeq 4000 | 18,117,125 - 29,360,643 read-pairs |

**Supplementary Table 4: Read sets and alignments**
PE: Paired-end
†Downloaded from the T2T consortium data repository:
`https://github.com/nanopore-wgs-consortium/CHM13`
‡Geuvadis data available at:
`https://www.internationalgenome.org/data-portal/data-collection/geuvadis`

| **Name** | **Template reads†** | **Haplotype-specific transcripts‡** | **Expression values** | **Simulation parameters** | **Number of read-pairs** |
|---|---|---|---|---|---|
| vg sim (SRR, uniform) | SRR1153470 | Personal (NA12878) | Uniform | Indel error: 0.001, mean: 277, sd: 43 | 50,000,000 |
| vg sim (ENC, uniform) | ENCSR000AED | Personal (NA12878) | Uniform | Indel error: 0.001, mean: 216, sd: 24 | 50,000,000 |
| RSEM (ENC, uniform) | ENCSR000AED | Personal (NA12878) | Uniform | theta0: 0 | 50,000,000 |
| vg sim (SRR, RSEM) | SRR1153470 | Personal (NA12878) | RSEM estimates | Indel error: 0.001, mean: 277, sd: 43 | 50,000,000 |
| vg sim (ENC, RSEM) | ENCSR000AED | Personal (NA12878) | RSEM estimates | Indel error: 0.001, mean: 216, sd: 24 | 50,000,000 |

**Supplementary Table 5: Simulated read sets**
†See Supplementary Table 4 for more details on the read sets used to fit the simulation model
‡See Supplementary Table 3 for more details on the haplotype-specific transcript sets

| Software tool(s) or library | Used for step involved in | Version(s) or GitHub commit(s)† |
|---|---|---|
| bcftools | Filtering, subsetting, normalizing and annotating variants. | v1.9 & v1.11 |
| samtools | Filtering, sorting, converting and indexing alignments. | v1.9 |
| bedtools | Converting alignments to regions and calculating coverage. | v2.29.1 |
| seqtk | Subsampling reads. | v1.3 |
| HISAT2 | Indexing graphs and mapping reads. | v2.2.0 & v2.2.1 |
| STAR | Indexing references and mapping reads. | v2.7.9a |
| WASP | Bias correction and read counting. | v0.3.4 |
| Bowtie2 | Indexing pantranscriptomes and mapping reads. | v2.3.5.1 & v2.4.4 |
| RSEM | Indexing pantranscriptomes and inferring expression. | v1.3.1 & v1.3.3 |
| Kallisto | Indexing pantranscriptomes and inferring expression. | v0.46.1 & v0.46.2 |
| Salmon | Indexing pantranscriptomes and inferring expression. | v1.2.1 & v1.5.2 |
| rpvg | Inferring expression. | 1d91a9e3 |
| VEP | Predicting variant effects | v103.1 |
| vg construct, vg convert, vg rna, vg ids, vg index, vg stats & vg gbwt | Constructing graphs and pantranscriptomes. | v1.23.0, c861e23e, 8ff022c3 & c4bbd63b |
| vg gbwt | Constructing GBWT r-index. | 883f0f87 & c4bbd63b |
| vg snarls, vg prune & vg index | Constructing distance and GCSA graph index. | 8ff022c3 & c4bbd63b |
| vg mpmap, vg surject, vg stats & vg augment | Augmenting graph with HLA haplotypes. | c4bbd63b |
| vg paths & vg surject | Creating reference transcript alignments for mapping benchmark. | c861e23e |
| vg view & vg sim | Simulating reads from transcript paths. | 765d2215 |
| vg map & vg mpmap | Mapping reads to graph. | 2cea1e25 |
| vg inject, vg view, vg paths & vg surject | Converting alignments between graph (GAM) and reference (BAM). | 385fd636 & 2cea1e25 |
| vg stats, vg view & vg gampcompare | Comparing graph alignments for mapping benchmark. | 096bfdce |
| vgrna-project-scripts | Converting expression profiles for simulations. Inferring coverage, overlap and bias statistics for mapping benchmark. Comparing haplotype-specific transcript (HST) sequences for expression benchmark. Converting path counts and HST expression to allele expression. Calculating credibility intervals. Adding reference flanks and introns to HLA alleles. Analysing imprinting signatures. Plotting the results. | 71442ea4, 94176204, 34d563ba, 7eb08153, 93bc0a90, 2220bb08, f9db749e & 39f7d9b9 |
| SeqLib | Parsing alignments and calculating overlaps. | 08771285 |
| hlaseqlib | Imputing HLA alleles. | v0.0.2 |

**Supplementary Table 6: Versions of software used**
†Different subcommands in the vg toolkit and parts of the pipeline stabilized at different times during our development process, hence the variety of commits used.

### Supplementary Notes

#### vg mpmap algorithm details

##### Seeding

Algorithm 1 contains pseudocode for the algorithm used to extract supermaximal exact match (SMEM) seeds. This algorithm utilizes a GCSA2 index to query suffix array intervals, which can be located in the graph similarly to the FM-index [11, 12]. After finding a longest maximal exact match (MEM), the algorithm navigates upward in the implicit suffix tree using a longest common prefix (LCP) array. After computing the SMEMs of a read, we find the minimally-more-frequent MEMs for each SMEM, subject to a minimum length. This algorithm for detecting them (Algorithm 3) proceeds by querying the number of occurrences of a "probe substring" with a size equal to the minimum length (Algorithm 2). If the number of occurrences equal to the number of occurrences of the SMEM, then no minimally-more-frequent MEM that meets the minimum length criterion can contain that probe substring. Alternatively, if the substring occurs more frequently than the SMEM, then it must be part of a minimally-more-frequent MEM. The endpoint of this MEM is found using a bisection search. A benefit of the design of this algorithm is that, if the minimum length is longer than a match of a random sequence is likely to be, it is possible to skip over many characters in the SMEM without ever querying them.

##### Reachability between seeds

Algorithm 4 presents a simplified version of the algorithm for computing reachability between exact match seeds. The simplification lies in that seed positions are treated as synonymous with nodes in the graph. In reality, seeds correspond to a path through the graph, not single nodes. Also, collinear seeds sometimes overlap each other either on the read or in the graph. For instance, this occurs when there are indel errors in a homopolymer, in which case the MEMs on either side of the error will both try to match the entire homopolymer. The full algorithm in VG MPMAP handles these nuances, but the details are cumbersome to present. The three stages of the reachability algorithm achieve linear run time in the typical case in different ways. In stage 1 (Algorithm 5), most transitive reachability relationships are not discovered, so the number of edges is usually linear. In stage 2 (Algorithm 6), the quadratic factor is limited to the number of noncollinear seeds that can nonetheless reach each other in the pangenome graph. In stage 3 (Algorithm 7), an early stopping condition limits the quadratic factor to the number of seeds that have more than one collinear successor.

##### Dynamic programming with multiple traceback

The pseudocode in Algorithm 8 presents a simplified version of the multiple traceback algorithm in which only the two highest-scoring alignments are returned. The logic extends naturally to finding the top $k$ alignments, but the details are somewhat complicated and not particularly enlightening. The pseudocode also presents the multiple-traceback algorithm for a Needleman-Wunsch alignment for the sake of simplicity, but the generalization to POA is straightforward. The insight underlying this algorithm is that the second highest-scoring traceback diverges from the highest-scoring one at the cell where the difference is minimized between the score that is entered in the dynamic programming matrix and the other scores that could have been entered there. Moreover, the score of this alignment differs from the highest-scoring alignment's score by exactly that difference.

**Algorithm 1:** Stage 1 of MEM finding

**Input:** GCSA2 index $G$, LCP array $L$, read $R$

**Output:** SMEMs between $R$ and sequence graph

```
Function FindSMEMs(G, L, R):
    b ← |R|, e ← |R|                          // ends of a match's read interval
    s ← G.fullSAInterval()                    // suffix array interval on the graph
    while b > 0 do
        // extend match by one character
        ŝ ← G.LF(s, R[b − 1])
        if ŝ.empty() then
            // the SMEM is exhausted
            yield (s, b, e)
            n ← L.parent(s)                   // match's parent suffix tree node
            e ← b + n.longestCommonPrefix()
            s ← n.interval()
        else
            // the match was successful
            b ← b − 1
            s ← ŝ
        end
    end
    yield (s, b, e)
```

**Algorithm 2:** Check if a substring is more frequent than an SMEM (Subroutine of Stage 2 of MEM-finding)

**Input:** GCSA2 index $G$, read $R$, SMEM $(s, b, e)$, probe substring $(pb, pe)$ with $pb \geq b$ and $pe \leq e$

**Output:** MEM $(t, p, pe)$, where $p \geq pb$ is the minimum index such that $(p, pe)$ occurs more times than $(b, e)$ in the graph.

```
Function MoreFrequentProbe(G, R, (s, b, e), (pb, pe)):
    p ← pe
    t ← G.fullSAInterval()
    while p > pb do
        t̂ ← G.LF(t, R[p − 1])
        if G.count(t̂) > G.count(s) then
            p ← p − 1
            t ← t̂
        else
            break
        end
    end
    return (t, p, pe)
```

**Algorithm 3:** Stage 2 of MEM-finding

**Input:** GCSA2 index $G$, read $R$, SMEM $(s, b, e)$, minimum length $L$

**Output:** All minimally-more-frequent MEMs

**Function** `MoreFrequentMEMs(`$G$, $R$, $(s, b, e)$, $L$`):`
- $mb \leftarrow b$, $me \leftarrow b + L$ `// the probe substring`
- **while** $me \leq e$ **do**
  - $t, pb, pe \leftarrow$ `MoreFrequentProbe(`$G$, $R$, $(s, b, e)$, $(mb, me)$`)`
  - **if** $pb \neq mb$ **then**
    - `// probe cannot occur in any minimally-more-frequent MEM`
    - $mb \leftarrow pb + 1$
    - $me \leftarrow mb + L$
  - **else**
    - `// probe occurs in minimally-more-frequent MEM, bisect to find end`
    - $l \leftarrow me$, $h \leftarrow e$
    - **while** $h \neq l$ **do**
      - $c \leftarrow \lfloor (h + l)/2 \rfloor$
      - $t, pb, pe \leftarrow$ `MoreFrequentProbe(`$G$, $R$, $(s, b, e)$, $(mb, c)$`)`
      - **if** $pb \neq mb$ **then**
        - $h \leftarrow c$
      - **else**
        - $l \leftarrow c$
      - **end**
    - **end**
    - **yield** $(t, mb, h)$
    - $me \leftarrow h + 1$
    - $mb \leftarrow me - L$
  - **end**
- **end**

**Algorithm 4:** Construct the connectivity graph between seeds

**Input:** $D$ a DAG, $S = \{(n, b, e)\}$ the MEM seeds (represented by a node $n$ and a read interval $[b, e)$)

**Output:** Transitive reduction of graph with seeds as nodes and edges between the seeds if they are connected by a path in $D$ and the read intervals are collinear.

**Function** `SeedGraph(`$G$, $S$`):`
- `/* make an initial graph ignoring collinearity on the read */`
- $G_0 \leftarrow$ `TentativeSeedGraph(`$D, S$`)`
- `/* rewire edges to respect collinearity */`
- $G_1 \leftarrow$ `CollinearSeedGraph(`$G_0$`)`
- `/* remove transitive edges (if any) */`
- $G_2 \leftarrow$ `TransitiveReduction (`$G_1$`)`
- **return** $G_2$

**Algorithm 5:** Stage 1 of constructing connectivity graph between seeds

**Input:** $D$ a DAG, $S = \{(n, b, e)\}$, the set of MEM seeds represented by a node $n$ from $D$, and a read interval $[b, e)$

**Output:** Graph with seeds for nodes and edges between two seeds whenever there is a walk in the graph that does not include any other seed

```
Function TentativeSeedGraph(G, S):
    G_0.nodes() ← S                                  // nodes correspond to seeds
    foreach n in D.topologicalOrder() do
        if ∃ some seed s = (n, b, e) then
            // seeds that can reach n can reach s
            foreach t in n.predecessorSeeds() do
                G_0.addEdge(t, s)
            end
            // s blocks earlier seeds and reaches n's successors
            foreach m in n.successors() do
                m.predecessorSeeds().insert(s)
            end
        else
            // all seeds that reach n can reach its successors
            foreach m in n.successors() do
                foreach s in n.predecessorSeeds() do
                    m.predecessorSeeds().insert(s)
                end
            end
        end
    end
    return G_0
```

**Algorithm 6:** Stage 2 of constructing connectivity graph between seeds

**Input:** $G_0$, graph with seeds as nodes and edges indicating that there is a walk connecting the two seeds in the graph (may exclude transitive edges)

**Output:** Graph with seeds for nodes and edges indicating 1) that there is a walk connecting the two seeds, and 2) the seeds are collinear on the read. Transitive edges may be excluded.

```
Function CollinearSeedGraph(G_0):
    G_1.nodes() ← G_0.nodes()
    foreach s in G_0.topologicalOrder() do
        // initialize a queue with s's predecessors
        q.init()
        foreach t in s.predecessors() do
            q.enqueue(t)
        end
        while q is not empty do
            t = q.dequeue()
            // t is only in q if it can reach s, check for collinearity
            if t.readInterval() is collinear with s.readInterval() then
                G_1.addEdge(t, s)
                // don't explore t's collinear predecessors, they will only produce
                   transitive edges
            else
                s.noncollinearPredecessors().insert(t)
                // must explore t's collinear predecessors
                foreach u in t.predecessors() do
                    q.enqueue(u)
                end
            end
            // always explore t's noncollinear predecessors
            foreach u in t.noncollinearPredecessors() do
                q.enqueue(u)
            end
        end
    end
    return G_1
```

**Algorithm 7:** Stage 3 of constructing connectivity graph between seeds

```
Input: G1, a DAG
Output: The transitive reduction of G1
Function TransitiveReduction (G1):
    G2.nodes() ← G1.nodes()
    foreach s in G1.topologicalOrder() do
        if s.numSuccessors() = 1 then
            // all walks out of s use the edge, it cannot be transitive
            continue
        end
        // keep track of which seeds have been visited
        v ← ∅
        // iterate over neighbors in topological order
        foreach t in s.neighbors() do
            if t ∈ v then
                // t is reachable from an earlier edge, (s, t) is transitive
                G2.removeEdge(s, t)
            else
                // mark all nodes reachable from this edge as visited
                v.insert(t)
                foreach u ∈ t.reachableByDFS() do
                    v.insert(u)
                end
            end
        end
    end
```

**Algorithm 8:** Find the two highest-scoring alignments from a single DP matrix

**Input:** $M$ the dynamic programming matrix of an alignment, $g$ gap penalty, $S$ score matrix, $Q_1$ and $Q_2$ the sequences being aligned

**Output:** The two top-scoring alignments

```
Function MultipleTraceback(M, g, S, Q1, Q2):
    i = M.numRows(), j = M.numCols()
    s ← M[i, j]                                         // the alignment score
    a1 ← ∅, a2 ← ∅                                      // alignments to trace
    d ← ∅                                  // point of deflection for 2nd alignment
    Δ ← s                                  // score difference for 2nd alignment
    while i ≠ 0 or j ≠ 0 do
        if M[i, j] = M[i − 1, j − 1] + S[Q1[i − 1], Q2[j − 1]] then
            i′, j′ ← i − 1, j − 1
        else if M[i, j] = M[i − 1, j] − g then
            i′, j′ ← i − 1, j
        else
            i′, j′ ← i, j − 1
        end
        a1.prepend(i′, j′)
        // look at suboptimal extensions to find next-best traceback
        if i′, j′ ≠ i − 1, j − 1 and M[i, j] − (M[i − 1, j − 1] + S[Q1[i − 1], Q2[j − 1]]) < Δ then
            Δ ← M[i, j] − (M[i − 1, j − 1] + S[Q1[i − 1], Q2[j − 1]])
            d ← (i, j ⟶ i − 1, j − 1)
        end
        if i′, j′ ≠ i − 1, j and M[i, j] − (M[i − 1, j] − g) < Δ then
            Δ ← M[i, j] − (M[i − 1, j] − g)
            d ← (i, j ⟶ i − 1, j)
        end
        if i′, j′ ≠ i, j − 1 and M[i, j] − (M[i, j − 1] − g) < Δ then
            Δ ← M[i, j] − (M[i, j − 1] − g)
            d ← (i, j ⟶ i, j − 1)
        end
        i, j ← i′, j′
    end
    // the 2nd best traceback
    i = M.numRows(), j = M.numCols()
    while i ≠ 0 or j ≠ 0 do
        if d.from() = i, j then
            // this is where 2nd best traceback differs from optimal
            i′, j′ ← d.to()
        else if M[i, j] = M[i − 1, j − 1] + S[Q1[i − 1], Q2[j − 1]] then
            i′, j′ ← i − 1, j − 1
        else if M[i, j] = M[i − 1, j] − g then
            i′, j′ ← i − 1, j
        else
            i′, j′ ← i, j − 1
        end
        a2.prepend(i′, j′)
        i, j ← i′, j′
    end
    return (a1, s), (a2, s − Δ)
```
